# Cas9-enriched nanopore sequencing enables comprehensive and multiplexed detection of repeat expansions

**DOI:** 10.1101/2025.11.24.690106

**Authors:** Seungbok Lee, Chanju Jung, Minjeong Kim, Narae Kim, Gue-Ho Hwang, Soon-Tae Lee, Kon Chu, Sang Kun Lee, Han-Joon Kim, Jong-Hee Chae, Sangsu Bae, Jangsup Moon

## Abstract

Short tandem repeat (STR) expansion is a major genetic mechanism underlying numerous neurogenetic disorders. However, traditional PCR amplification and short-read next-generation sequencing (NGS)-based methods often fail to detect complex and large expansions including methylation information. Here, we modified an amplification-free nanopore Cas9-targeted sequencing (nCATS) platform and developed a dedicated analysis algorithm, STRiker, for the simultaneous assessment of all currently defined STR loci (56 sites) using a single test with genomic DNA from patient-derived blood cells. We ultimately identified pathogenic repeat expansions in 12 of 37 patients (32.4%) with cerebellar ataxia who remained genetically undiagnosed despite extensive prior genetic testing, in *FGF14* (n=4), *ATXN8OS*, *NOP56*, *RFC1* (n=2 each), and *PRNP* and *NOTCH2NLC* (n=1 each). Additionally, family-based cascade screening revealed six relatives with repeat expansions in five families. These results demonstrated a broader diversity of pathogenic repeat structures, particularly in *FGF14*, and illustrated that CpG methylation can mitigate the pathogenic effects of repeat expansions. This approach offers a powerful strategy for improving the diagnosis of STR-related neurogenetic diseases, for cerebellar ataxia and other STR-related diseases in the future.

**One Sentence Summary:** Improved nanopore Cas9-targeted sequencing with STRiker enhances diagnosis of STR- related neurogenetic diseases and offers broader molecular insights.

## Introduction

Short tandem repeats (STRs) are repetitive sequences of short motifs, typically 2–7 base pairs. More than one million STR loci have been reported in the human genome (*1*), and STR expansions at specific sites can produce numerous genetic disorders (*2*). The STR expansion loci associated with causing diseases are constantly being updated; as of the start of this study (*i*.*e*., late 2023), approximately 56 different loci have been identified to cause Mendelian diseases, particularly neurological and neuromuscular (*2*). For example, spinocerebellar ataxias (SCAs) are a group of genetically heterogeneous, progressive, and neurodegenerative diseases, many of which are caused by STR expansions in specific loci (*3*).

However, detecting and diagnosing the disease-causing STR expansions in patients remains challenging. Traditionally, fragment analysis methods based on PCR amplification and short-read next-generation sequencing (NGS) platforms have been employed for diagnosing STR-related disorders (*4*). Although existing methods are useful for detecting repetitive expansion occurrences, they still have several important limitations. For example, PCR-based fragment analyses lack sequence information, limiting the assessment of interruptions and motif structures. Additionally, targeted amplicon sequencing approaches using PCR are susceptible to polymerase errors inherent to repeat motif characteristics (*5*). Conventional NGS methods, which typically generate read lengths of 150–300 bp, cannot accurately determine repeat structures when the pathogenic STR expansion (*i*.*e*., motif size × repeat count) exceeds this sequencing length limitation. Whole-genome sequencing-based approaches also fail to capture accurate repeat information, as the assembly of repetitive sequences remains highly challenging, leading to a loss of precise repeat count data. Furthermore, DNA methylation information is crucial for determining whether a disease is actually occurring; however, DNA methylation can be lost during the PCR amplification process.

To overcome these limitations, a PCR amplification-free nanopore Cas9-targeted sequencing (nCATS) method has been previously developed (*6*). This nCATS approach also adopted CRISPR–Cas9 to enrich specific genomic regions for long-read nanopore sequencing, enabling the detection of small variants, STR expansions, and epigenetic modifications, such as methylation. Moreover, nCATS has been employed to identify multiple STR loci associated with genetic disorders simultaneously (*6–10*). Furthermore, nCATS has facilitated the identification of novel repeat expansion motifs and interruption patterns within known ataxia loci, such as *FGF14* (*11, 12*).

Thus, this study aimed to improve the nCATS method to ensure that a single test can simultaneously diagnose all existing STR expansion loci. This study employed the following strategies: i) we selected optimal guide RNAs (gRNAs) with the best performance at each gene and optimized CRISPR RNA (crRNA):trans-activating crRNA (tracrRNA) ratios for *in vitro* cleavage assay; ii) we designed each gRNA to generate approximately 10 kb fragments to minimize the existing length bias in nanopore sequencing; iii) we developed a fast, dedicated program to detect *de novo* STR contexts and interruption patterns. Building on the optimized nCATS method, we tested the revised method in 37 patients with cerebellar ataxia who had undergone extensive genetic testing but remained genetically undiagnosed. Subsequently, we newly diagnosed 12 of these 37 patients (32.4%); the process of DNA extraction to diagnosis was achieved rapidly, within 25 hours. We identified several *de novo* repeat contexts during the methodological process, particularly in *FGF14*, and methylation changes during transmission in *NOTCH2NLC*. Collectively, our optimized nCATS method can provide a fast and precise strategy for enhancing the diagnosis of STR-related neurogenetic diseases such as cerebellar ataxia.

## Results

### Optimization of gRNAs to maximize *in vitro* cleavage efficiency

First, we conducted *in vitro* experiments to improve the nCATS method and evaluate all existing STR regions in a single test (**Fig. 1A**). We grouped 56 STR regions for which an association with genetic disorders is either known or suggested into two panels (**fig. S1**): Panel A specifically included STR loci associated with cerebellar ataxia; whereas Panel B encompassed all other loci implicated in STR-related genetic disorders. Since the potential to read relatively shorter DNA fragments more abundantly in nanopore sequencing has previously been reported (*5, 13*), we designed gRNAs to generate approximately 10 kb DNA fragments for nCATS, including the region of interest (ROI), to minimize the length bias (**Fig. 1A**). We conducted *in vitro* cleavage assays with three or five gRNA candidates upstream and downstream of each ROI (**Fig. 1B**). In this experiment, we used single-guide RNA (sgRNA) formation constructed by linking dual RNA components of crRNA and tracrRNA. Ultimately, we selected a pair of gRNAs that showed the best DNA cleavage activities upstream and downstream of each ROI **(Fig. 1C** and **fig. S2**). The gRNAs chosen for the final panel exhibited remarkably higher cleavage efficiencies, with most exceeding 90%, compared to the other gRNA candidates (**Fig. 1D**).

**Fig. 1.**
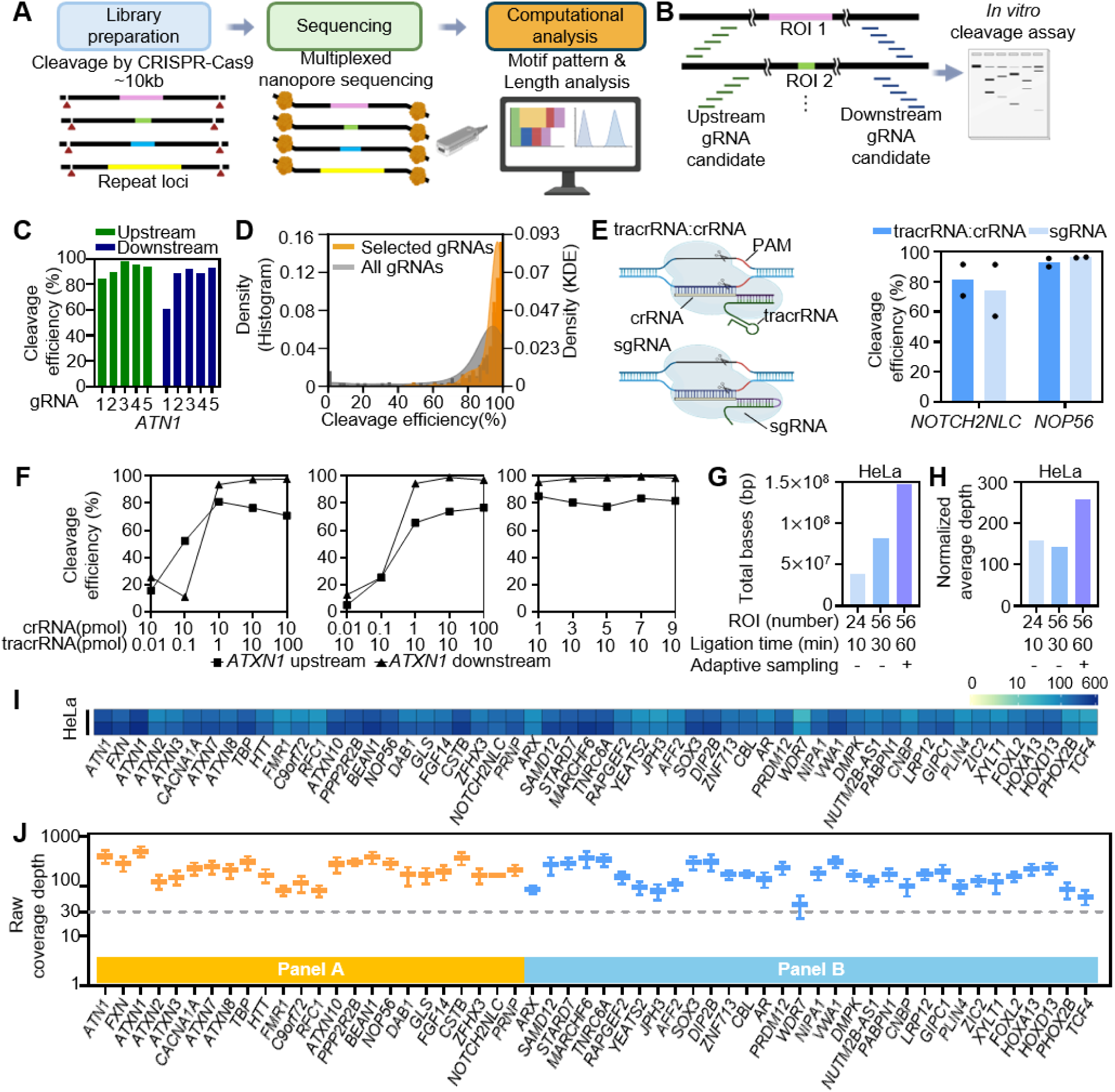
Optimization of the nCATS workflow for multiplex short tandem repeat sequencing. (A) Overview of the nanopore Cas9-targeted sequencing (nCATS) pipeline. Approximately 10 kb DNA fragments surrounding short tandem repeat (STR) regions were excised by CRISPR–Cas9 and underwent Oxford nanopore sequencing. Computational analysis allowed visualization of repeat motif patterns and read length distributions. Created with BioRender.com. (B) Schematic of guide RNA (gRNA) design and *in vitro* cleavage (IVC) assay. Three or five gRNA candidates were tested on each side of the region of interest (ROI) to identify the most efficient pair. Created with BioRender.com. (C) IVC results for upstream and downstream gRNAs targeting the *ATN1* locus. Cleavage efficiency is shown for each guide. (D) Histogram and kernel density estimate (KDE) of a comparison between cleavage efficiencies of selected gRNAs (orange) and all tested gRNAs (gray); most selected gRNAs exhibited >90% cleavage efficiency. (E) Comparison of cleavage efficiency when Cas9 was guided by either an sgRNA or a tracrRNA:crRNA duplex; similar activities are observed for both RNA formats. Created with BioRender.com. (F) Cleavage efficiencies were measured under varying crRNA and tracrRNA concentrations, with the other component held constant. (G–H) Effect of adapter ligation time and adaptive sampling (AS) on sequencing yield and coverage. Extension of ligation time and use of AS improved total bases and normalized depth in HeLa cells. (I) Heatmap of the per-locus sequencing coverage across 56 targeted STR loci in HeLa cells, demonstrating balanced coverage across all targets. (J) Raw coverage depth for each target locus is summarized across all sequencing runs in HeLa cells, indicating uniform read depth across all genes included in Panels A and B. Panel A (orange) includes genes associated with cerebellar ataxia, and Panel B (blue) includes additional STR-related genes.

Next, we compared the cleavage efficiency of Cas9 guided by sgRNA versus tracrRNA:crRNA. The sgRNA construct has been commonly utilized due to its simple structure formation compared to the tracrRNA:crRNA (*14–19*). However, we hypothesized that the Cas9/tracrRNA:crRNA construct might exhibit similar cleavage efficiency to Cas9/sgRNA *in vitro* and might also have advantages in terms of multiplexing. Expectedly, we observed that Cas9/tracrRNA:crRNA yielded comparable cleavage efficiency over Cas9/sgRNA (**Fig. 1E**). Given that sgRNA is approximately 112 nucleotides long, while crRNA and tracrRNA are about 36 and 67 nucleotides in length, respectively, we hypothesized that the Cas9/tracrRNA:crRNA construct would have advantages in terms of multiplexing and cost. Additionally, we performed experiments to determine the optimal combination ratio of crRNA and tracrRNA *in vitro*. Thus, one RNA concentration was fixed at 10 pmol, while the other was titrated from 0.01 to 100 pmol. Saturation of cleavage activity was observed at concentrations above 1 pmol in both conditions (**Fig. 1F**). Based on this, we used 6 pmol tracrRNA and 3 pmol crRNA per ROI in all subsequent experiments.

### Optimization of the nCATS protocol to ensure uniform coverage across STR regions

With the selected 56 gRNA pairs, we next improved the nCATS protocol to enhance the overall coverage depth and ensure more uniform coverage across the STR target panel. Using genomic DNA (gDNA) extracted from HeLa cells, we optimized the nCATS protocol through three adjustments: i) modifying the dA-tailing step as separate steps, ii) extending the adapter ligation time to enhance the ligation efficiency, and iii) applying nanopore adaptive sampling to ensure maximum enrichment of the ROI regions (**fig. S3**). Typically, Cas9 cleavage and dA-tailing were performed together. However, we speculated that any residual Cas9 following cleavage might interfere with the dA-tailing reaction; therefore, we separated this process into two steps: Cas9 was first removed via bead purification, and then dA-tailing was performed. An extension of the ligation step from 10 to 60 minutes resulted in a higher number of total bases (*i*.*e*., the sum of “coverage depth × ROI length” per condition); meanwhile, application of adaptive sampling further improved the overall coverage depth across STR loci (**Fig. 1G**). Ultimately, we could obtain length-normalized average coverage of 258 across all 56 STR loci, indicating that our optimized nCATS method can be applied to diagnose all known STR-related diseases (**Fig. 1H**).

To evaluate the coverage uniformity for all target loci, we created heatmaps to visualize the per-locus read depth. When we applied the optimized nCATS workflow to HeLa samples with the full panel comprising 56 STR regions (*i*.*e*., Panel A + Panel B), evenly distributed coverage was observed across most targets, as indicated by the relatively uniform color patterns across rows, which present loci per individual **(Fig. 1I)**. Quantitative analysis further showed that raw coverage depth was consistent among loci across the entire panel **(Fig. 1J)**. These results demonstrate that the optimized nCATS method achieved robust and uniform per-locus coverage across all 56 targets, confirming the reliability and scalability of the modified method for multiplexed STR analysis.

### Application of the optimized nCATS method for patient-derived blood cells

Building on the optimized nCATS workflow, we next applied this modified method for gDNA from patient-derived blood cells (**Fig. 2A**). Unlike gDNA from HeLa cells, which is abundantly available in the laboratory, gDNA from patient-derived blood cells is often limited in quantity. Therefore, we attempted to use minimal amounts of gDNA from patient-derived blood cells (*i*.*e*., 5 µg) in contrast to the previously applied 10 µg of gDNA from HeLa cells that was used to optimize the nCATS protocol. As part of the proof-of-concept, we tested 15 patient-derived gDNA samples with 24 STR regions (*i*.*e*., Panel A) and 7 samples with the full panel of 56 STR regions (*i*.*e*., Panel A + Panel B).

**Fig. 2.**
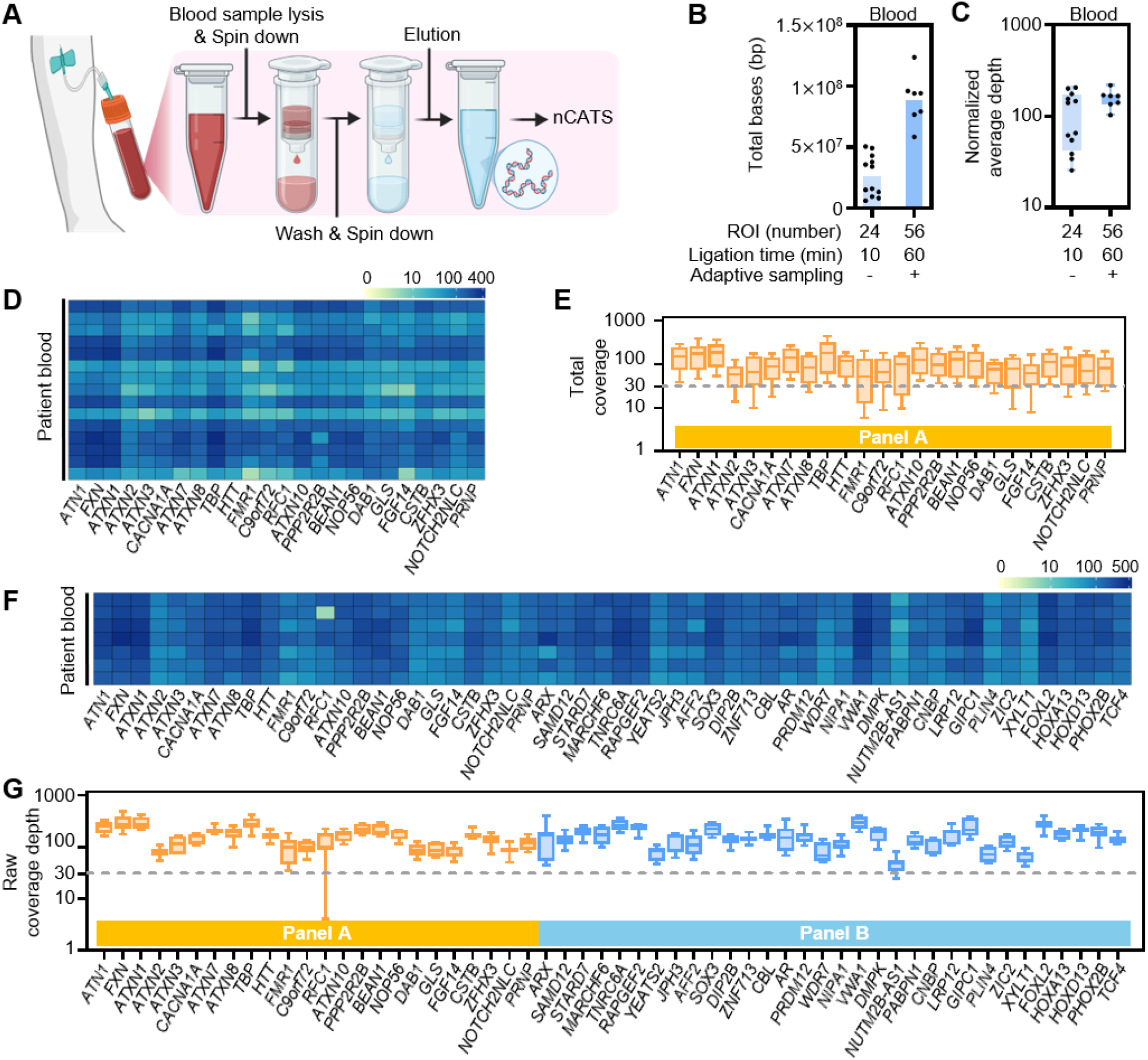
Application of the optimized nCATS workflow to patient blood samples. (A) Workflow for gDNA extraction from blood samples of patients. After lysis and elution, nCATS was performed. Created with BioRender.com. (B–C) Effect of ligation time (10 min vs. 60 min) and adaptive sampling on total bases (B) and normalized sequencing depth (C). (D–G) Sequencing coverage across blood-derived samples. Heatmaps show per-locus sequencing depth, where color intensity represents read depth per locus and demonstrates consistent coverage distribution. Boxplots with a summary of the total coverage per STR locus; a consistent distribution of coverage was observed. (D, E) Results from the 24-locus panel (Panel A). (F, G) Results from the 56-locus panel (Panel A + Panel B).

Consistent with the results from HeLa cells, extending the ligation time from 10 to 60 minutes and applying adaptive sampling likewise improved total bases, thereby increasing overall coverage despite the increased number of ROIs (**Fig. 2B**). Using patient-derived gDNA samples, the length-normalized average depth reached 108 across 24 loci and 156 across all 56 loci (**Fig. 2C**). Although the total number of base reads obtained from patient-derived gDNA samples was slightly lower than that obtained from HeLa cells, the sequencing depth was sufficient to diagnose the STR-related genetic diseases.

To confirm whether the uniform coverage achieved in HeLa cells could be reproduced in clinical samples, heatmaps were again created to visualize the per-locus read depth of patient-derived gDNA. When we treated samples with 24 STR regions (*i*.*e*., Panel A), the coverage appeared evenly distributed across most targets, as indicated by the relatively uniform color patterns across rows (*i*.*e*., loci per individual) (**Fig. 2, D** and **E**). When we further treated samples with a full panel of 56 STR regions (*i*.*e*., Panel A + Panel B), the coverage similarly appeared evenly distributed across most targets (**Fig. 2, F** and **G**). Thus, these findings validate that the uniform coverage established during HeLa-based optimization is stably reproducible in patient-derived samples, supporting the robustness and clinical scalability of the optimized nCATS method and confirming its usability for comprehensive STR analysis.

### STRiker-based analysis of STR expansion in 56 different loci

After reading the STR regions, determining their sequence context and size remained challenging. Short-read-based programs, such as ExpansionHunter and GangSTR, rely on previously reported motifs (*20, 21*), whereas long-read-based tools, including Straglr and HMMSTR, enable more accurate detection of repeat expansions (*22, 23*). Thus, this study developed (i) a dedicated analysis algorithm, named STRiker (https://github.com/BaeLab/STRiker), for detecting STR expansions from nanopore sequencing data, and (ii) a user-friendly pipeline that identifies novel repeat motifs, detects interruption patterns, and automatically generates diagnostic reports for streamlined interpretation **(Fig. 3A)**. STRiker is capable of identifying both reference and *de novo* motifs directly from sequencing reads. In STRiker, any sequence units observed in tandem in the reference or input reads ≥3 times are automatically defined as a candidate STR motif. To account for rotational symmetry, motifs that could be transformed into each other through rotation were recognized as a single motif. This enables the discovery of previously unreported motifs while maintaining consistency with known pathogenic variants.

**Fig. 3.**
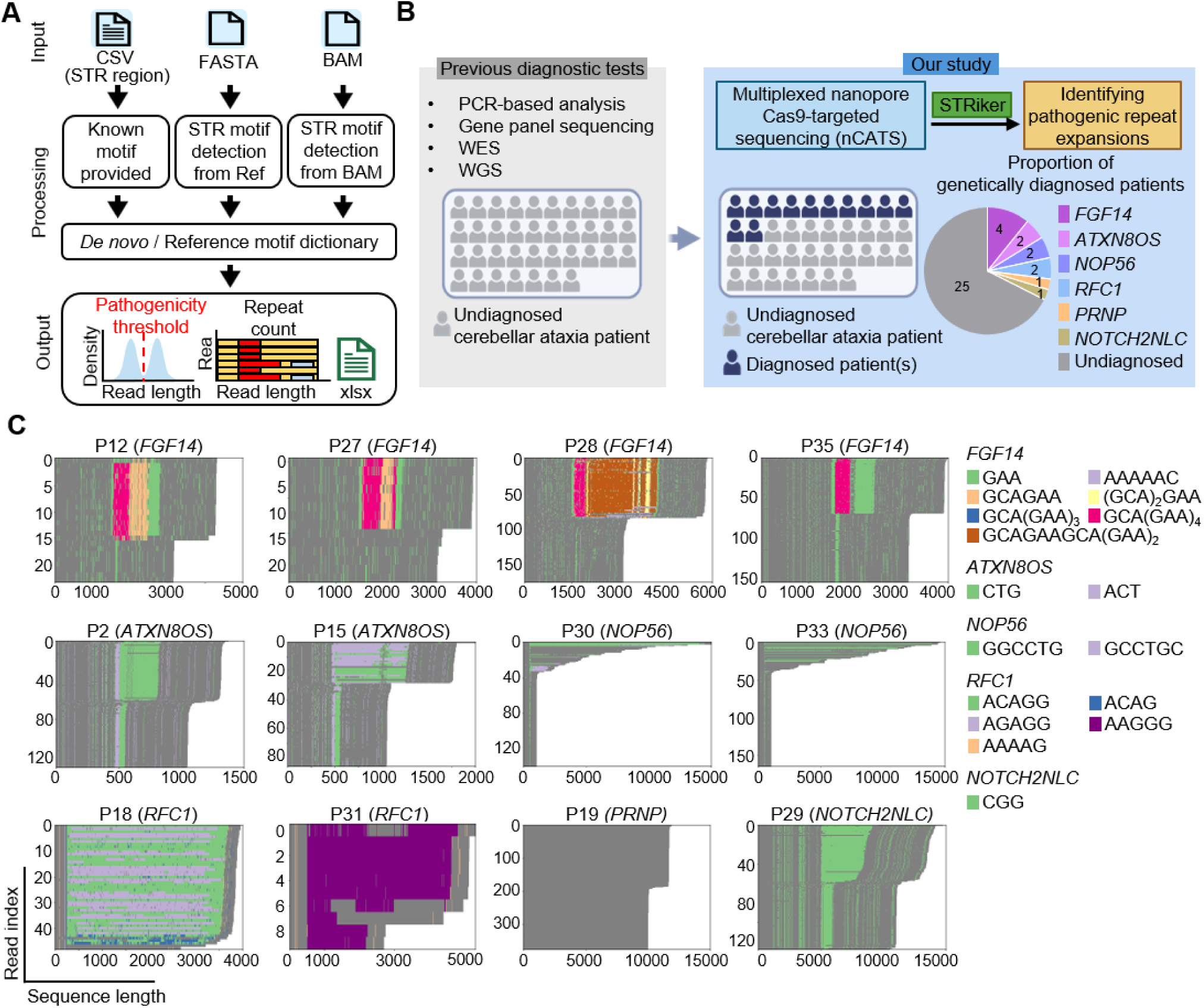
STRiker analysis pipeline and representative visualization outputs. (A) Overview of the STRiker pipeline developed for motif detection and repeat quantification from nCATS sequencing data. Using a BAM file and a target locus file as the input, STRiker identified both known (reference) and novel (*de novo*) repeat motifs and generated output files containing a length distribution plot and motif heatmap plot. (B) Diagnostic summary of 37 genetically undiagnosed patients with cerebellar ataxia. A total of 12 patients were newly diagnosed with pathogenic STR expansions, corresponding to a diagnostic yield of 32.4%. Created with BioRender.com. (C) Heatmap of repeat motifs in newly diagnosed patients identified through the STRiker pipeline. Different colors indicate different motifs within the same gene. *PRNP* repeat expansions comprise an octapeptide motif (PHGGGWGQ) of 24 bp and are, therefore, displayed without motif color differentiation. Ref, reference sequence used for motif detection; WGS, whole-genome sequencing; WES, whole-exome sequencing.

The STRiker analysis pipeline integrates three input sources to generate comprehensive STR profiles (**Fig. 3A**). The tool accepts STR region coordinates, along with optional known pathogenic motifs (CSV format), a reference genome (FASTA format), and aligned sequencing data (BAM format) as input. STRiker performs motif detection from both the reference genome and sequencing reads, building a unified reference/*de novo* motif count report file. STRiker generates two primary outputs: (i) visual representations, including repeat count distributions that clearly delineate normal from pathogenic expansions, and heatmap visualizations showing motif patterns across individual reads; (ii) an Excel file (xlsx format) containing detailed motif composition data with both reference/*de novo* motifs, and their respective repeat counts.

In the output file (PDF format), each gene-specific figure includes a heatmap illustrating the distribution of repeat motifs within individual reads, where distinct colors represent different motif types, and a read length density plot displaying Gaussian-fitted peaks corresponding to normal and expanded alleles. The red vertical line indicates the threshold for pathogenic expansion, allowing rapid visual distinction between normal and disease-associated alleles.

### Diagnostic utility in patients with cerebellar ataxia

Building on the improved experimental workflow and analysis algorithm, we ultimately applied the optimized nCATS methods to a total of 37 patients with cerebellar ataxia who were genetically undiagnosed (**table S1**). Since the patients exhibited clinically clear symptoms of cerebellar ataxia, we first applied Panel A to 30 patients, and then we used the full Panel for seven additional patients. It should be emphasized that most of the patients (35 out of 37) had previously undergone extensive genetic studies, including targeted PCR-based fragment analysis (89.2%), gene panel sequencing (75.7%), whole-exome sequencing (62.2%), and whole-genome sequencing (18.9%).

Notably, we identified 12 patients with STR expansions in *FGF14* (4 patients), *ATXN8OS*, *RFC1*, and *NOP56* (2 patients each), as well as *NOTCH2NLC* and *PRNP* (1 patient each). The overall diagnostic rate was 32.4% (12 out of 37 patients) (**Fig. 3, B and C, and table S2**). This allowed the patients to complete their long diagnostic odyssey, with an average time from symptom onset to genetic diagnosis of 8.9 years (range, 1–32 years; median, 5.5 years).

Representative examples of STRiker-generated visualizations are presented for each genetically diagnosed patient, including per-read sequence alignments that depict the structure and motif composition of the expanded alleles, as well as density plots summarizing overall read length distributions (**Fig. 3C** and **fig. S4**). These graphical outputs allow intuitive examination of allele structures, repeat length distributions, and sequence interruptions, providing information that was previously difficult to visualize using conventional methods. Such visualization-based reporting enables clinicians and researchers to readily recognize the configuration and interruption patterns of pathogenic alleles, thereby improving interpretability and facilitating clinical application of nCATS-based STR analysis.

Following the genetic diagnosis of each patient, we performed cascade screening in six families to further test relatives of patients who have already been diagnosed by our improved nCATS method. Further results showed that four affected relatives were also diagnosed (two with *FGF14* and one each with *ATXN8OS* and *NOP56*) (**fig. S5**). In addition, we could identify two asymptomatic (without clinical symptoms) or presymptomatic (currently symptom-free but expected to develop symptoms later) carriers with repeat expansions in *FGF14* and *PRNP*, respectively. One carrier with the *FGF14* expansion was the father of Patient 28. While both the patient and her sister exhibited dystonia and gait disturbance in early childhood, the carrier father remained neurologically asymptomatic even in his 50s. The presymptomatic carrier was the cousin of Patient 19, who possessed a *PRNP* octapeptide repeat expansion. This carrier is currently in his late 20s and has no significant symptoms. However, since his mother developed symptoms in her 30s and passed away in her 40s, our research team is striving to identify preventive management strategies for this presymptomatic carrier.

### Identification of novel patterns in disease-related STRs

STRiker revealed novel STR patterns in specific ROIs from the sequencing data of 37 patients. Moreover, certain regions were observed to contain more than two distinct expansion motifs. For example, three distinct motifs (ACAGG, AGAGG, AAAAG) were identified for *RFC1* (**Fig. 4A**), which aligns with previous reports (*24–26*). Notably, repeat expansions have been reported with GAA and GAAGGA motifs for *FGF14*, as well as a (GAA)nGCA pattern (*12, 27–29*). Furthermore, we identified additional novel expansions, such as AAAAAC, and a complex GCAGAAGCA(GAA)2 repeat motif in one of our pedigrees (Patient 28) (**Fig. 4B**). Interestingly, the affected siblings with novel *FGF14* repeat motifs exhibited distinctive clinical features, including childhood onset, dystonia, spasticity, and T2 signal abnormalities at globus pallidus, which are rarely observed in typical cerebellar ataxia (**table S2**). These findings demonstrate the broader diversity of pathogenic repeat structures in *FGF14*-related disorders than previously recognized. The presence of novel repeat motifs at other loci also warrants confirmation, emphasizing the need for further investigation into their potential clinical and biological significance.

**Fig. 4.**
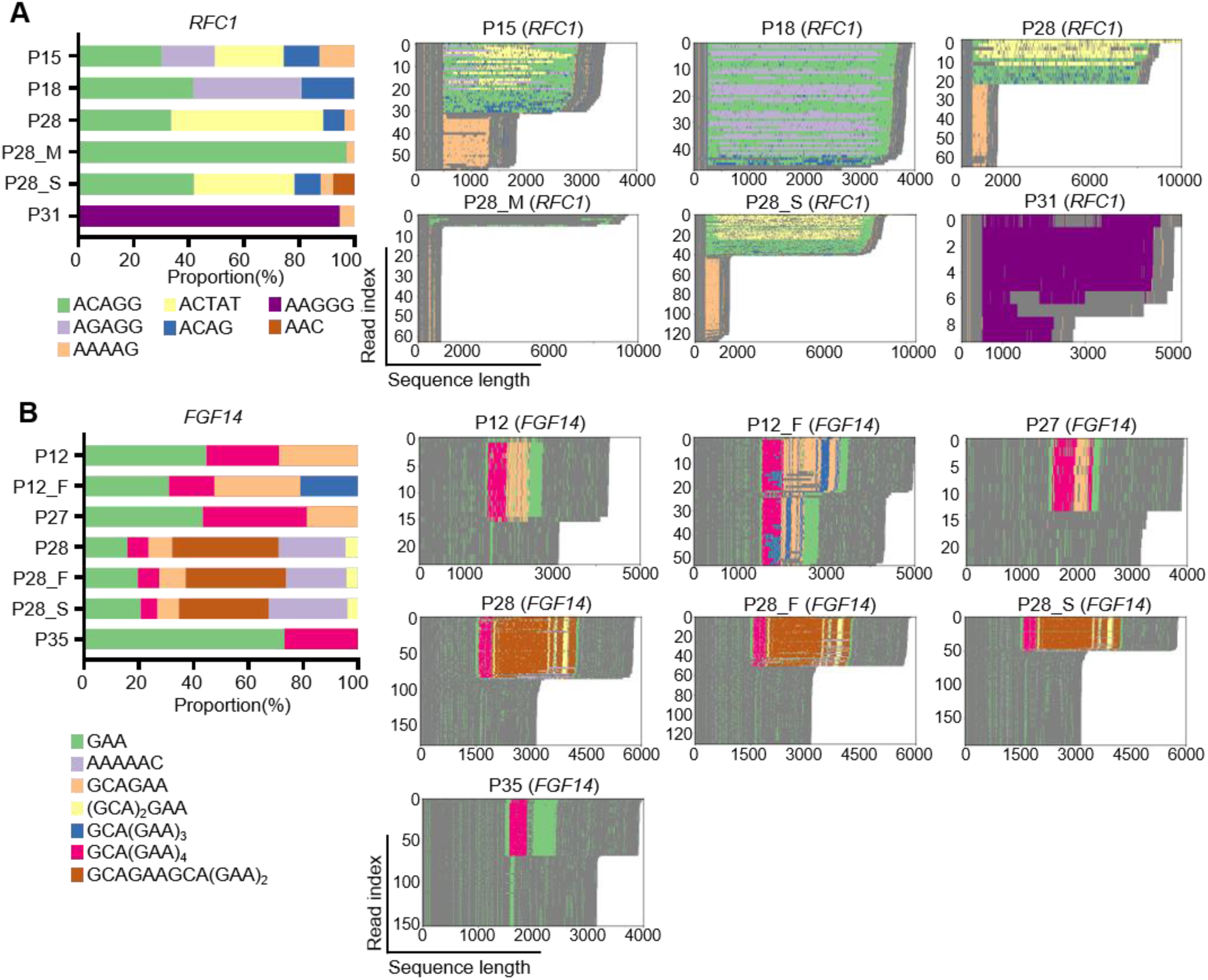
Analysis of STR motif composition and length distribution in *RFC1* and *FGF14* expansion carriers. (A–B) Stacked bar plots of the proportion of each repeat motif identified from nanopore sequencing reads, with distinct colors indicating different motif types. Heatmaps of the individual sequencing reads aligned to the repeat region, visualizing the arrangement and overall length of repeat units for each allele. (A) *RFC1* expansions analyzed by STRiker revealed three distinct repeat motifs that were consistent with previously reported pathogenic configurations: ACAGG, AGAGG, and AAAAG. These results demonstrate that the STRiker pipeline reliably detects known pathogenic repeat structures. (B) *FGF14* expansions displayed variable combinations of GAA-based motifs, including previously reported GAA, GAAGGA, and (GAA)nGCA configurations, as well as newly identified patterns, such as the AAAAAC expansion and a GCAGAAGCA(GAA)2 repeat motif. These findings illustrate that STRiker identified both canonical and novel repeat structures. Notably, multiplexed nCATS sequencing enabled the simultaneous detection of repeat expansions from different genes in a single diagnostic test, as in P28, carrying heterozygous expansions in RFC1 (carrier) and FGF14 (pathogenic).

### Intergenerational dynamics of repeat expansion and CpG methylation

Furthermore, we investigated the DNA methylation information, which is known to be crucial in determining whether a disease is actually occurring. The preservation of native DNA modifications by the optimized nCATS allowed us to extract CpG methylation information from the same reads. Although the multiplexed nCATS enables methylation profiling across all targeted loci, we specifically focused on the *NOTCH2NLC* locus, which is associated with neuronal intranuclear inclusion disease (NIID) **(Fig. 5A)** (*30, 31*), given our previous findings on this disorder **(fig. S6)** (*32*). We then analyzed methylation profiles of individuals in association with repeat expansions at this locus.

**Fig. 5.**
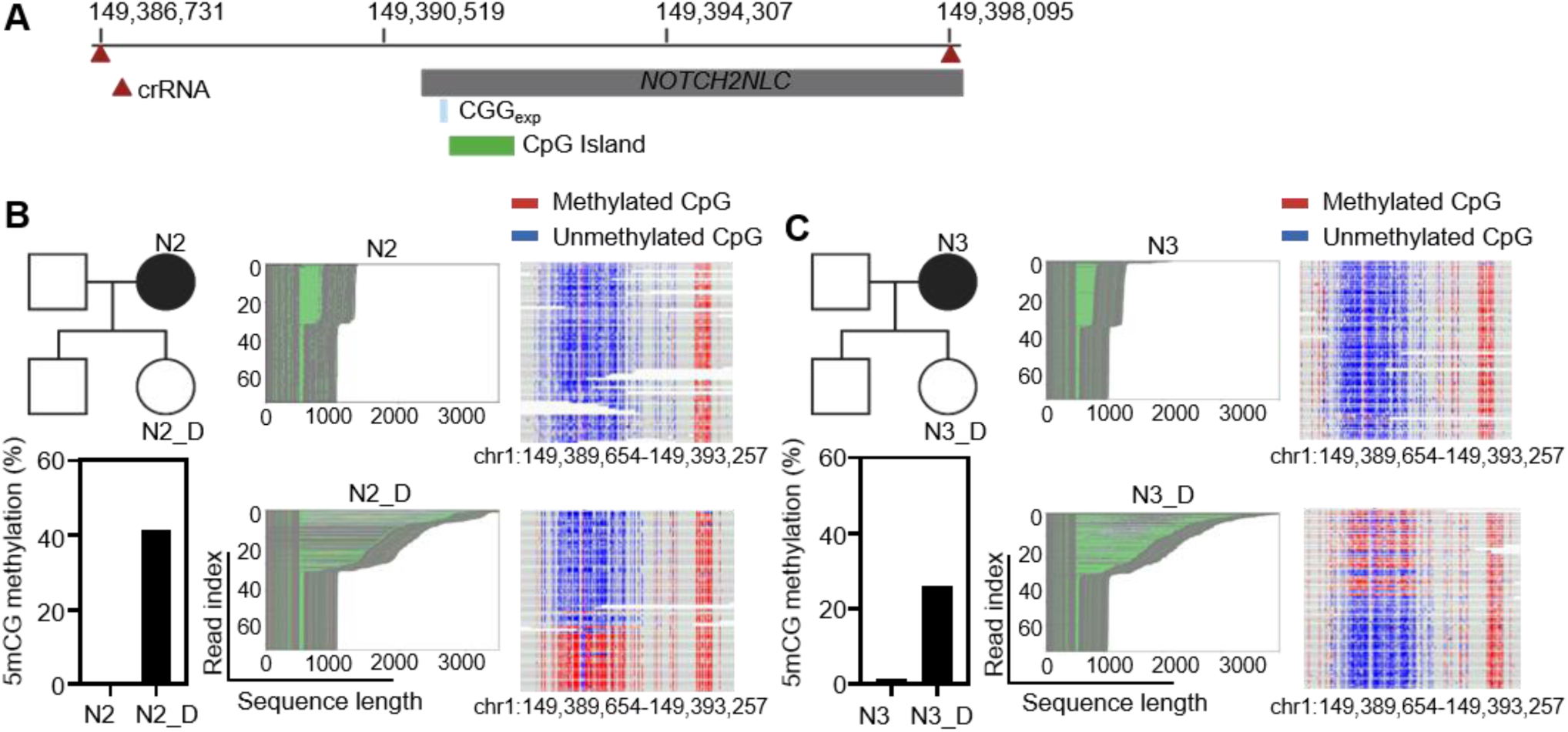
Intergenerational interplay between repeat expansions and CpG methylation at the *NOTCH2NLC* locus. (A) Schematic of the *NOTCH2NLC* locus with the expanded CGG repeat region (CGGexp), upstream CpG island, and targeted crRNA sites. CpG methylation states were obtained directly from native nCATS reads. Created with BioRender.com. (B, C) Two independent families with the expansion during maternal inheritance. In both cases, the mother (N) carried an unmethylated expanded allele and was affected, while the daughter (N_D) inherited a longer repeat with extensive methylation and remained asymptomatic. In each case, repeat structures are visualized on the left, and CpG methylation states (methylated: red; unmethylated: blue) are shown on the right. Corresponding 5mCG methylation levels are summarized in the bar plots below each pedigree. Created with BioRender.com.

Previous studies have described several cases of repeat contraction during paternal transmission, in which the fathers carried longer GGC repeat expansions but remained asymptomatic due to hypermethylation. However, the offspring of these fathers inherited alleles that were contracted relative to the paternal expansion yet still exhibited pathologically expanded traits, and developed NIID—likely as a result of demethylation and loss of epigenetic suppression (*30–34*). In our previous study, we also identified an NIID family with the same pattern, in which both children were affected, whereas the father, who had hyperexpansion, remained asymptomatic **(fig. S6)**. However, we observed a distinct pattern in our cohort, with expansion occurring during maternal transmission. In two unrelated families, the mothers carried expanded alleles without CpG methylation and were affected, whereas the offspring inherited even longer expansions but remained asymptomatic until the last follow-up, at which the offspring were aged in their 40s–50s, and extensive methylation was also observed across the repeat-flanking region (**Fig. 5B** and **5C**).

Together, these findings demonstrate a close association between the repeat expansion length and the methylation state, which may be important for predicting disease manifestation. Particularly, these findings also suggest that repeat contraction or expansion across generations may vary depending on whether the transmission is paternal or maternal.

### Summary of the optimized nCATS–STRiker workflow

Our optimized nCATS method integrates streamlined library preparation, multiplexed Cas9-targeted nanopore sequencing, and STRiker analysis to enable rapid and comprehensive detection of STR expansions (**Fig. 6**). The total time from patient-derived gDNA extraction to sequencing completion was approximately 25 hours (∼2 hours for DNA extraction, 5 hours for library preparation, and 18 hours for sequencing). The subsequent computational analysis with STRiker was completed within several minutes. This represents a marked reduction in diagnostic time compared with conventional workflows, in which PCR-based fragment analysis and NGS panel testing are performed sequentially or in parallel, often requiring two to six months from sample submission to final reporting in a clinical setting. In contrast, our approach enables a complete STR analysis within one to two days, achieving a substantial improvement in diagnostic efficiency.

**Fig. 6.**
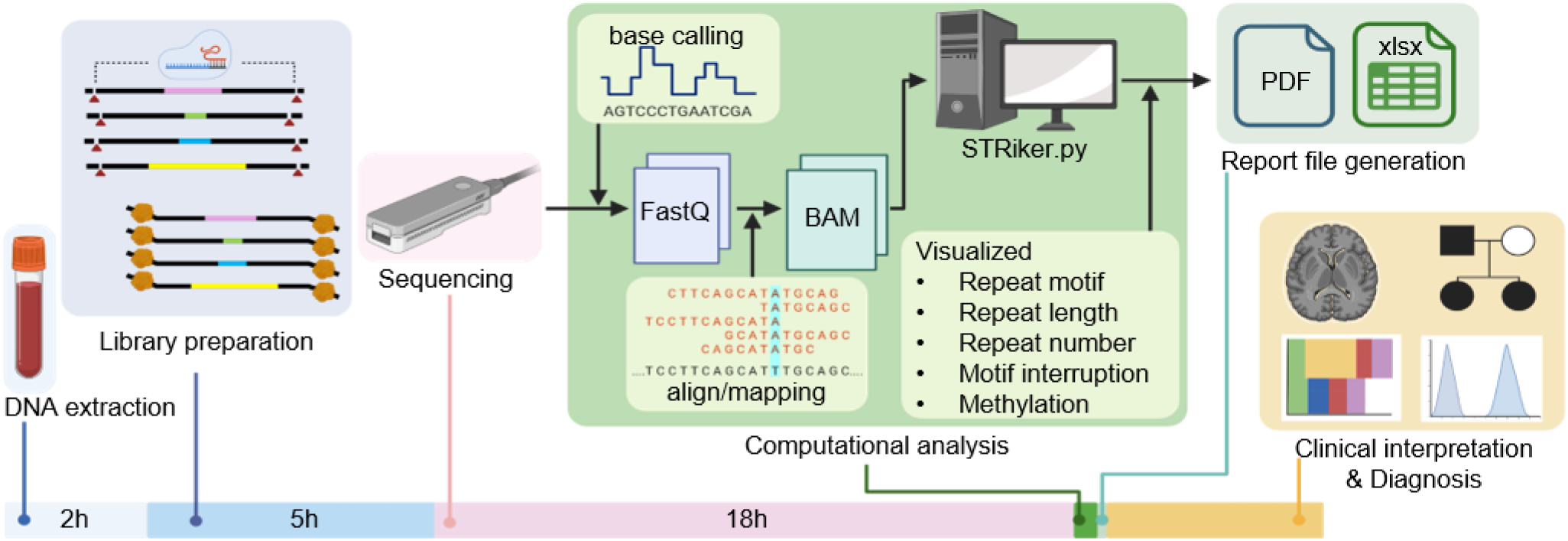
Schematic overview of the nCATS-based diagnostic workflow for STR expansion disorders. Sample processing and sequencing workflow. Genomic DNA extracted from patient blood samples underwent nCATS for 56 STR-associated loci. After multiplexed nanopore sequencing, the raw data were base-called and aligned, and STRiker was used to analyze repeat structures and lengths. The pipeline generates a visualized report (PDF, Excel) to support clinical interpretation. The total turnaround time from DNA extraction to data analysis was approximately 25 hours (∼2 hours for DNA extraction, 5 hours for library preparation, 18 hours for sequencing), with the subsequent computational analysis completed within several minutes. Created with BioRender.com.

## Discussion

This study established an optimized nCATS method to provide a comprehensive analysis of STR expansions, which is based on an amplification-free, long-read nanopore sequencing platform. For this, we selected gRNAs with high activity *in vitro*, adjusted cleavage lengths to approximately 10 kb to mitigate the length bias of nanopore sequencing, and employed adaptive sampling and methylation analysis with nanopore sequencing. We also developed a dedicated algorithm to identify unbiased repeat motifs. Moreover, in the study of complex STR patterns from long-read data, STRiker characterized both reference and *de novo* motifs within repeat expansions. Notably, STRiker can process datasets containing millions of reads within a reasonable timeframe. Ultimately, we were able to perform a simultaneous assessment of all currently defined STR loci (*i*.*e*., 56 sites) using a single test, with the gDNAs from the blood cells of patients.

Cerebellar ataxia is a group of disorders resulting from mutations in various genes, genetic variants, and different inheritance patterns. While some patients can be diagnosed through short-read NGS technologies, a substantial proportion of patients remain undiagnosed, especially in adult-onset cases. Throughout the optimized nCATS method, we newly identified pathogenic repeat expansions in 12 patients (32.4%) among 37 patients with cerebellar ataxia who remained genetically undiagnosed despite extensive prior genetic testing. Notably, *FGF14* expansions were identified in over 10% of the patients (4 out of 37), suggesting that this locus may be underrecognized in clinical settings. Repeat expansions in *ATXN8OS*, *NOP56*, and *RFC1* were also identified in multiple individuals, highlighting that loci often missed by conventional approaches can now be more reliably screened using long-read sequencing. Furthermore, the *PRNP* octapeptide repeat expansion identified in our cohort represents the first reported case in an Asian population. Nonetheless, further studies are warranted to determine the proportion of undiagnosed cases attributable to each of these loci.

In addition, cascade screening of 12 diagnosed families identified family members carrying pathogenic expanded alleles in five families, including two individuals who were asymptomatic or presymptomatic carriers. Such family-based testing enables not only early diagnosis in at-risk relatives but also informed reproductive planning, including the use of preimplantation genetic testing, to allow the birth of unaffected offspring.

Our observations further demonstrate that CpG methylation can mitigate the pathogenic effects of repeat expansions, suggesting that epigenetic protection may act during both contraction and expansion events across generations. In particular, although previous studies reported that pathogenic alleles in the *NOTCH2NLC* gene can arise through contraction following paternal transmission, our study newly identified the opposite pattern in two unrelated pedigrees, in which maternally inherited alleles underwent expansion and the STR region became hypermethylated in the offspring. Meanwhile, whether STR changes differ depending on maternal or paternal inheritance remains to be investigated; however, these findings highlight the importance of simultaneously profiling repeat length and methylation status to accurately assess disease risk in STR-associated disorders, even in familial cases.

However, our study has several potential limitations. First, the pathogenicity of the novel repeat motifs identified in this study could not be fully validated. As more patients are diagnosed using our platform, we anticipate that the clinical relevance of these novel motifs will become clearer. Recent reports have described that some individuals with *FGF14* expansions exhibit alternative phenotypes, such as Parkinson’s disease (*35*), raising the possibility that distinct repeat motifs may contribute to phenotypic variability. Supporting this, patients in our cohort who harbored novel *FGF14* repeat motifs also exhibited atypical clinical presentations. Second, in asymptomatic individuals carrying hypermethylated expanded alleles, although we hypothesized that methylation-mediated suppression occurs, age-dependent penetrance cannot be ruled out. As our diagnostic platform enables the identification of previously undiagnosed repeat expansion disorders, the accumulation of additional cases will help improve our understanding of these mechanisms. Despite these limitations, we believe that this approach can contribute to elucidating the genotypic and phenotypic diversity of repeat expansion disorders, similar to the role that whole-exome sequencing has played in advancing our understanding of many rare genetic diseases.

Collectively, our results demonstrate the clinical utility of long-read, amplification-free STR sequencing in the diagnosis of neurogenetic disorders. The optimized nCATS platform, featuring an intuitive visualization tool, provides a robust and scalable solution for the accurate detection and interpretation of repeat expansions. As new disease-associated STR loci are continually discovered and curated, our platform will be actively updated to incorporate these loci, ensuring that diagnostic coverage remains comprehensive and current. Indeed, the ability of this platform to simultaneously screen multiple STR loci holds promise not only for improving diagnostic outcomes but also for deepening our understanding of disease pathogenesis in cerebellar ataxia and other repeat-associated diseases.

## Materials and Methods

### Study design

This study was designed as a single-center, retrospective observational study combined with methodological development of the nCATS platform and a dedicated analysis pipeline (STRiker) for comprehensive detection of disease-associated short tandem repeat (STR) expansions. The work comprised three components: (i) optimization of the nCATS experimental workflow, including guide RNA selection and library preparation conditions; (ii) development of the STRiker analysis pipeline for automated repeat-length estimation, motif identification, and generation of user-friendly visual reports; (iii) application of the finalized nCATS–STRiker workflow to a diagnostic cohort of 37 patients with cerebellar ataxia, followed by cascade screening of available family members to characterize inheritance patterns and generation-to-generation changes in repeat length and CpG methylation.

The clinical component of the study used retrospectively collected genomic DNA and clinical information from patients and relatives who fulfilled the eligibility criteria described in the “Study participants” section. The optimized nCATS–STRiker workflow was applied to all available samples using either Panel A or the full 56-locus panel, and results were interpreted with reference to established pathogenic repeat-size thresholds, methylation profiles, and individual clinical phenotypes. The sample size was determined by the availability of eligible samples during the study period and was considered sufficient for methodological optimization and estimation of diagnostic yield.

The primary endpoint was the diagnostic yield of the nCATS–STRiker workflow, defined as the proportion of previously undiagnosed patients with cerebellar ataxia in whom a pathogenic or likely pathogenic STR expansion was identified. Secondary objectives included evaluation of on-target sequencing performance (total on-target bases, length-normalized coverage depth, and per-locus coverage uniformity), characterization of known and novel STR motifs and interruption patterns across 56 loci, assessment of intergenerational dynamics of repeat length and CpG methylation (with a particular focus on *NOTCH2NLC*), and benchmarking of STRiker runtime performance on nanopore datasets.

### Study participants

Participants were retrospectively enrolled from Seoul National University Hospital (SNUH) based on the following eligibility criteria: (1) Korean patients who visited the Department of Neurology at SNUH; (2) patients suspected of having cerebellar ataxia, with evidence of cerebellar atrophy on brain imaging or neurological symptoms indicative of cerebellar dysfunction, such as ataxia, dysarthria, and dysmetria; (3) patients who remained genetically undiagnosed despite previous testing. Only patients meeting all these criteria were selected for further evaluation. A comprehensive retrospective review of medical records was conducted, and for patients with a family history, samples from affected family members were also collected when available. The study protocol was approved by the Institutional Review Board of Seoul National University Hospital (2004-042-1116 & 2402-149-1518) and was conducted in accordance with relevant guidelines and regulations.

### Cell line culture conditions

HeLa (ATCC, CLL-2) cells were cultivated in Dulbecco’s modified Eagle medium (DMEM) (Welgene, KR; cat. LM001-05) containing 10% fetal bovine serum (FBS) (Welgene; cat. PK004) and 1% antibiotics (Welgene; cat. LS203-1) in a humidified incubator at 37 °C with 5% CO2.

### Design and optimization of gRNA for nCATS in disease-related STR regions

We selected genes associated with STR-related disorders and designed gRNAs to target specific ROIs for nCATS. Our previous research demonstrated that, during nanopore sequencing, shorter DNA fragments are preferentially sequenced compared to longer fragments (*5, 13*). To address this, we designed gRNAs approximately 5000 bp upstream/downstream of both sides of the repeat regions to create DNA fragments of around 10 kb. This design ensures that long DNA fragments containing repeat expansions are sequenced without bias against other genomic regions, enabling equitable representation of the 56 target genes.

To optimize multiplexed gene sequencing using nCATS, we designed gRNAs using Cas-Designer, which evaluates potential off-target sites by searching for genomic sequences that differ by up to two nucleotides (*36–38*). We prioritized gRNAs that presented a single perfect on-target site and no predicted off-targets with mismatches of one or two bases, thereby ensuring high target specificity across nearly all target loci . We concluded that cleavage efficiency is the critical factor, and finalized one pair of gRNAs for each ROI. We added a second pair of gRNAs to enhance coverage for five genes (*ATXN8*, *TBP*, *DAB1*, *NUTM2B- AS1*, and *DIP2B*) that had insufficient sequencing depth.

### Synthesis of gRNAs by *in vitro* transcription

CRISPR RGEN tools (Cas-Designer) were used to design the gRNA sequences (*36, 37*). A list of oligos for the target sequences is provided in **table S3** and **table S4**. The gRNA template oligos were ordered from Macrogen and Cosmogenetech. The gRNAs were synthesized by *in vitro* transcription using T7 RNA polymerase (NEB) and template oligonucleotides. The gRNA product was then purified using the RNeasy mini kit (Qiagen) and quantified using a NanoDrop.

### In vitro cleavage

*Streptococcus pyogenes* Cas9 (SpCas9) nuclease was ordered from Enzynomics (Daejeon, Korea). To generate Cas9 RNP complexes, SpCas9 and gRNA were mixed in a 3:5 ratio and incubated at room temperature for 30 min. For the *in vitro* cleavage (IVC) assay, target regions encompassing the gRNA binding sites were first amplified from HeLa genomic

DNA using PCR with the primers listed in **table S5**. The PCR products were gel-purified and incubated with Cas9 RNP complexes at 37 °C for 10 min. The reaction mixtures were loaded onto 1% Tris–borate–EDTA (TBE) agarose gels, and gel images were captured using a Gel Doc system. Band intensities were quantified with ImageJ, corrected for fragment length, and used to calculate cleavage efficiency. The band intensities were normalized by dividing each value by the corresponding product length.

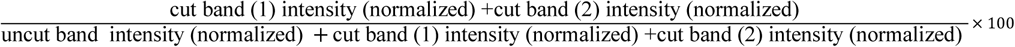

### Library preparation and Cas9-mediated nanopore long-read sequencing

Genomic DNA was extracted from whole blood samples using the Qiagen DNeasy blood and tissue kit (Qiagen; cat. 69504). The crRNAs and tracrRNAs were ordered from Integrated DNA Technologies (IDT, Coralville, Iowa, USA), and the Cas9 nuclease was purchased from Enzynomics and IDT. We prepared sequencing libraries from 10 µg of genomic DNA using the SQK-LSK114 kit (Oxford Nanopore Technologies, UK) and a Cas9- mediated targeted enrichment protocol (*39*). To increase the consistency and depth of the on-target sequencing coverage, we modified the standard Oxford Nanopore Technologies (ONT) protocol by separating the Cas9 cleavage and dA-tailing steps. While the original protocol performs both steps simultaneously, we instead removed the Cas9 protein via bead purification after cleavage, followed by dA-tailing using the dA-tailing module (NEB, E6053L) (**Fig. S3**). Prepared libraries were loaded onto flow cells (R10.4.1) and sequenced using the MinION platform (Oxford Nanopore Technologies).

### Coverage analysis

Coverage depth for each ROI was calculated from the aligned BAM files as the number of mapped bases divided by the target length (bp). To evaluate the overall sequencing performance, we computed two metrics: i) total on-target bases by summing the product of coverage depth and ROI length across all targets using the following formula:

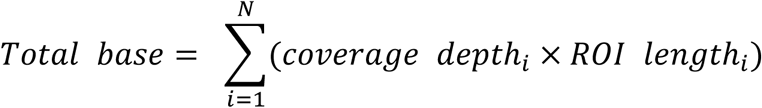

where *N* denotes the number of ROIs included in the analysis. This represents the total number of aligned bases within the targeted regions and reflects the on-target sequencing yield per condition. ii) To account for differences in ROI size when comparing average sequencing depth across conditions, the length-normalized average coverage depth was calculated as follows:

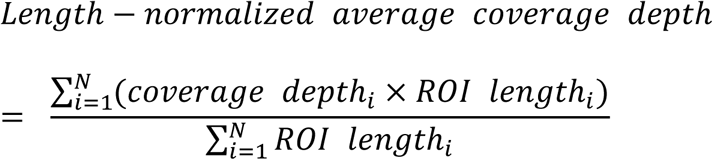

this approach minimizes length-dependent bias and represents the mean sequencing depth per target after normalization for ROI size. In contrast, for per-locus comparison, raw coverage depth values without length normalization were used to visualize coverage distribution across individual STR loci.

### Preparation of BED files for adaptive sampling

To facilitate adaptive sampling, a gene panel was designed, and its corresponding BED file was created (**table S6**). Given the potential variability in cleavage points at the terminal regions of the library DNA, which could affect alignment during adaptive sampling, the BED file was configured to extend 100 bp upstream and downstream of the cleavage points. This adjustment ensures that even if the cleavage points deviate slightly from the expected range, the corresponding library fragments will not be prematurely ejected during the adaptive sampling process, enabling accurate classification.

### Analysis of nCATS data and visualization

Sequencing was performed using MinION R10.4.1 flow cells, and the resulting POD5 files were base-called using the Dorado software (version 0.6.2). (https://github.com/nanoporetech/dorado). Subsequently, reads were aligned to the GRCh38 reference genome using minimap2 (version 2.28-r1209) (https://github.com/lh3/minimap2) (*40*). Subsequently, BAM files were sorted and indexed using Samtools (Version 1.18) (https://www.htslib.org/) (*41*).

Based on a known table of repeat expansion information (**table S7**), a motif analysis and methylation state analysis were conducted on the reads within the ROIs. It has been well-established that the phenotype of diseases caused by repeat expansion can be influenced by the presence of non-motif sequences between consecutive motifs, a phenomenon referred to as the "interruption effect" (*42*).

To account for this, both the highest consecutive repeat count and the total repeat count were investigated. Due to the relatively high error rate of nanopore sequencing and the variation in repeat counts within samples, confirming that the observed interruption effect was not a false positive was essential. Thus, visualization techniques were employed to distinguish between motifs and non-motif sequences, verifying the occurrence of interruptions.

### STRiker algorithm for STR analysis

We developed STRiker (https://github.com/BaeLab/STRiker), a computational pipeline for the comprehensive STR analysis of sequencing data. The algorithm accepts aligned reads (BAM format), reference genome (FASTA), and STR region coordinates with known pathogenic motifs (CSV).

Motif discovery: STRiker employs dual motif discovery strategies. First, reference sequences were extracted with flanking regions and scanned using a sliding window (3–30 bp) to identify tandem repeats that appeared consecutively ≥3 times. Second, *de novo* motif discovery was performed directly from sequencing reads overlapping each STR region. To handle circular permutations inherent to tandem repeats, all motifs were initially canonicalized to their lexicographically smallest rotation. However, if a discovered motif matched any rotation of a known pathogenic motif, this motif was reassigned to the known motif form to maintain clinical relevance. For example, if CAG was the known pathogenic motif and AGC was discovered, then AGC was reported as CAG rather than AGC.

Pattern decomposition: Sequences are decomposed using a greedy algorithm that matches the longest motif at each position, considering all rotational variants. When multiple rotational variants are possible, preference was given to known pathogenic motifs. Consecutive identical motifs were grouped to simplify patterns. For example, a sequence containing CAGCAGCAGCAACAGCAG was represented as ((CAG, 3), (CAA, 1), (CAG, 2)), explicitly capturing interruption patterns that may modulate disease severity.

Repeat quantification: For each STR locus, repeat counts were calculated by summing occurrences of the primary pathogenic motif and its rotational equivalents across all reads. Only reads that fully spanned both boundaries of the STR region were included in the analysis to ensure accurate repeat counting. Kernel density estimation was used to visualize repeat length distributions, enabling detection of expanded alleles and mosaicism. Pathogenic expansions were identified by comparing observed repeat counts against established clinical thresholds.

Quality control: Quality metrics included coverage depth with a minimum threshold of 30× for confident variant calling, motif concordance between reference and *de novo* discovery, and pattern complexity assessment. The pipeline outputs were as follows: (1) an Excel file comparing reference and *de novo* motifs with occurrence frequencies, (2) a PDF report with pattern visualizations and repeat distributions, and (3) coverage statistics for each analyzed locus.

Performance evaluation was conducted on a workstation equipped with an Intel Core i7-13700K processor, 32 GB of RAM, and running Ubuntu 20.04. STRiker was executed using 4 CPU cores with multiprocessing enabled, with the following parameters: minimum motif length of 3 bp, maximum motif length of 30 bp, minimum consecutive motif count of 10, and minimum motif coverage of 5 (**table S8**). Processing time was measured using the Unix time command across three independent runs for each dataset, with real time (wall-clock time) recorded as the primary performance metric (**table S9**).

### Statistical analysis

Statistical analyses were primarily descriptive and focused on summarizing assay performance and clinical diagnostic yield rather than formal hypothesis testing. Continuous variables (such as age at onset, time from symptom onset to genetic diagnosis, and repeat counts) were summarized using means and standard deviations, medians, and ranges as appropriate; categorical variables (such as sex, presence of a family history, and presence of specific STR expansions) were summarized as counts and percentages.

For *in vitro* optimization experiments, normalized cleavage efficiencies were calculated from ImageJ-quantified, length-corrected band intensities as described in the “*in vitro* cleavage” section. Sequencing coverage metrics for each region of interest, total on-target bases, and length-normalized average coverage depth were calculated from aligned BAM files as described in the “Coverage analysis” section, and coverage uniformity across loci and experimental conditions was assessed by comparing distributions of raw and length-normalized depth values rather than by formal hypothesis testing. STRiker-generated repeat-length distributions and motif compositions (including reference and *de novo* motifs) were analyzed using custom Python scripts integrated within the STRiker pipeline.

## List of Supplementary Materials

Fig S1 to S6 for multiple supplementary figures Table S1 to S9 for multiple supplementary tables

## Acknowledgements

Most of the sequencing data analysis was conducted using the computing server at the Genomic Medicine Institute Research Service Center.

## Author contributions

S.B. and J.M. conceived the project. C.J. developed the bioinformatics algorithms. M.K. and N.K. performed the cell and sequencing experiments. S.L., C.J., and G.-H.H. analyzed the sequencing data. S.L., S.-T.L., K.C., S.K.L., H.-J.K., J.H.C., and J.M. enrolled and clinically evaluated the study patients. S.B. and J.M. supervised the project. S.L., C.J., M.K., and S.B. wrote the manuscript with input from all authors.

## Data and materials availability

All data are available in the main text or the supplementary materials. The code related to this study is available at https://github.com/BaeLab/STRiker.

## Supplementary Materials

**Fig. S1.**
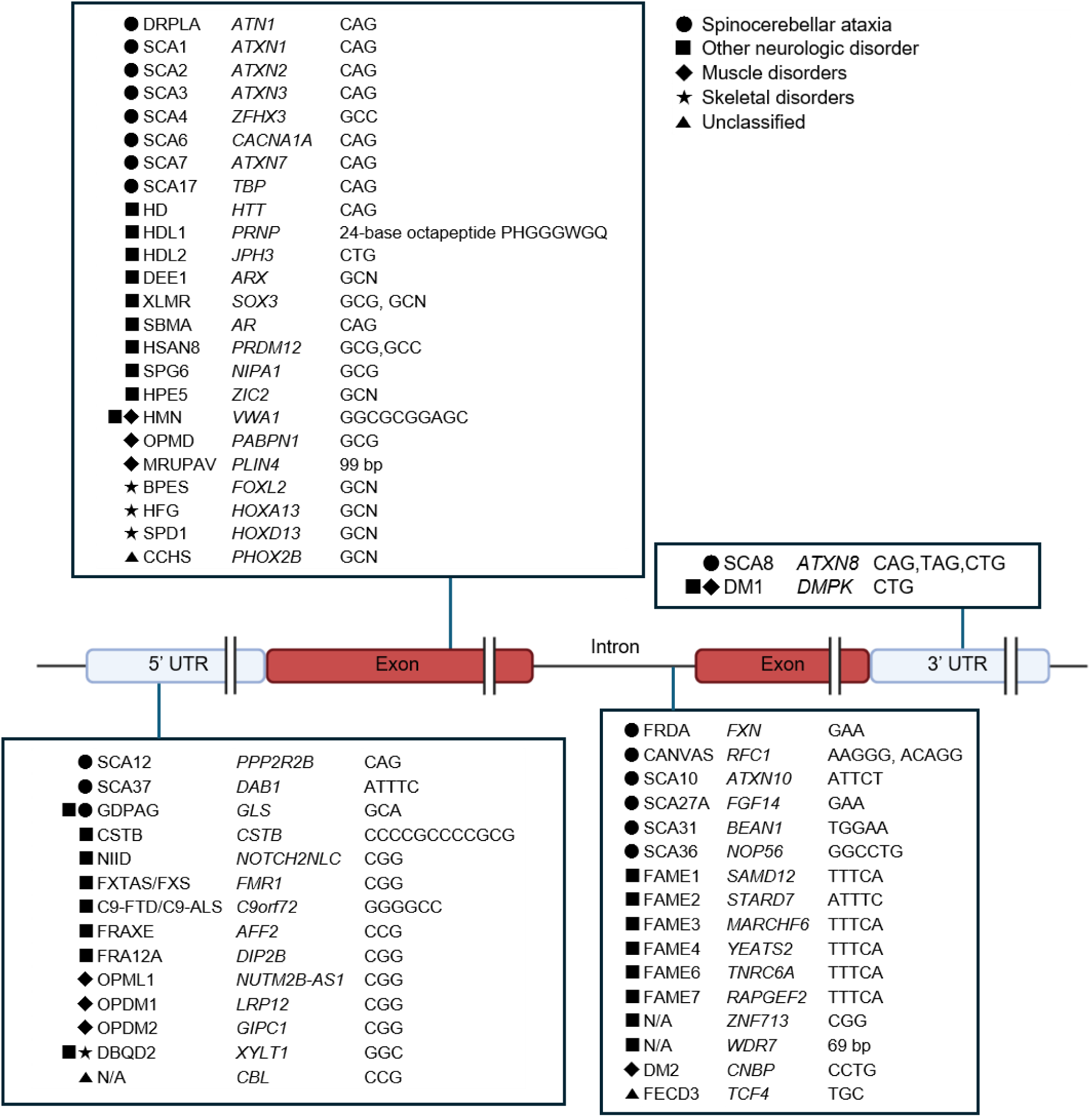
Overview of STR loci included in the 56-gene panel. Schematic representation of disease-associated short tandem repeat (STR) loci categorized by their genomic location (5′ untranslated region (UTR), exon, intron, or 3′ UTR) and repeat motif type. Each box lists representative disorders, associated genes, and repeat motifs. Symbols indicate the major clinical categories. This panel comprises 56 known pathogenic or candidate STR loci encompassing a wide range of repeat motifs implicated in neurological, muscular, and developmental disorders. Created with BioRender.com

**Fig. S2.**
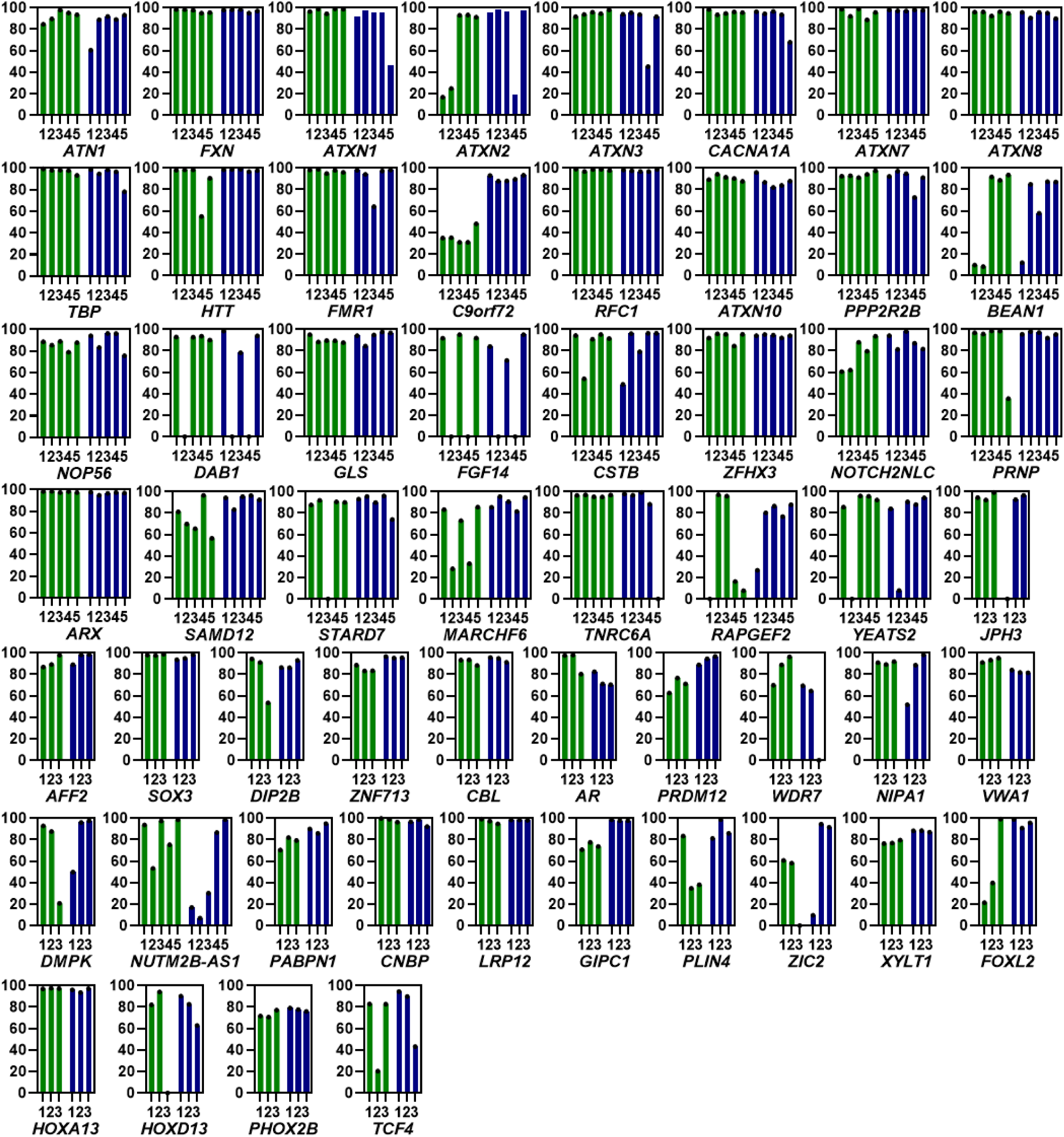
sgRNA screening for high cleavage efficiency via *in vitro* cleavage assay. Cleavage efficiency of sgRNAs targeting upstream and downstream regions of repeat expansion disease-associated gene regions of interest (ROIs), assessed by *in vitro* cleavage assay. Three or five different sgRNAs were tested for each region.

**Fig. S3.**
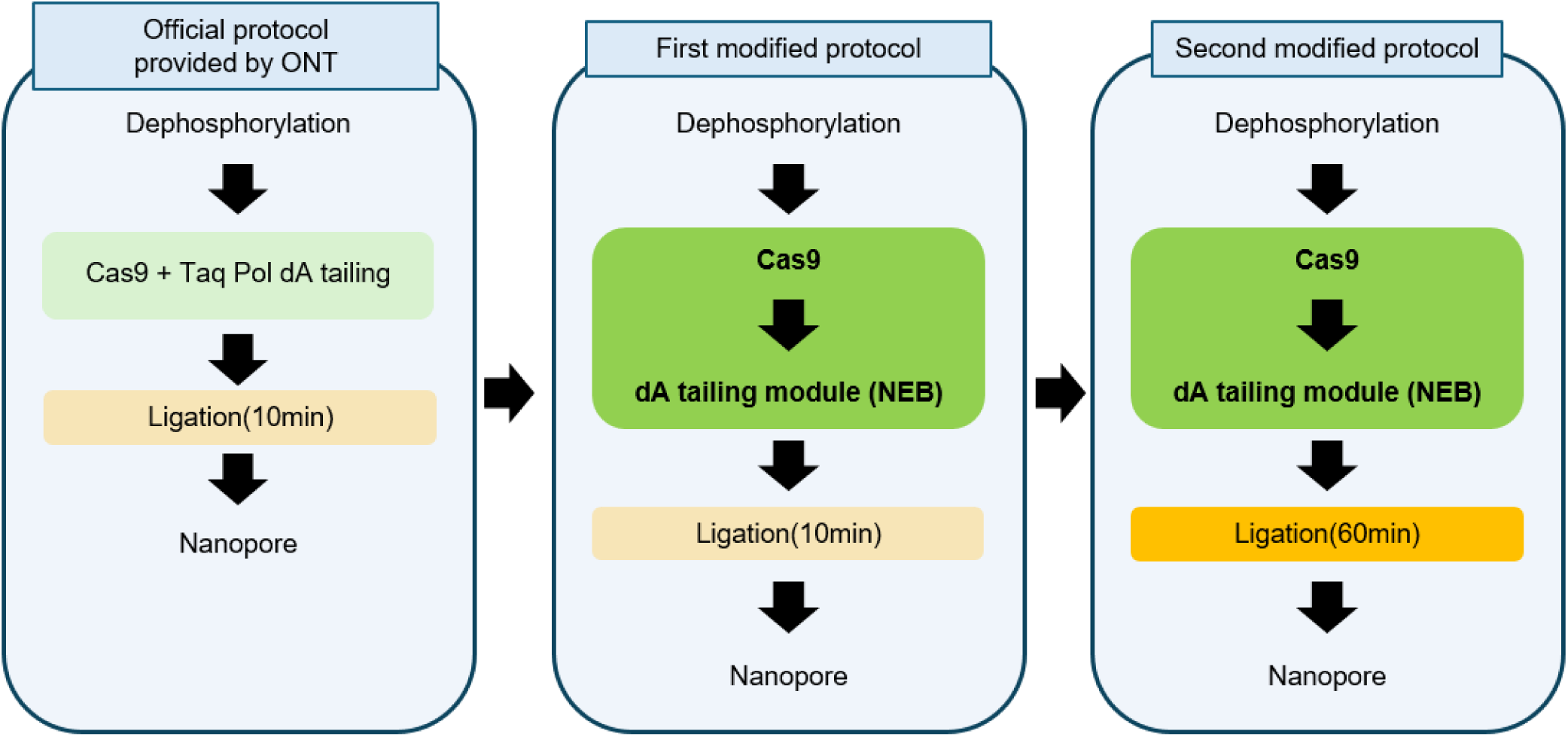
Schematic diagram with a comparison of the optimized nCATS protocol with the conventional protocol. The optimized protocol employs sequential Cas9 cleavage at both ROI termini, followed by bead purification, rather than simultaneous Taq polymerase and Cas9 treatment. Further optimization includes dA-tailing using a Klenow fragment (exo-) (NEB) and an extended nanopore adapter ligation time to 60 minutes.

**Fig. S4.**
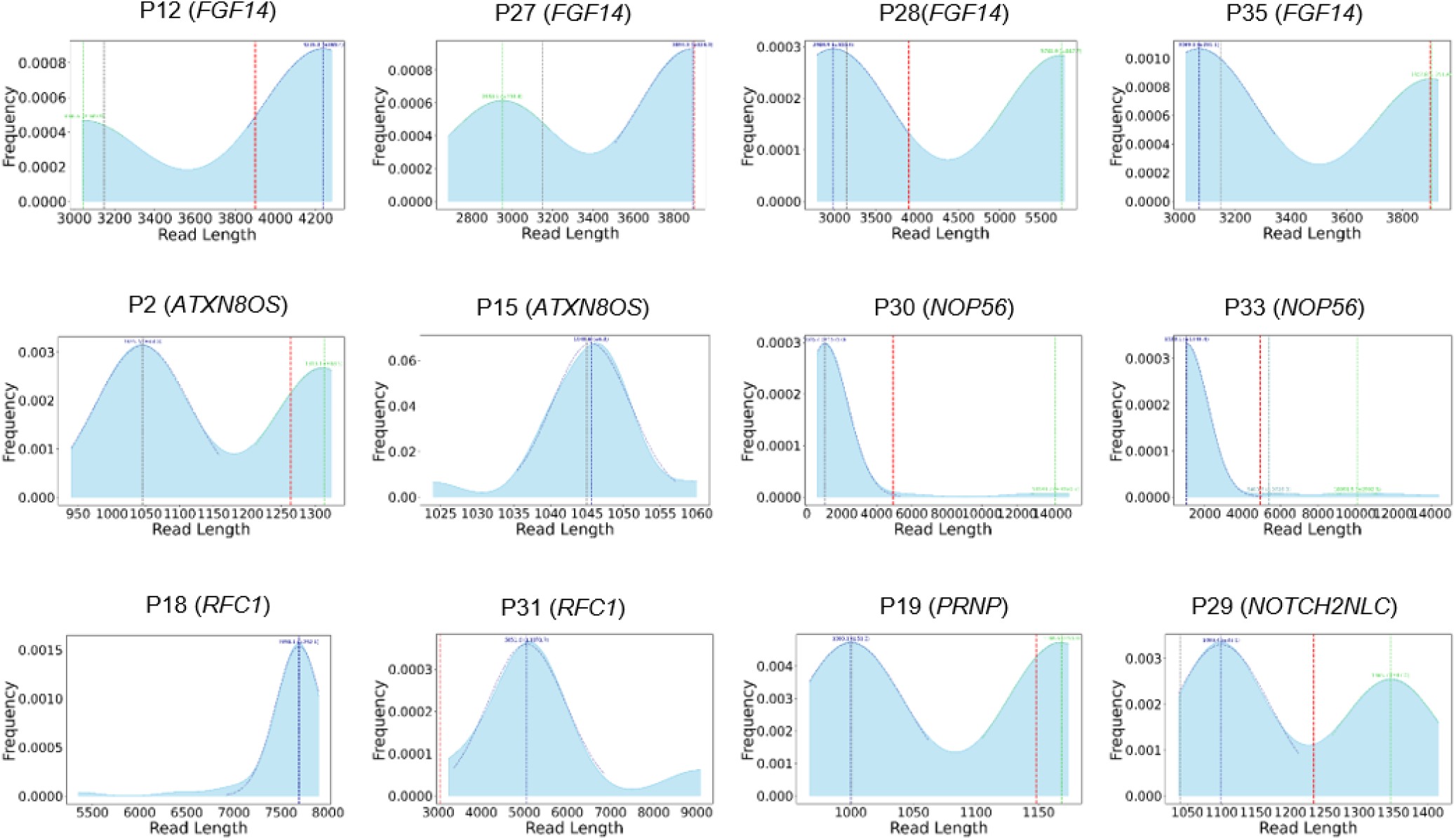
KDE read length distributions at expansion sites for newly diagnosed patients using the STRiker pipeline. Read lengths span the repeat locus plus 500 bp flanking regions on each side. Gaussian-fitted peaks are shown in blue (primary) and green (secondary). The red line indicates the threshold for pathogenic expansion.

**Fig. S5.**
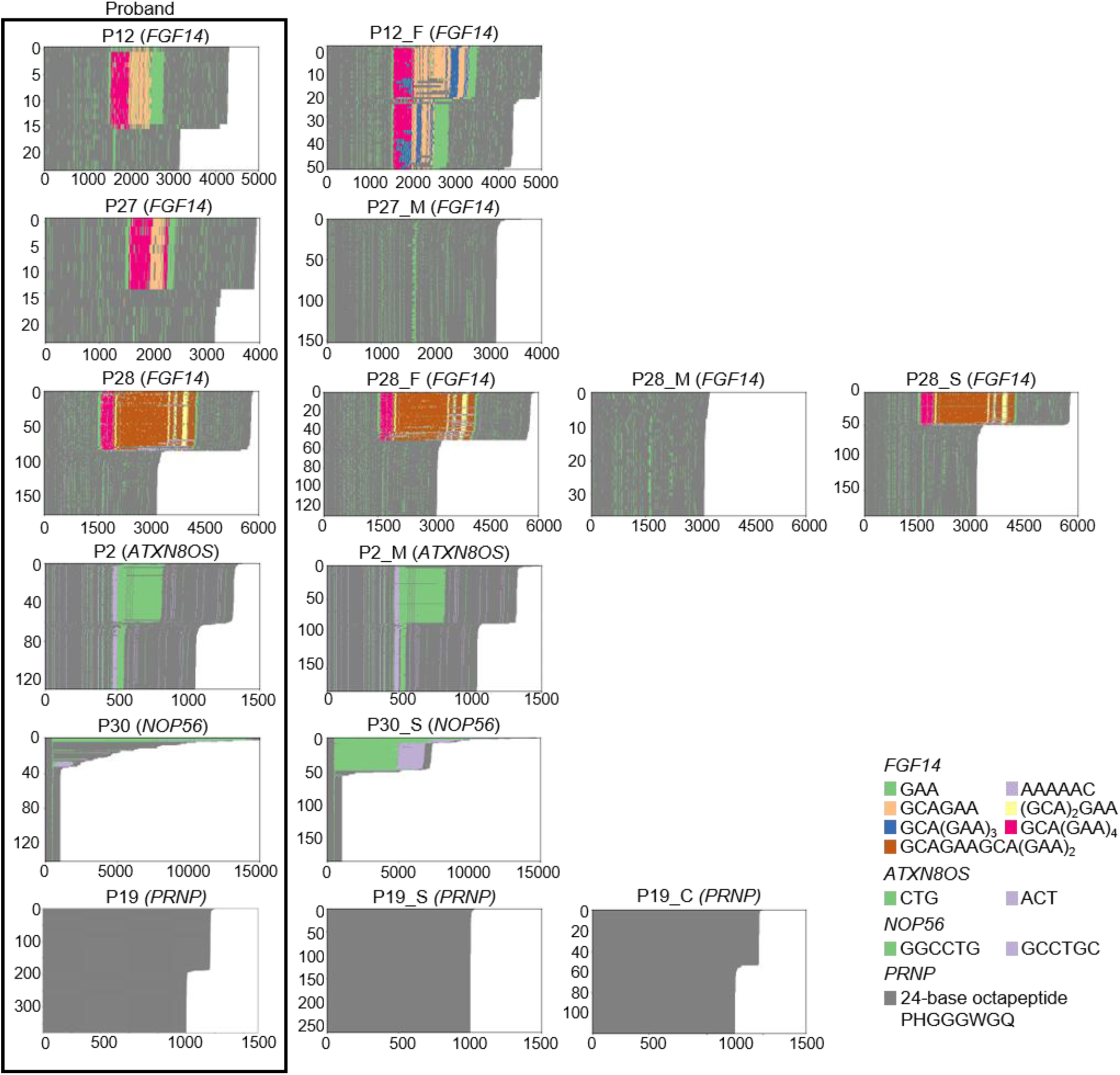
Repeat motif composition analysis in family members identified through cascade screening. Heatmaps of the repeat motif composition for newly diagnosed patients and their relatives identified by cascade screening. Each panel represents long-read alignments at disease-associated loci, with distinct colors indicating different motif types for each gene. *PRNP* repeat expansions consist of an octapeptide motif of 24 bp (PHGGGWGQ), for which no distinct motif colors are applied.

**Fig. S6.**
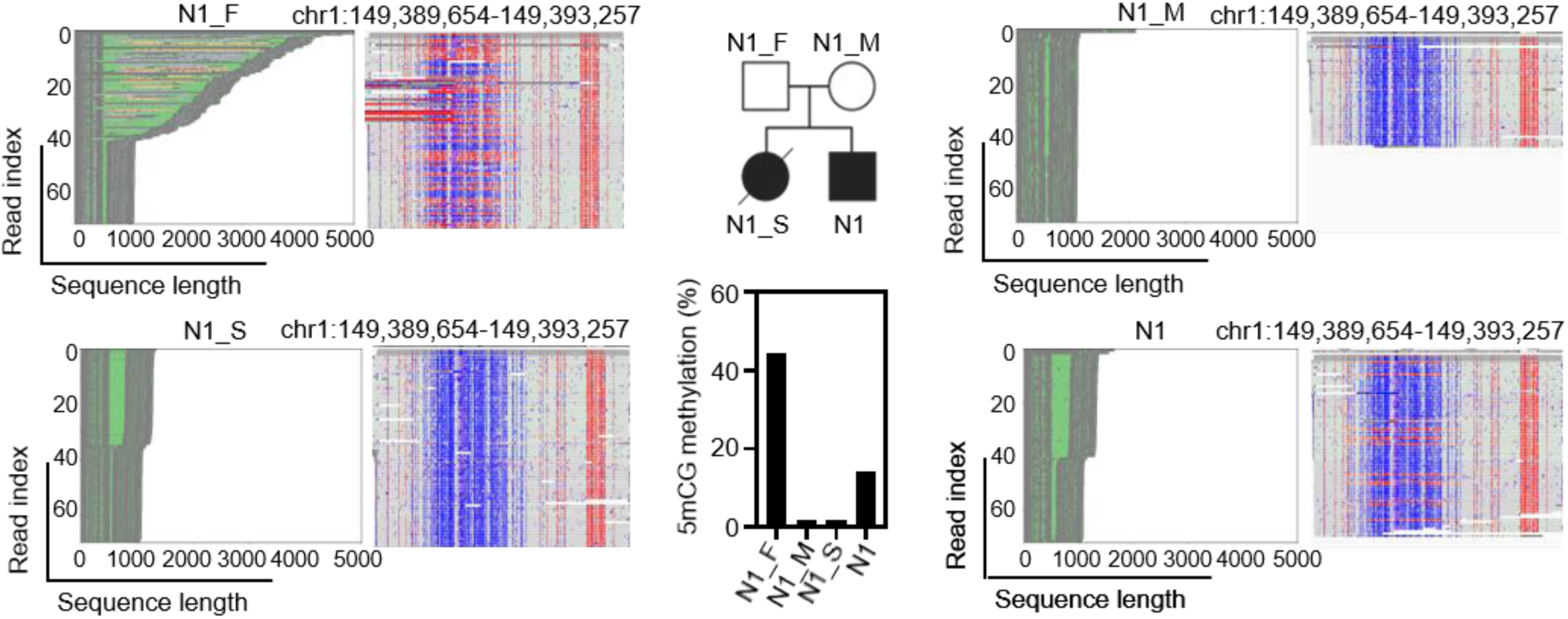
Case of contraction during paternal inheritance in a family with *NOTCH2NLC*-related neuronal intranuclear inclusion disease (NIID). Long-read alignment and methylation profiles of each family member (N1_F, N1_M, N1, and N1_S). The father (N1_M) carried a long GGC repeat with hypermethylation and was asymptomatic, whereas the patient (N1) inherited a shorter allele with hypomethylation, which is likely associated with the development of neuronal intranuclear inclusion disease (NIID). The older sister (N1_S) of the patient also carried a hypomethylated expanded allele, was clinically affected, and died. Created with BioRender.com

**Table S1.**
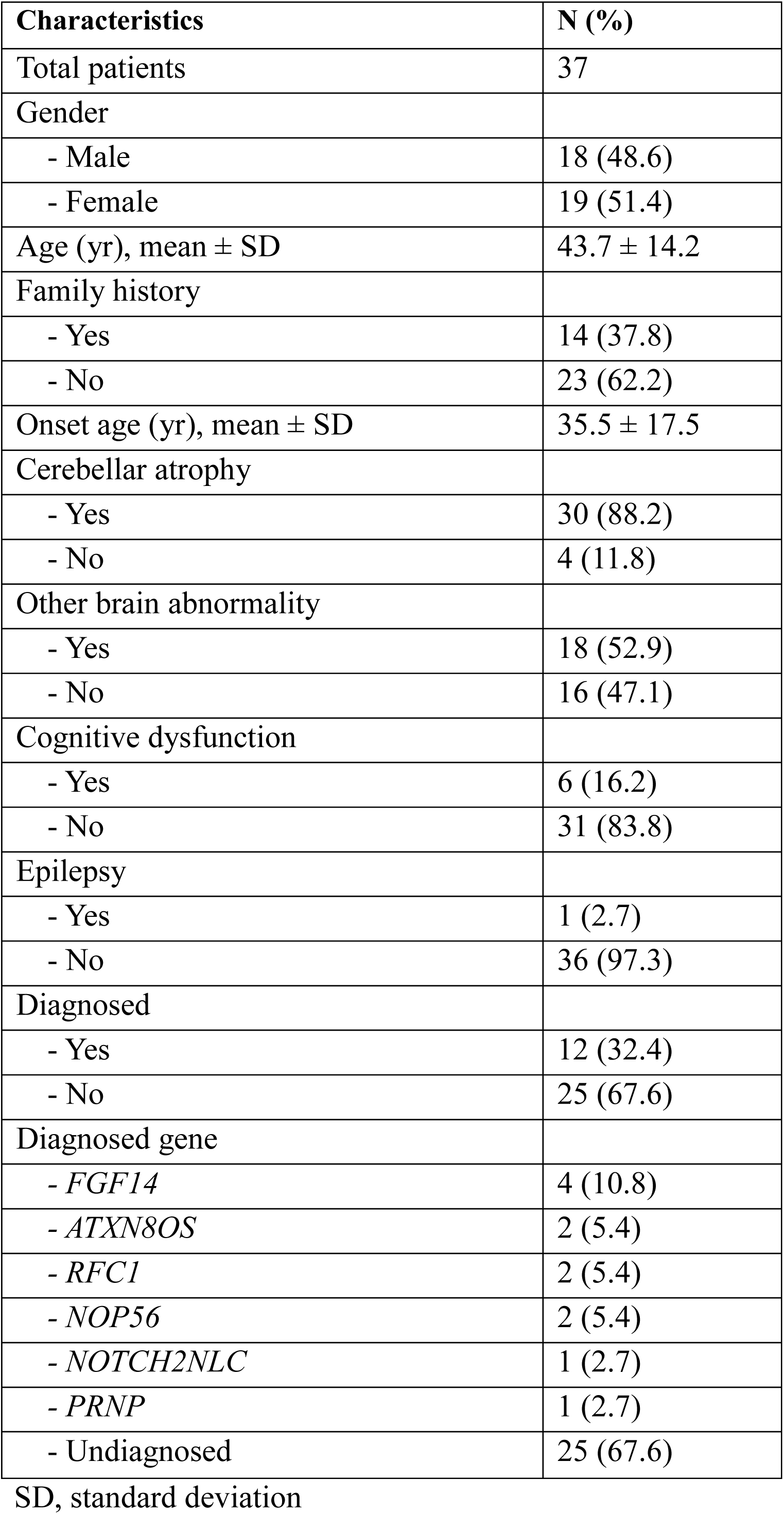
Patient demographics and clinical characteristics.

**Table S2.**
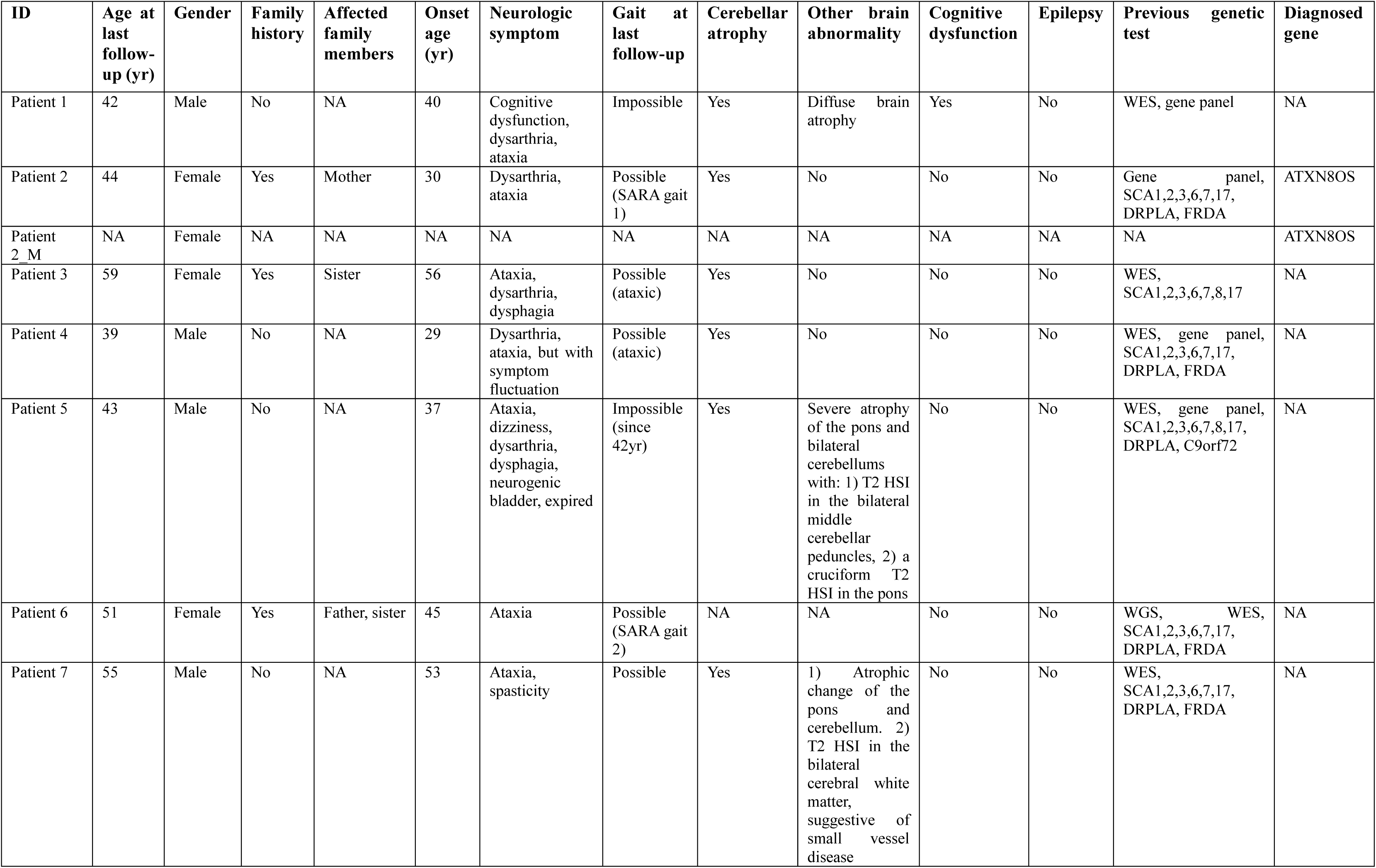
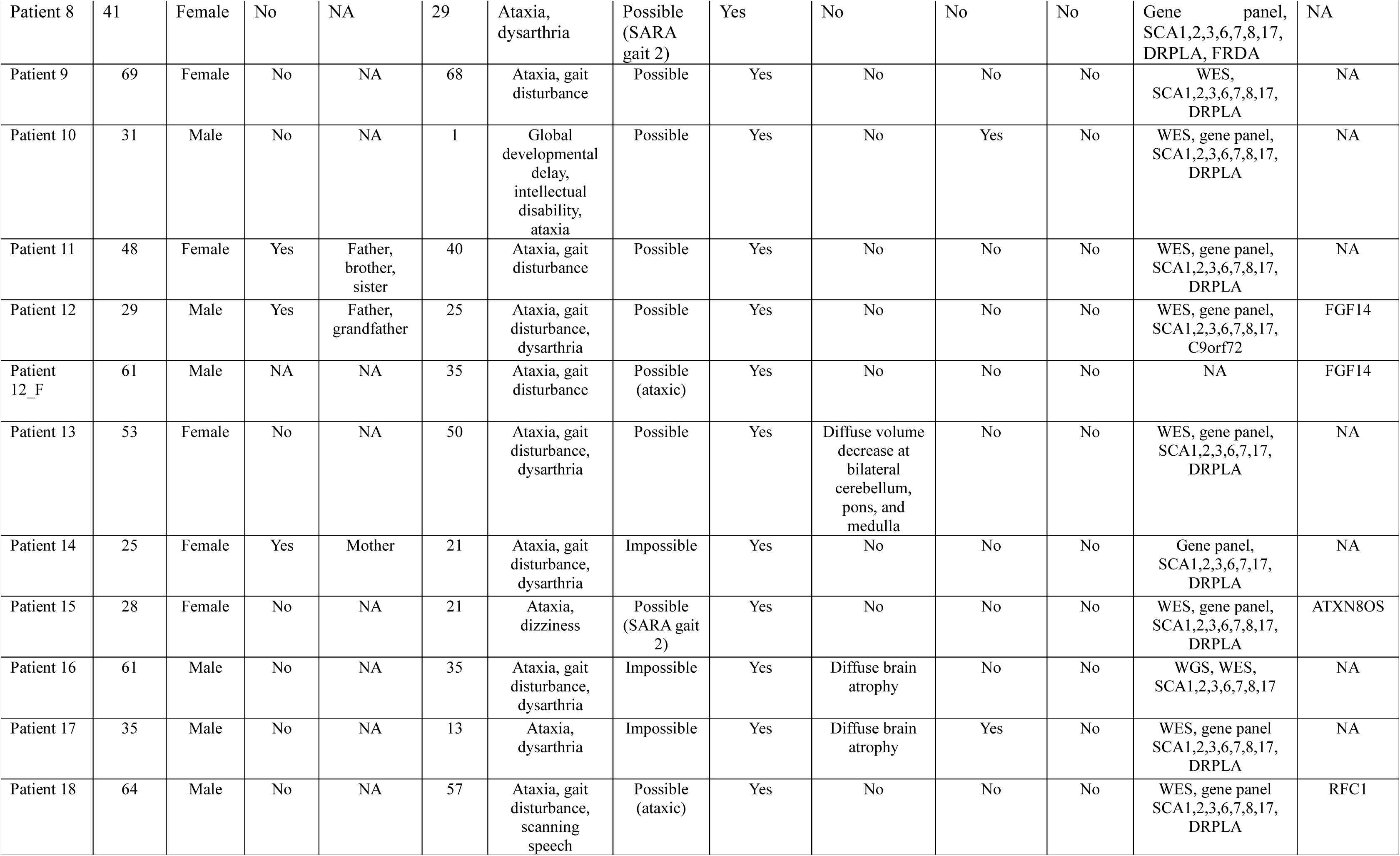
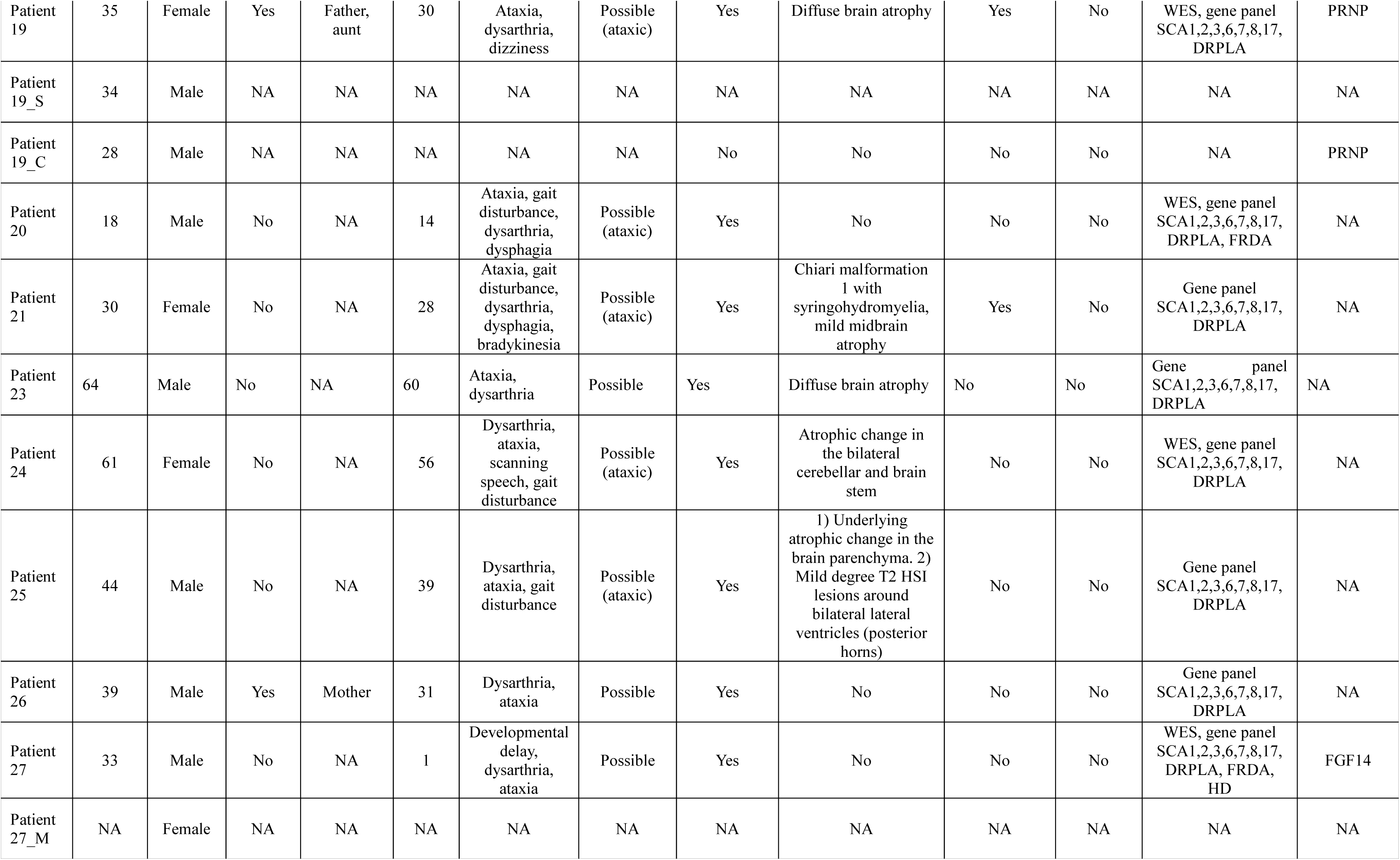
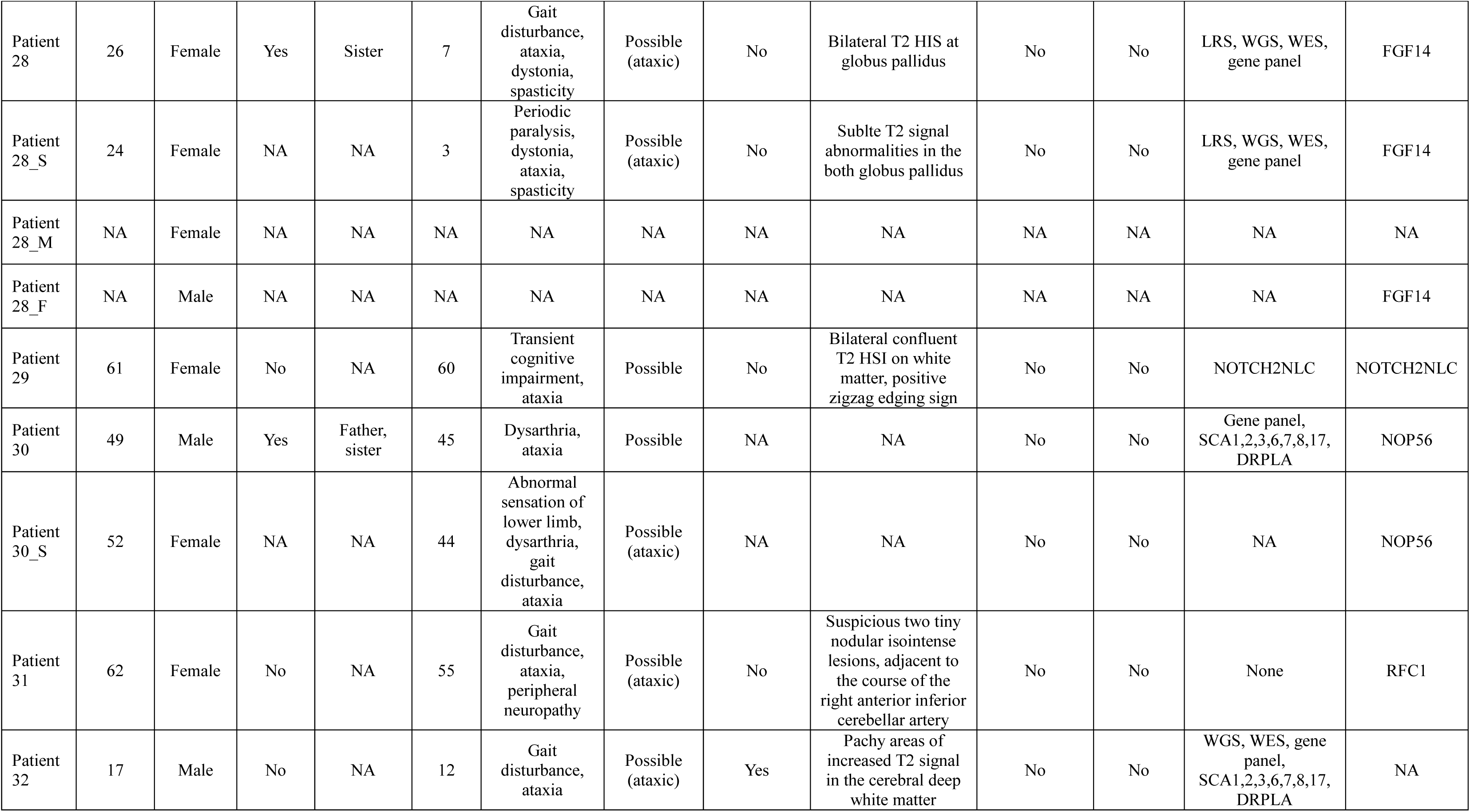
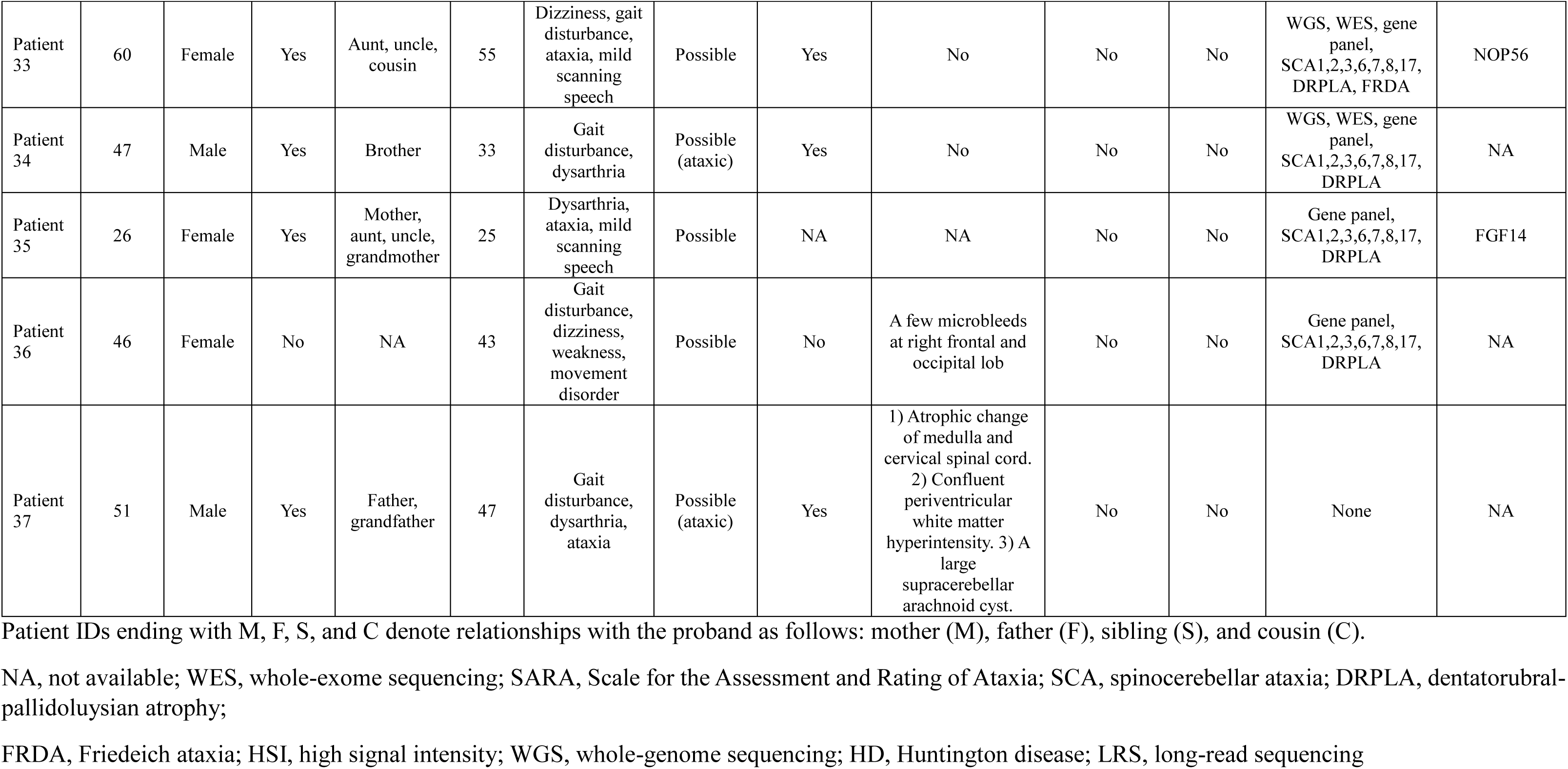
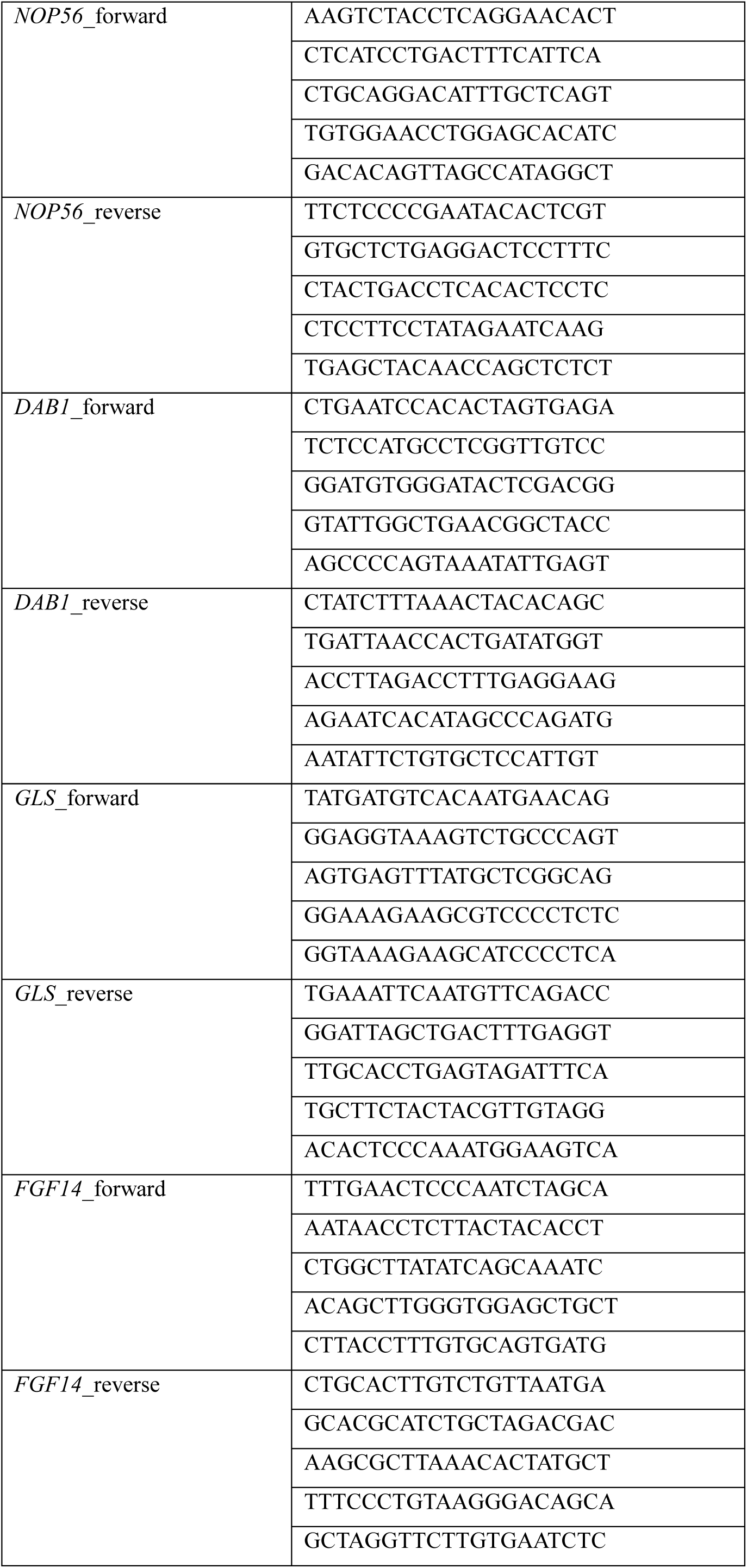
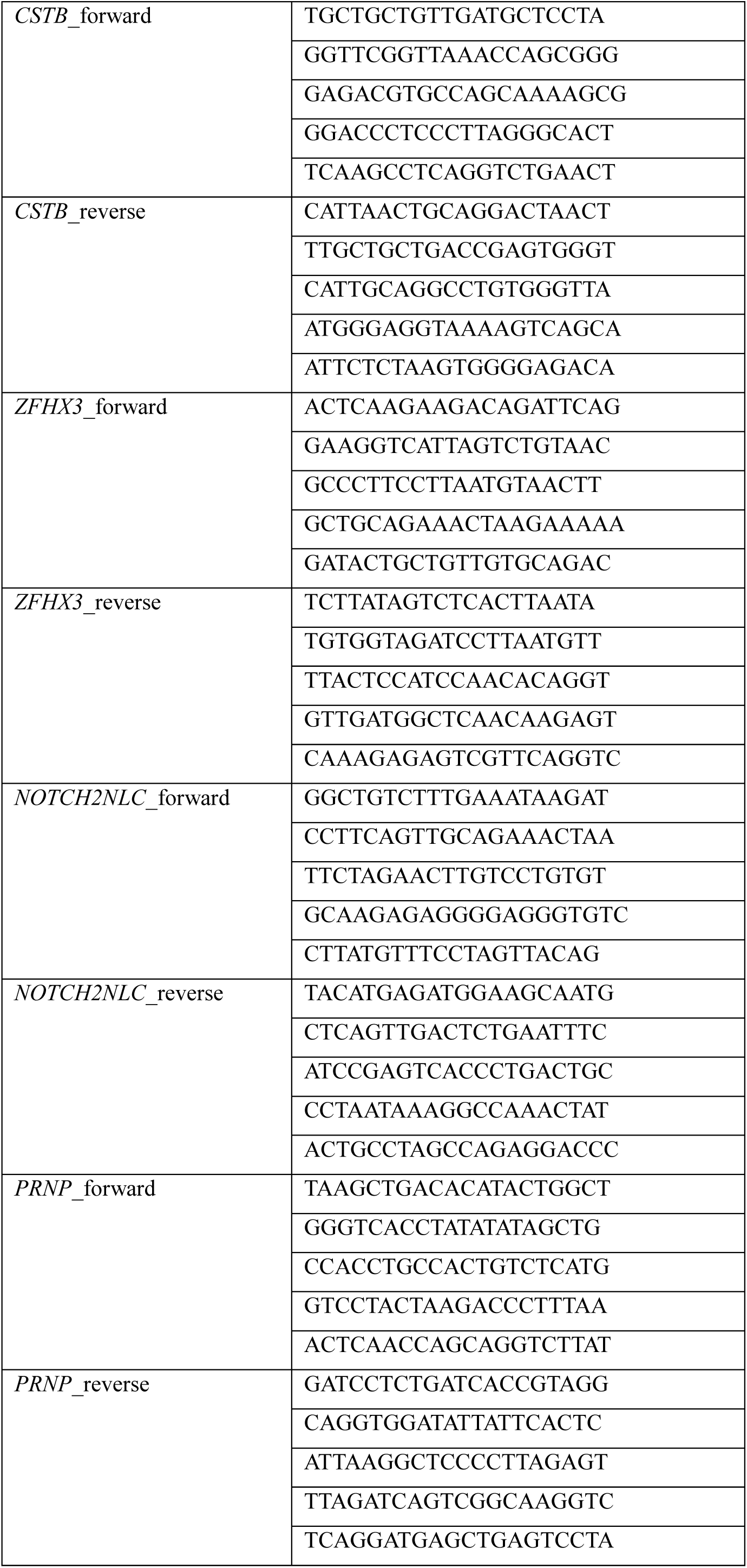
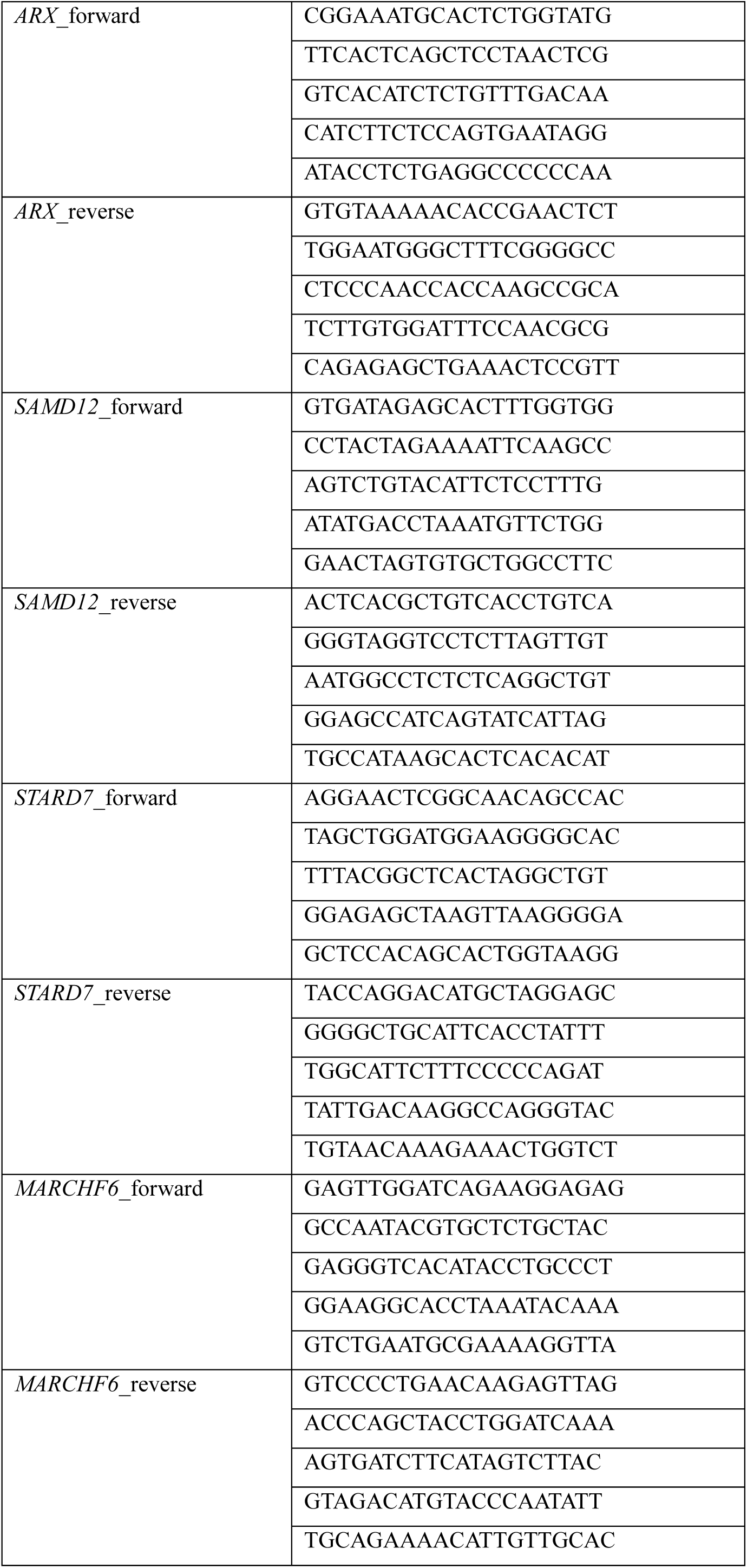
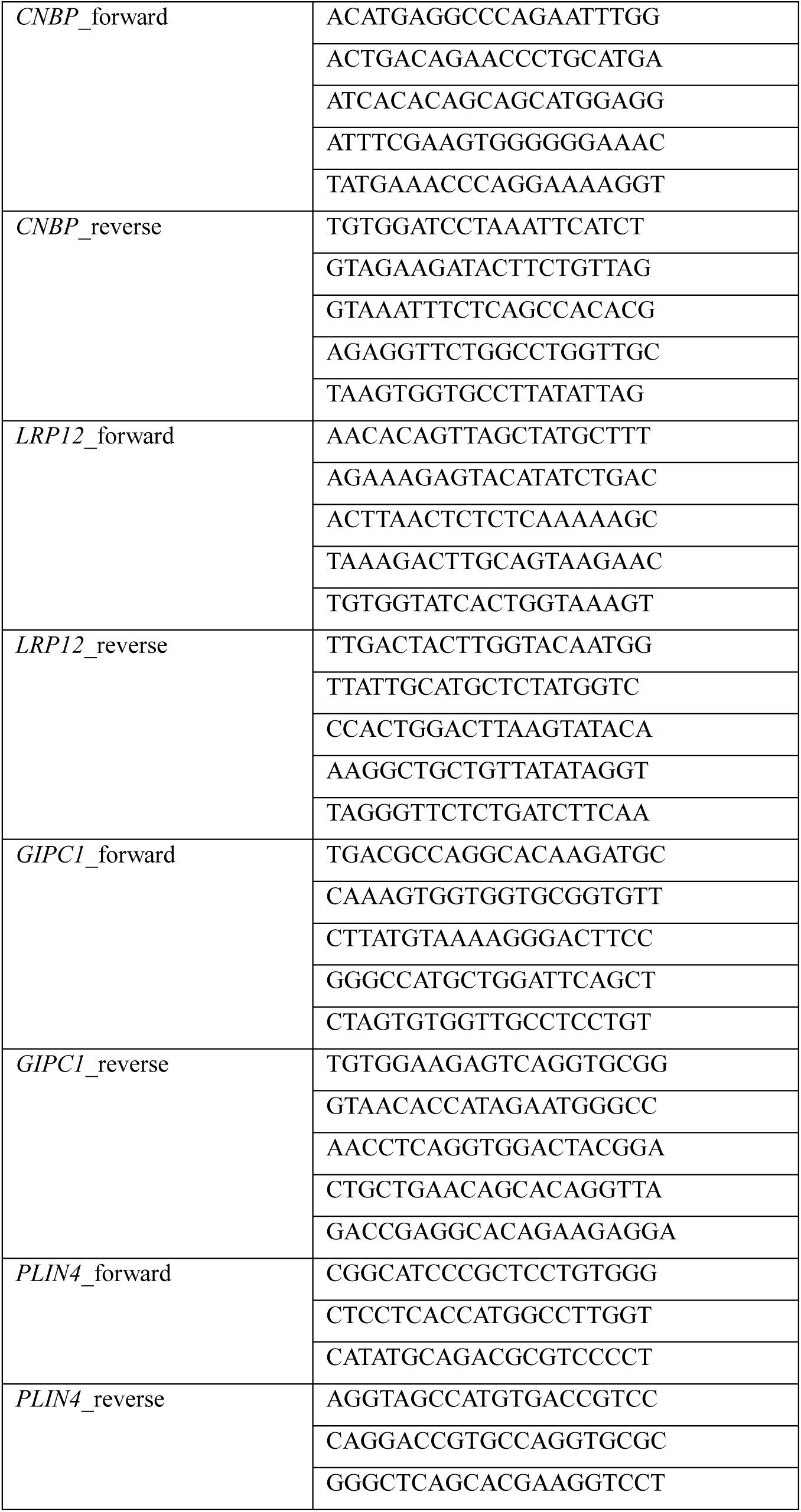
Clinical and genetic information of study participants.

**Table S3.**
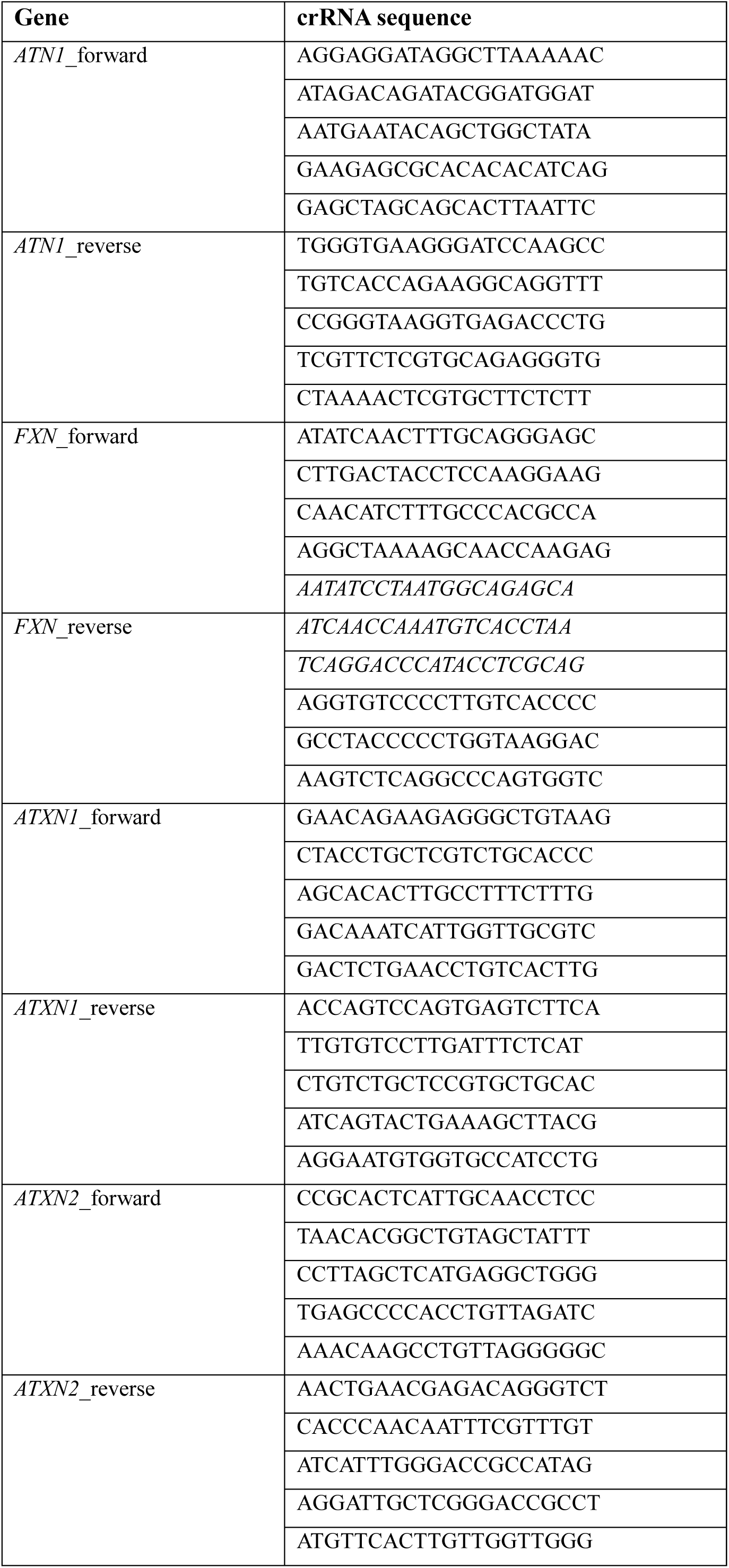
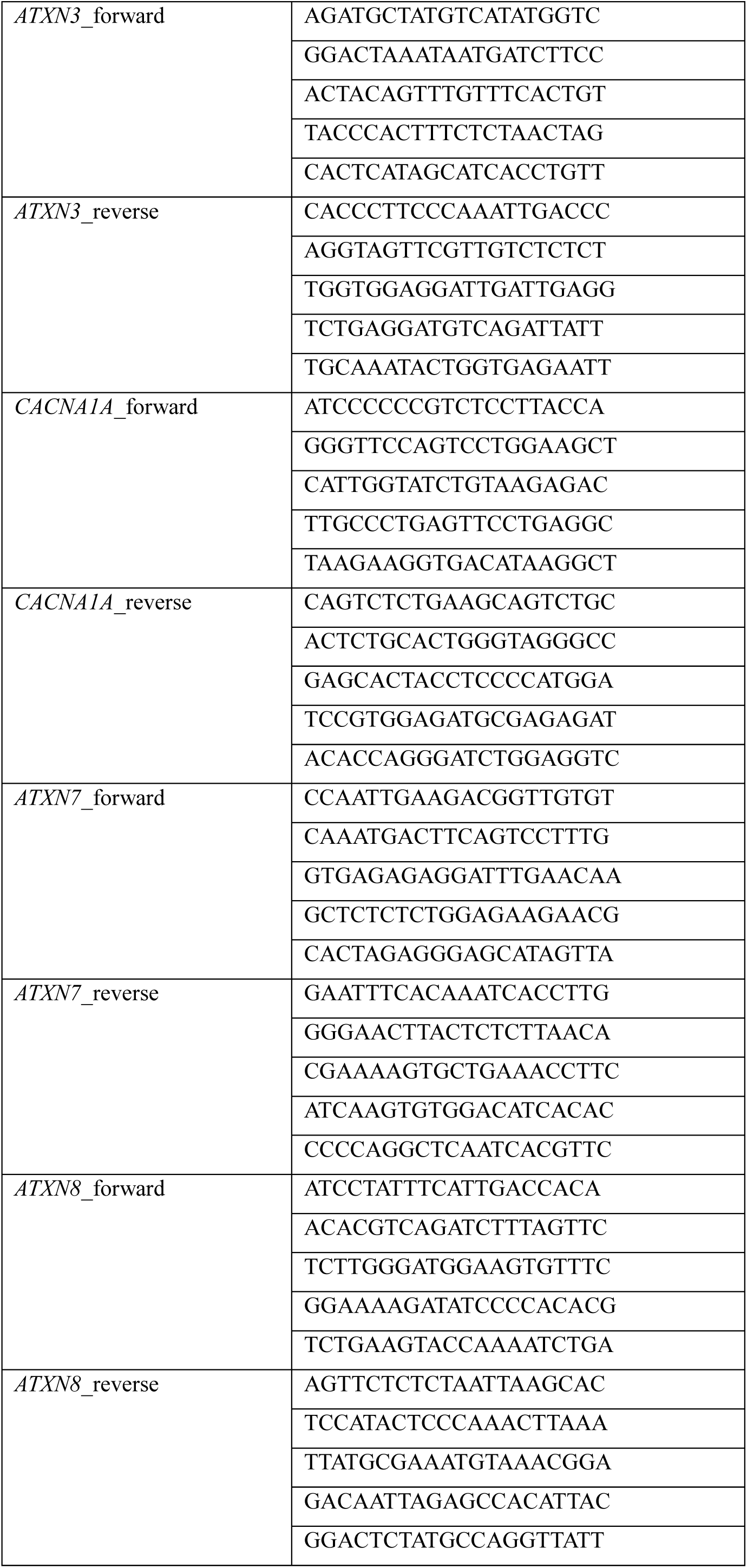
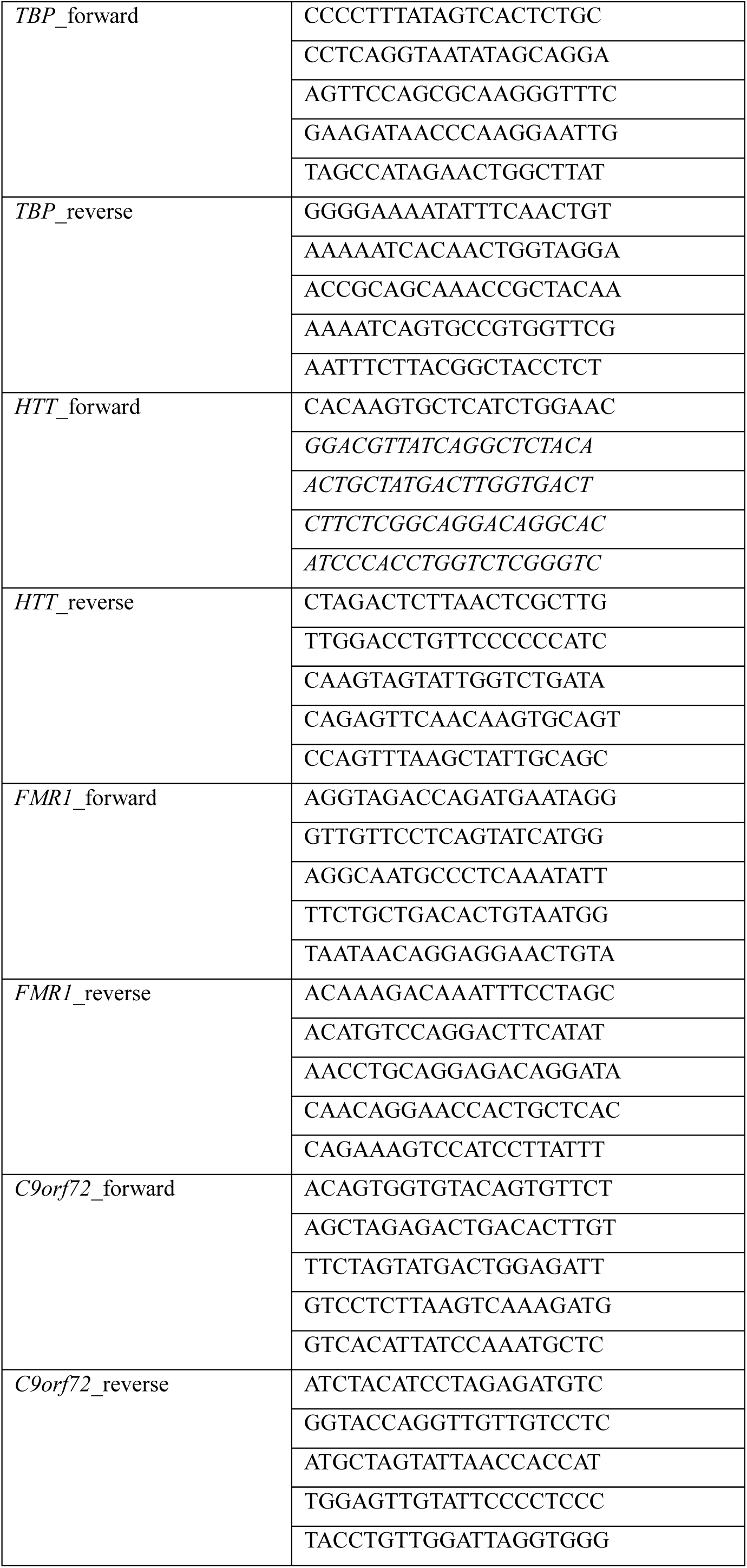
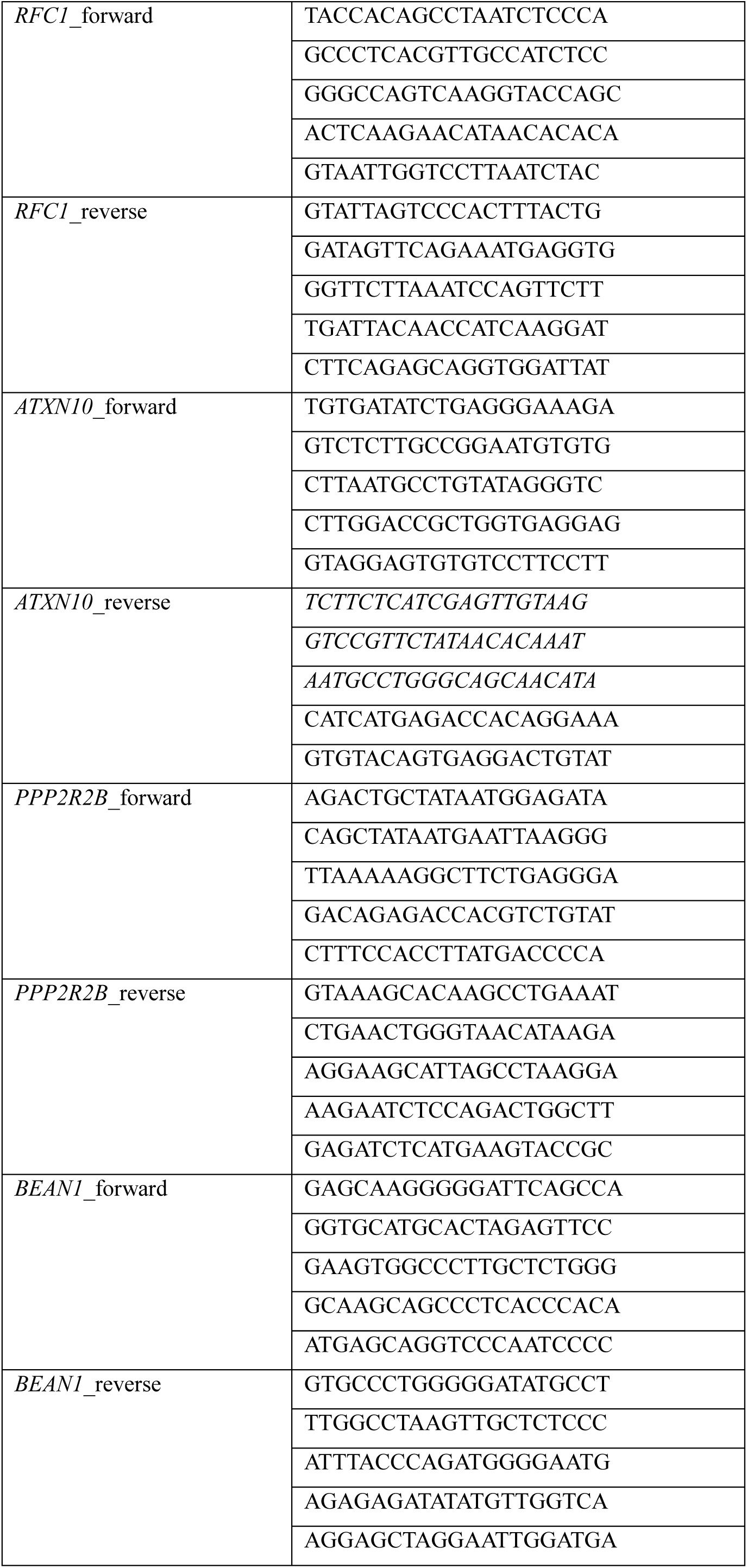
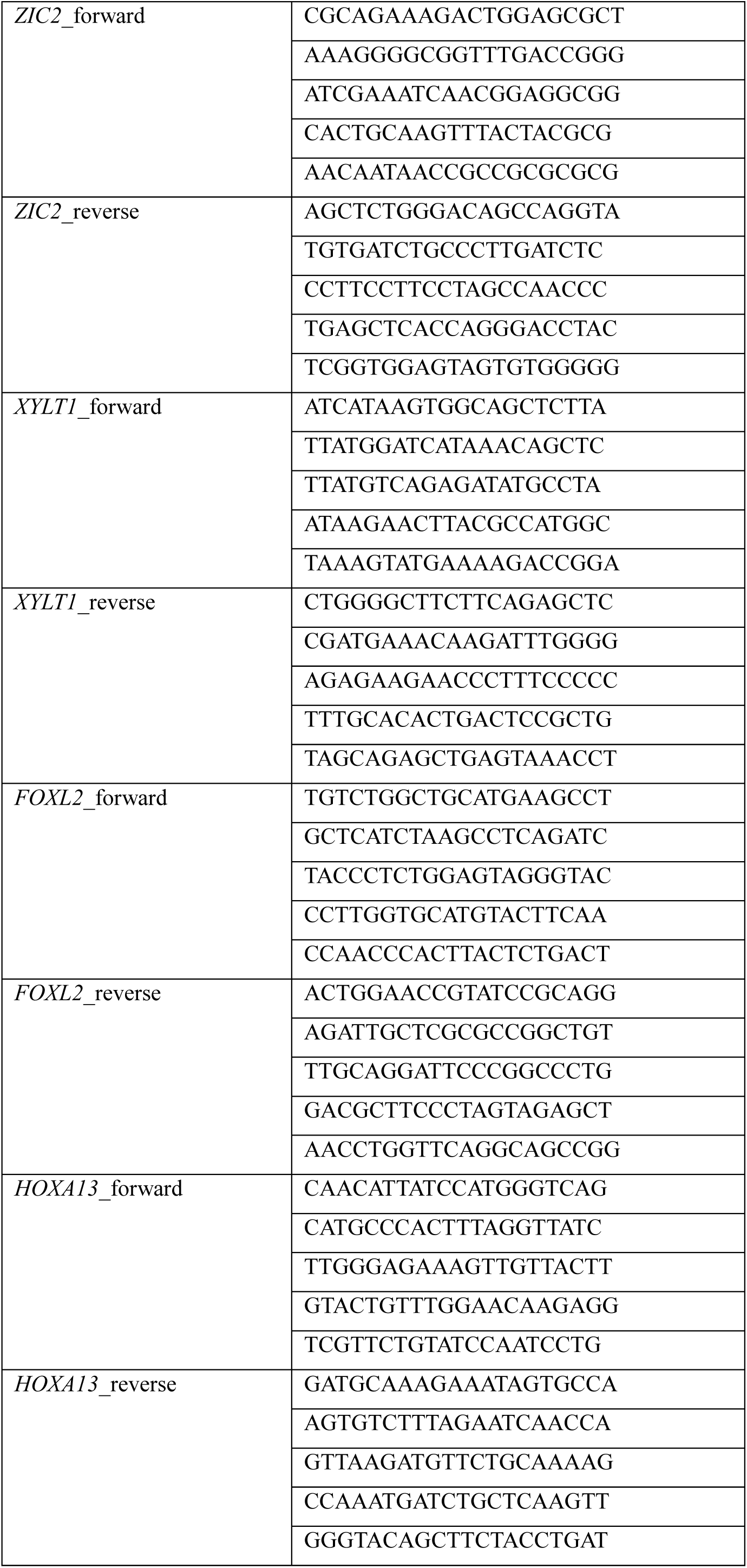
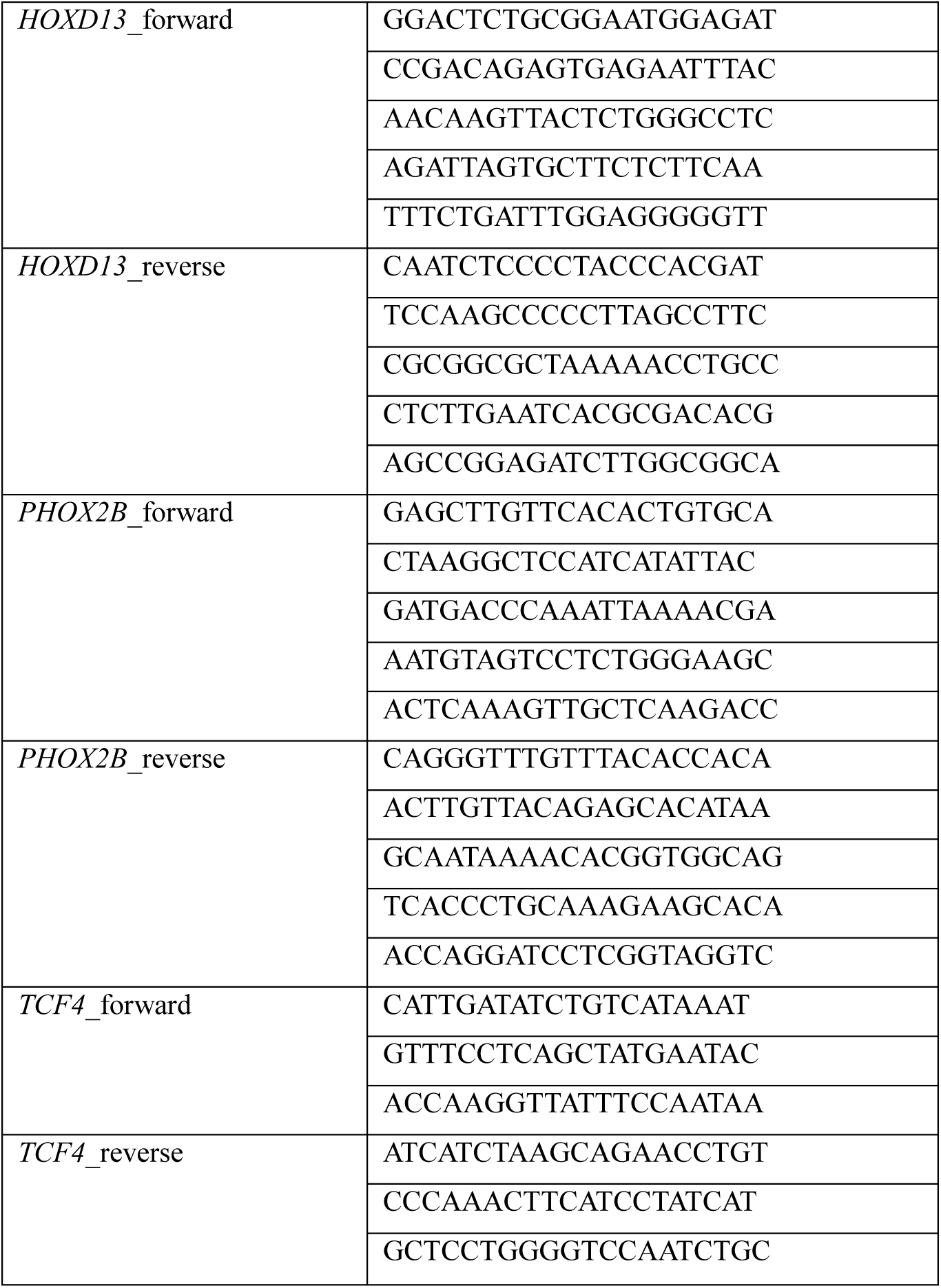
All crRNAs for nCATS determined through in vitro cleavage.

**Table S4.**
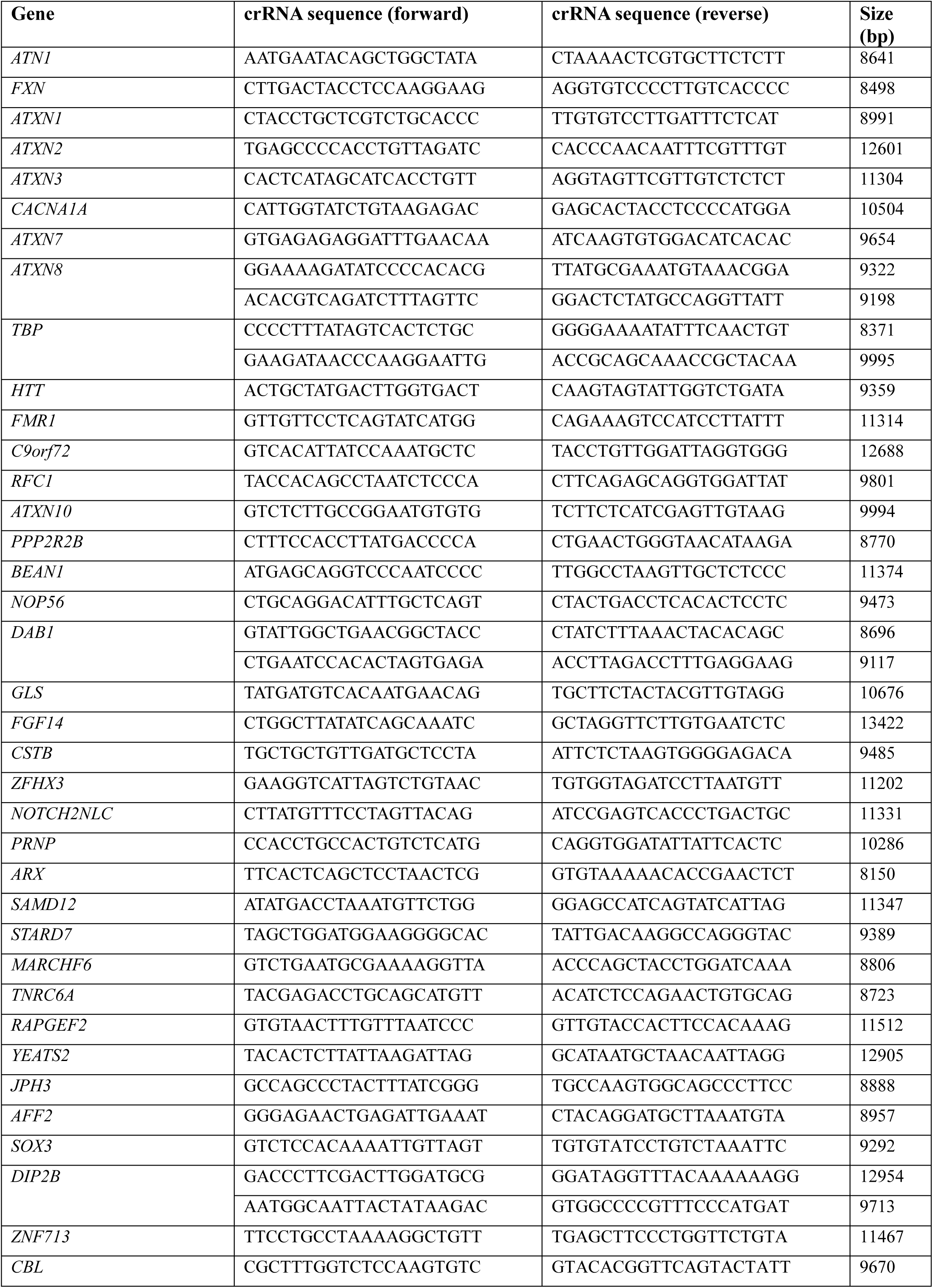
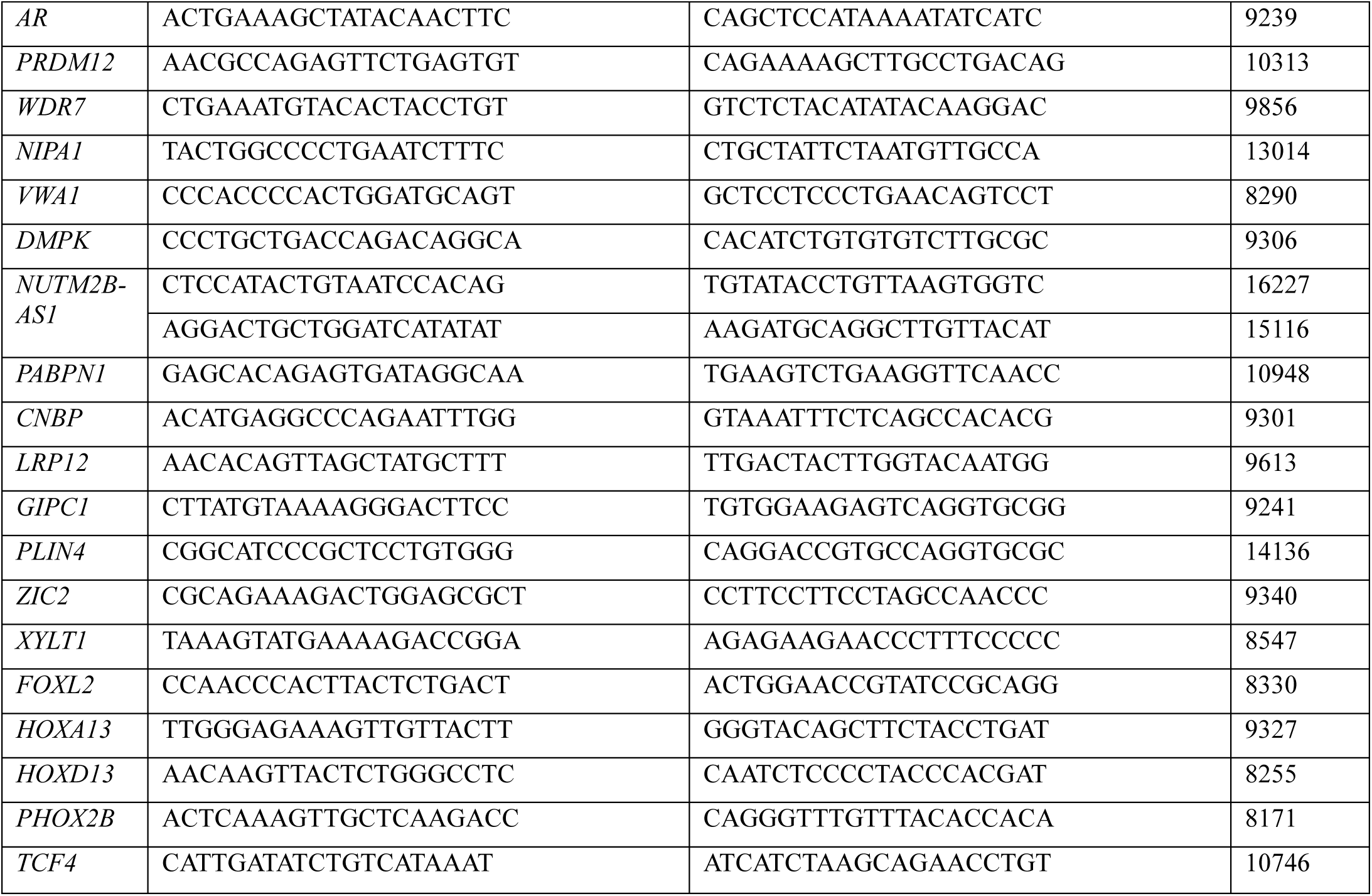
Selected crRNAs for nCATS determined through in vitro cleavage.

**Table S5.**
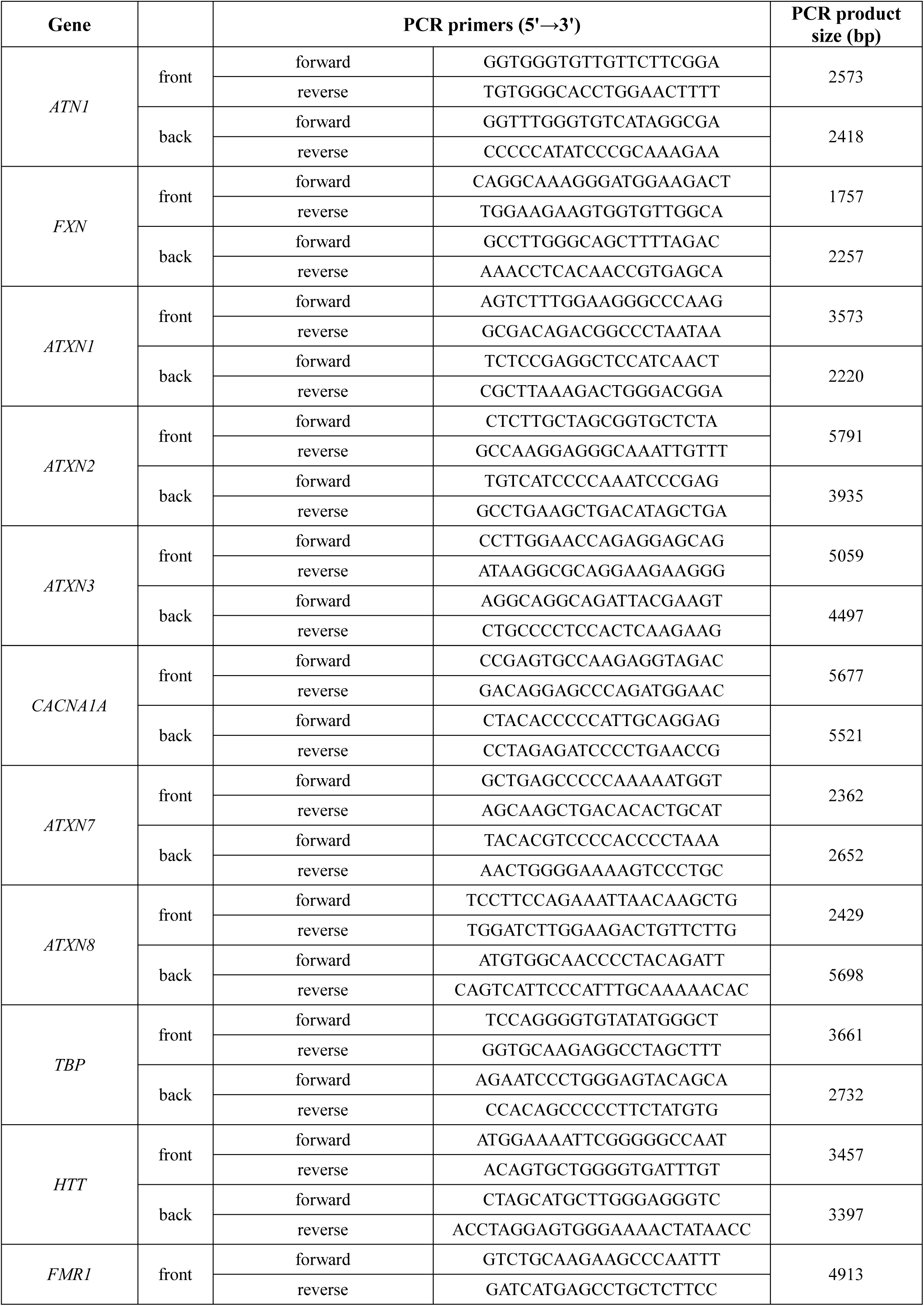
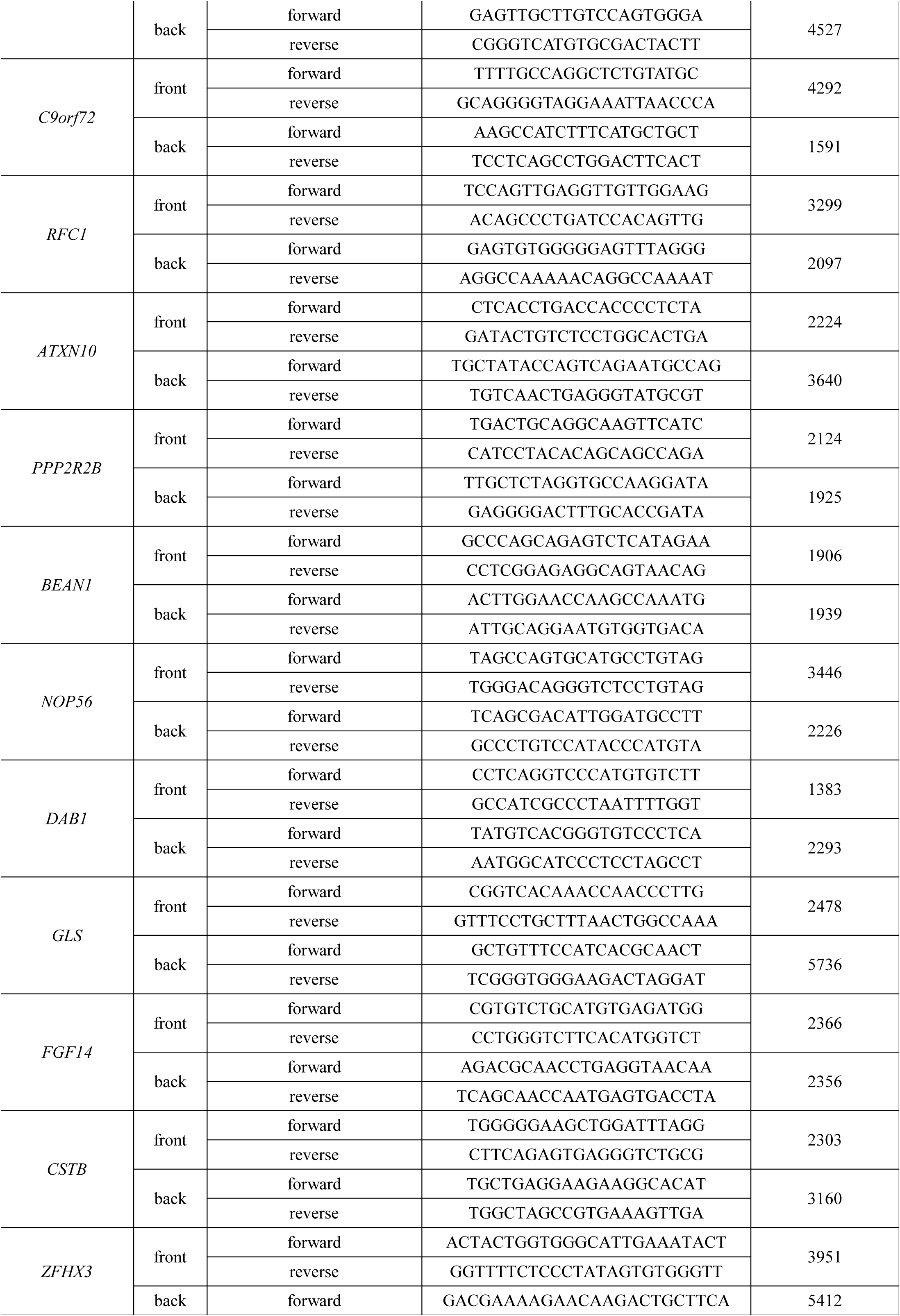
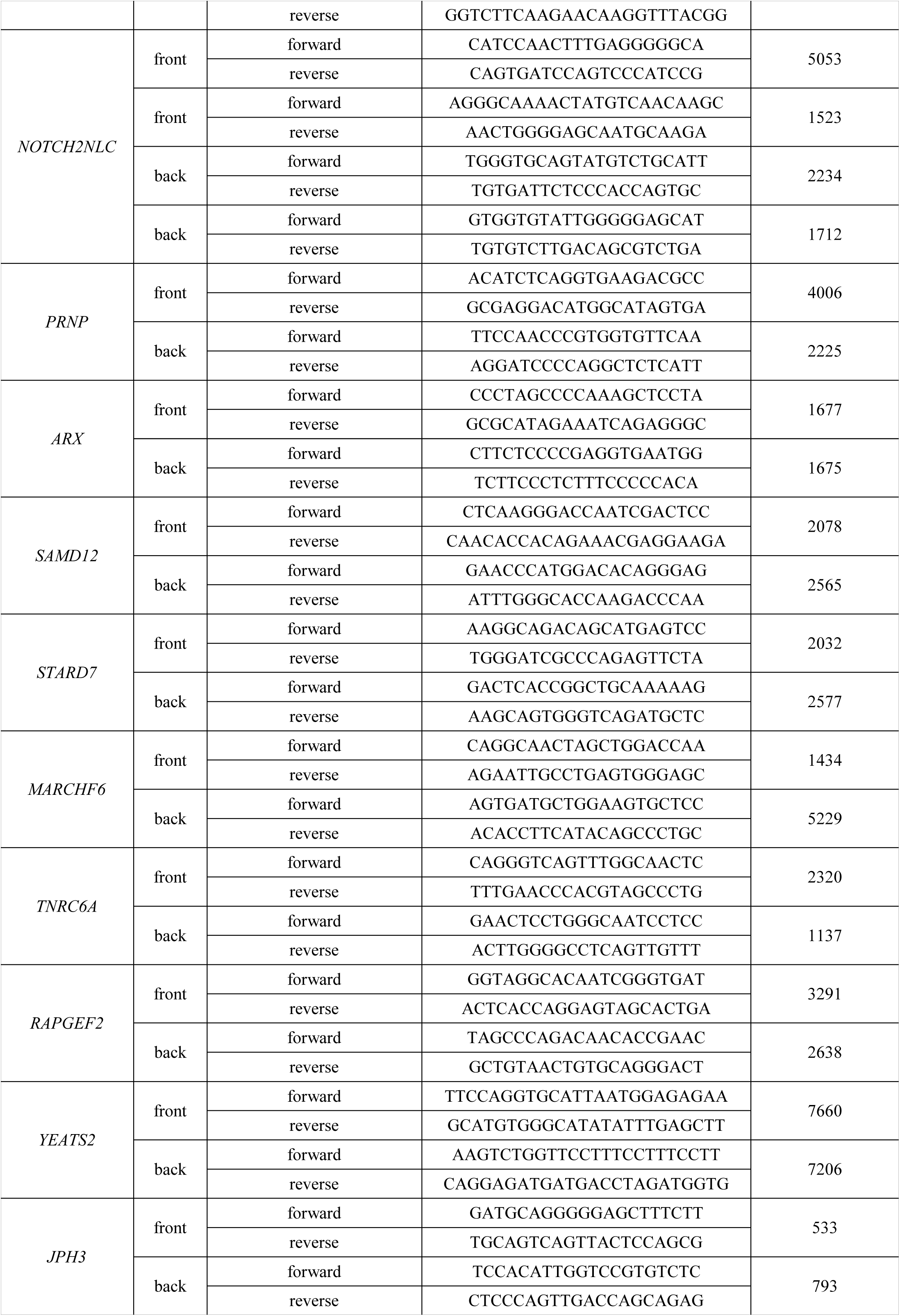
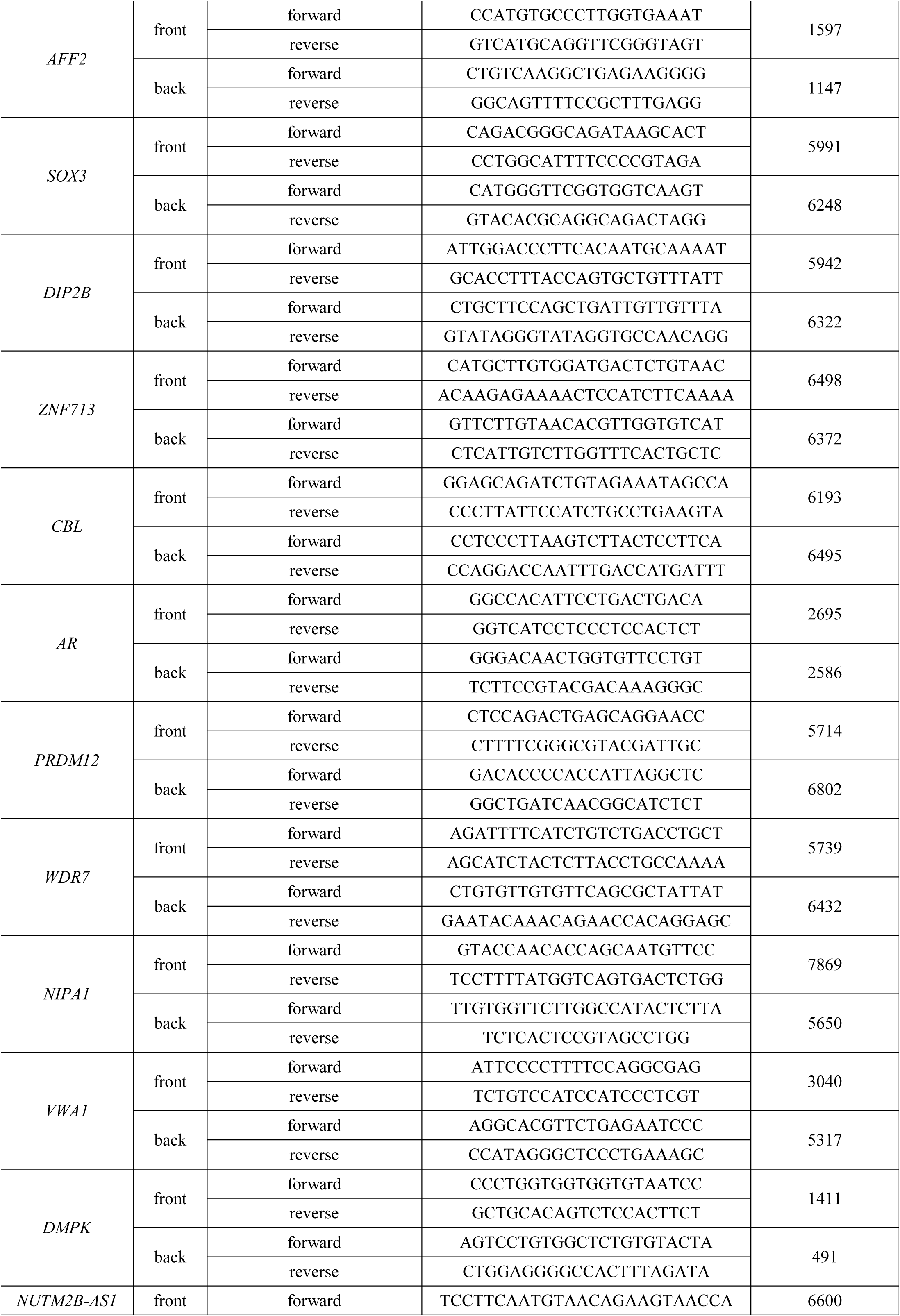
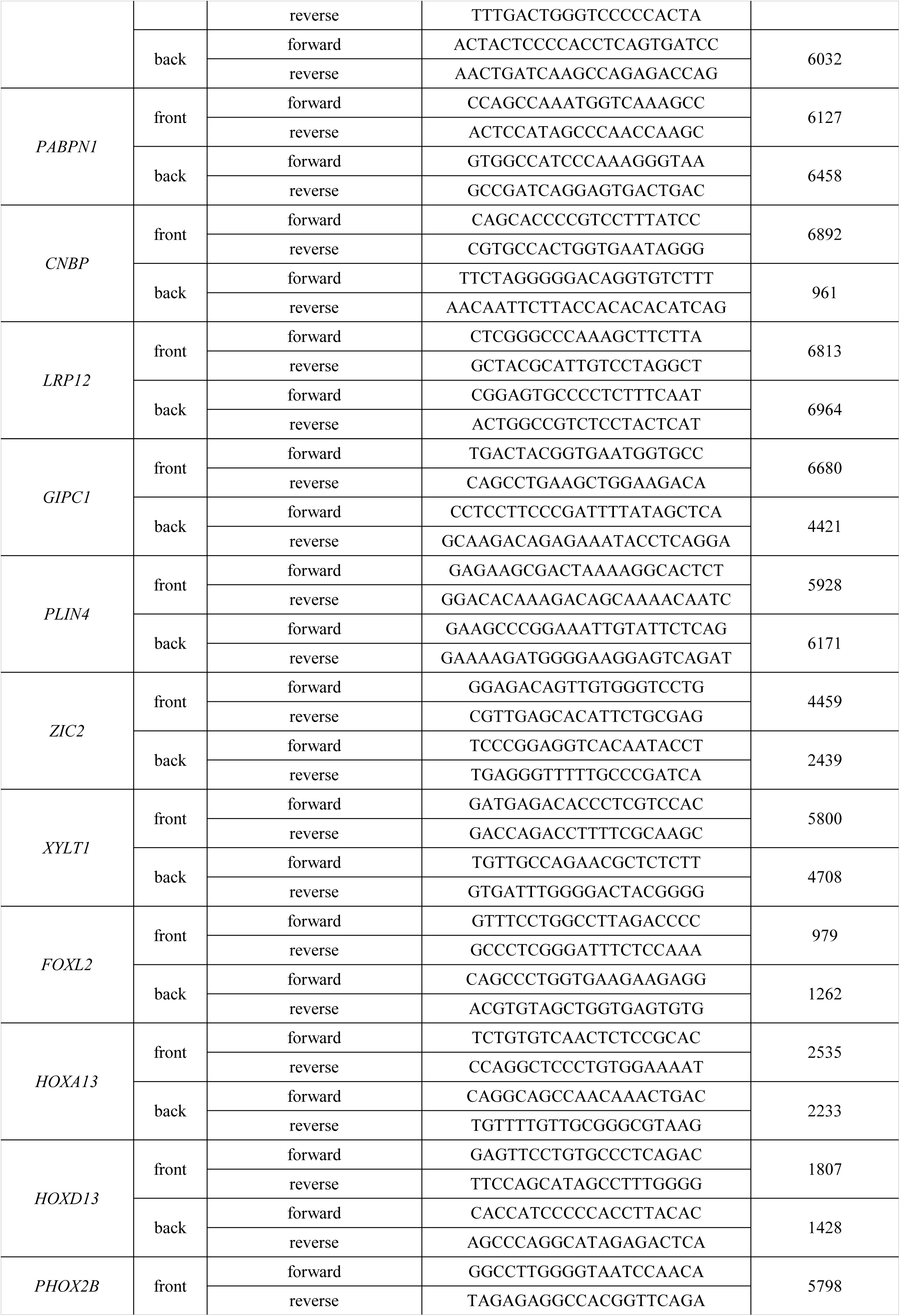
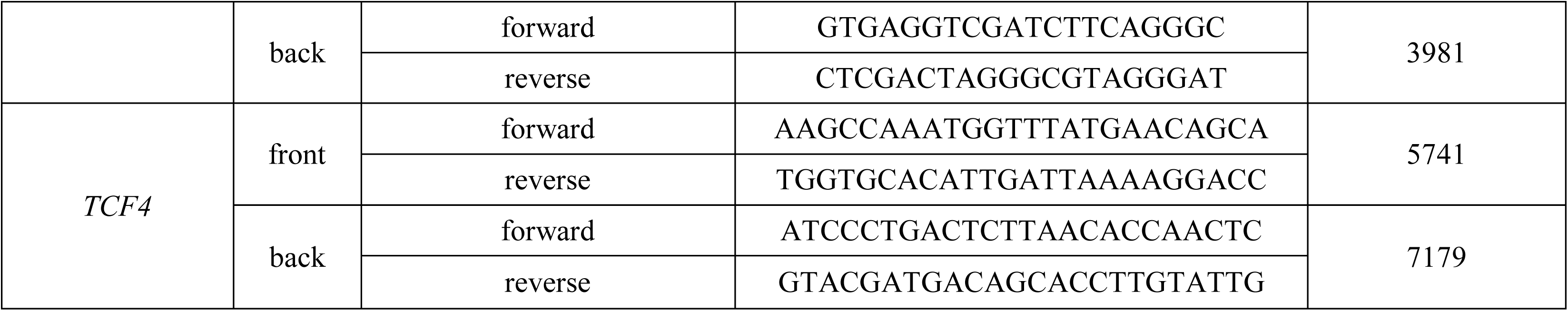
PCR primers targeting DNA fragments for in vitro cleavage assays.

**Table S6.**
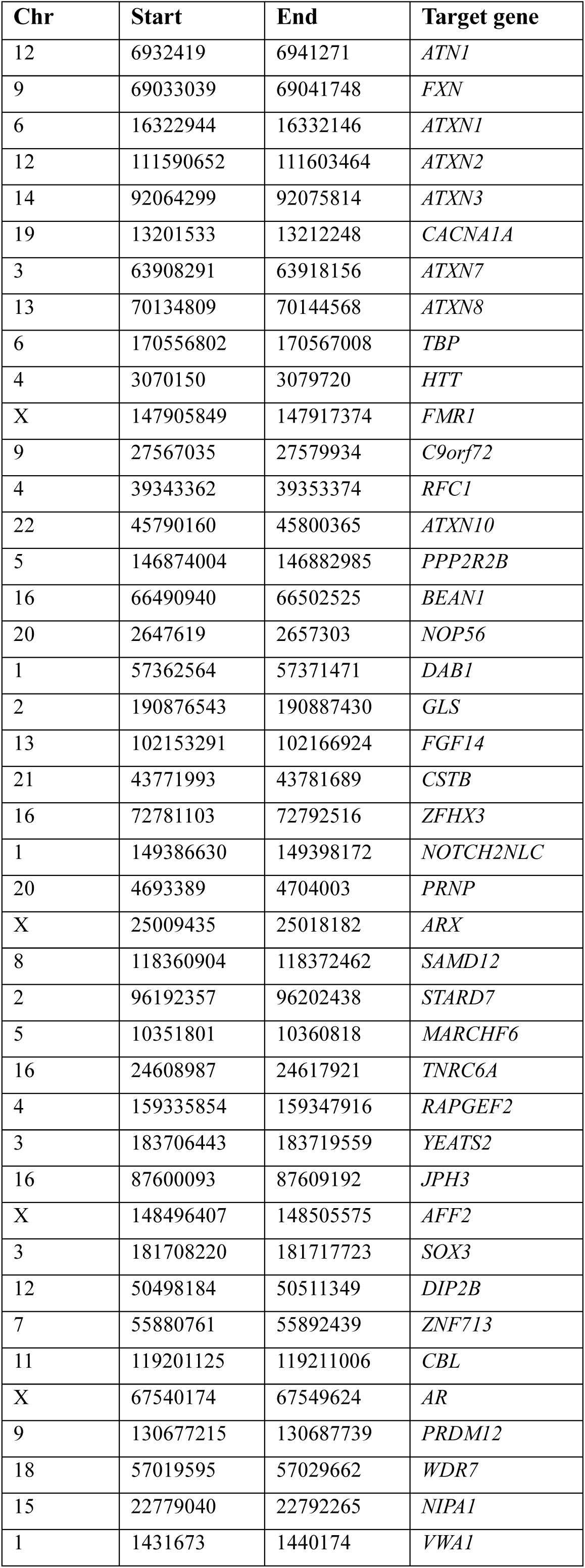
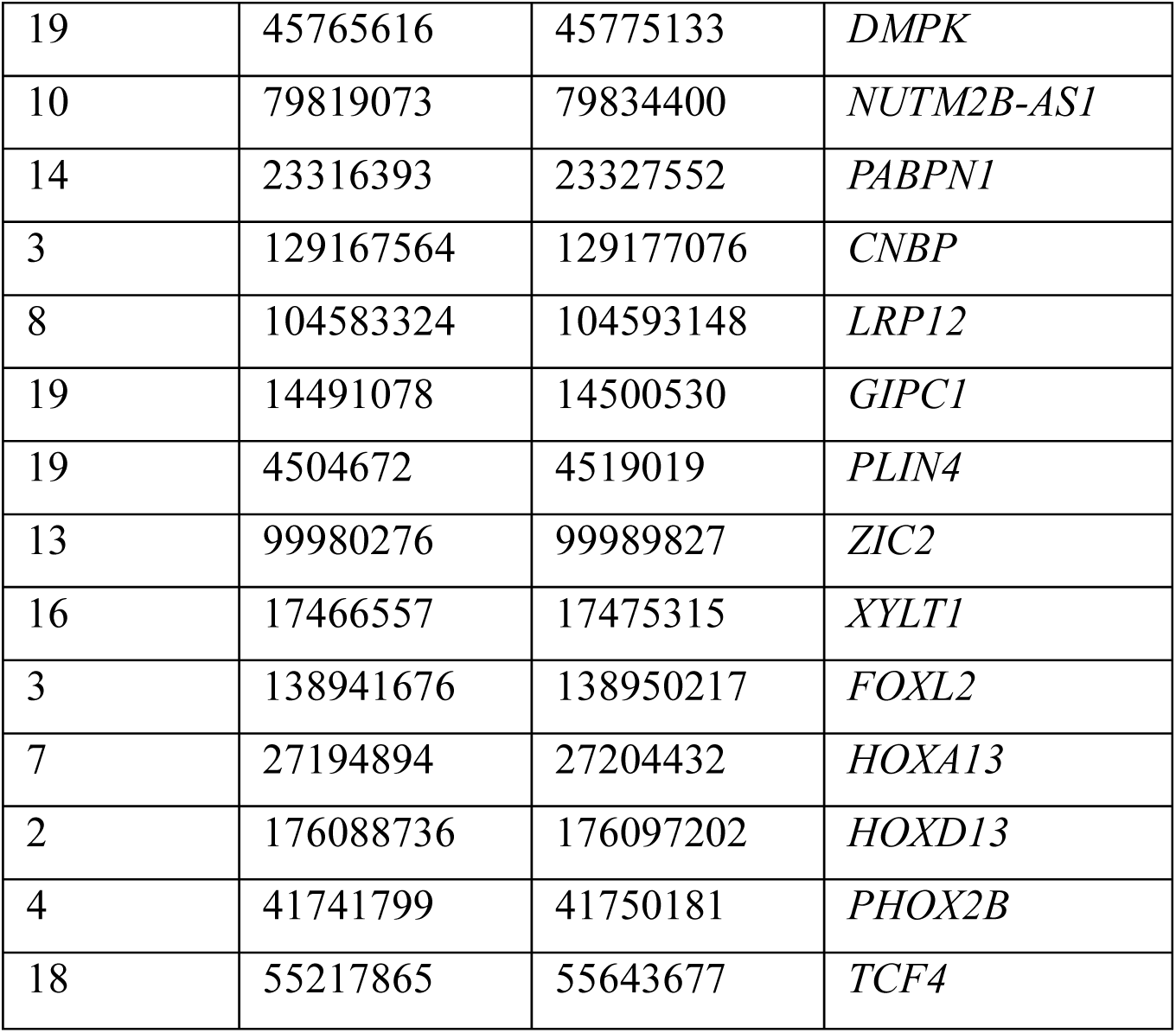
Adaptive sampling BED file.

**Table S7.**
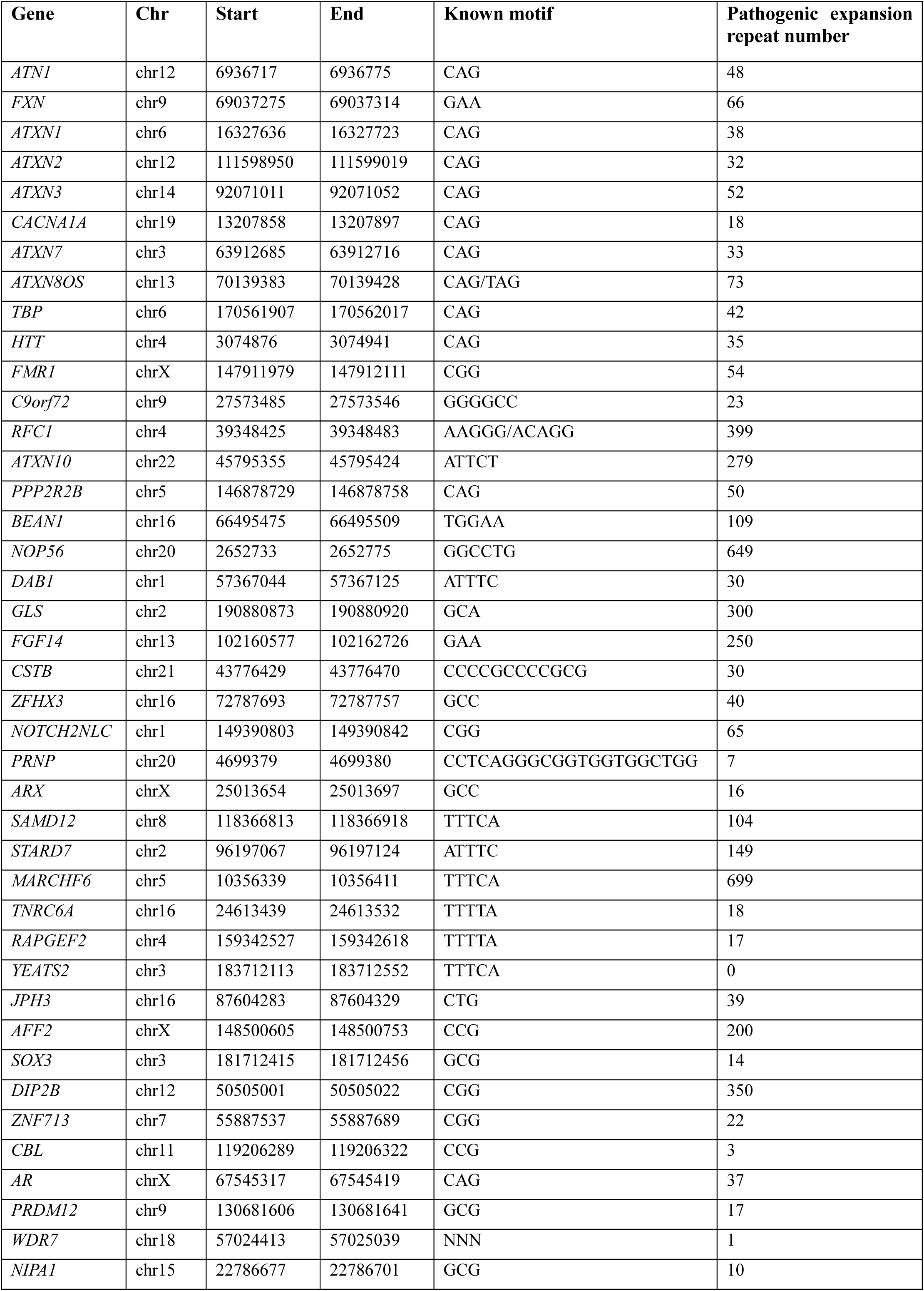
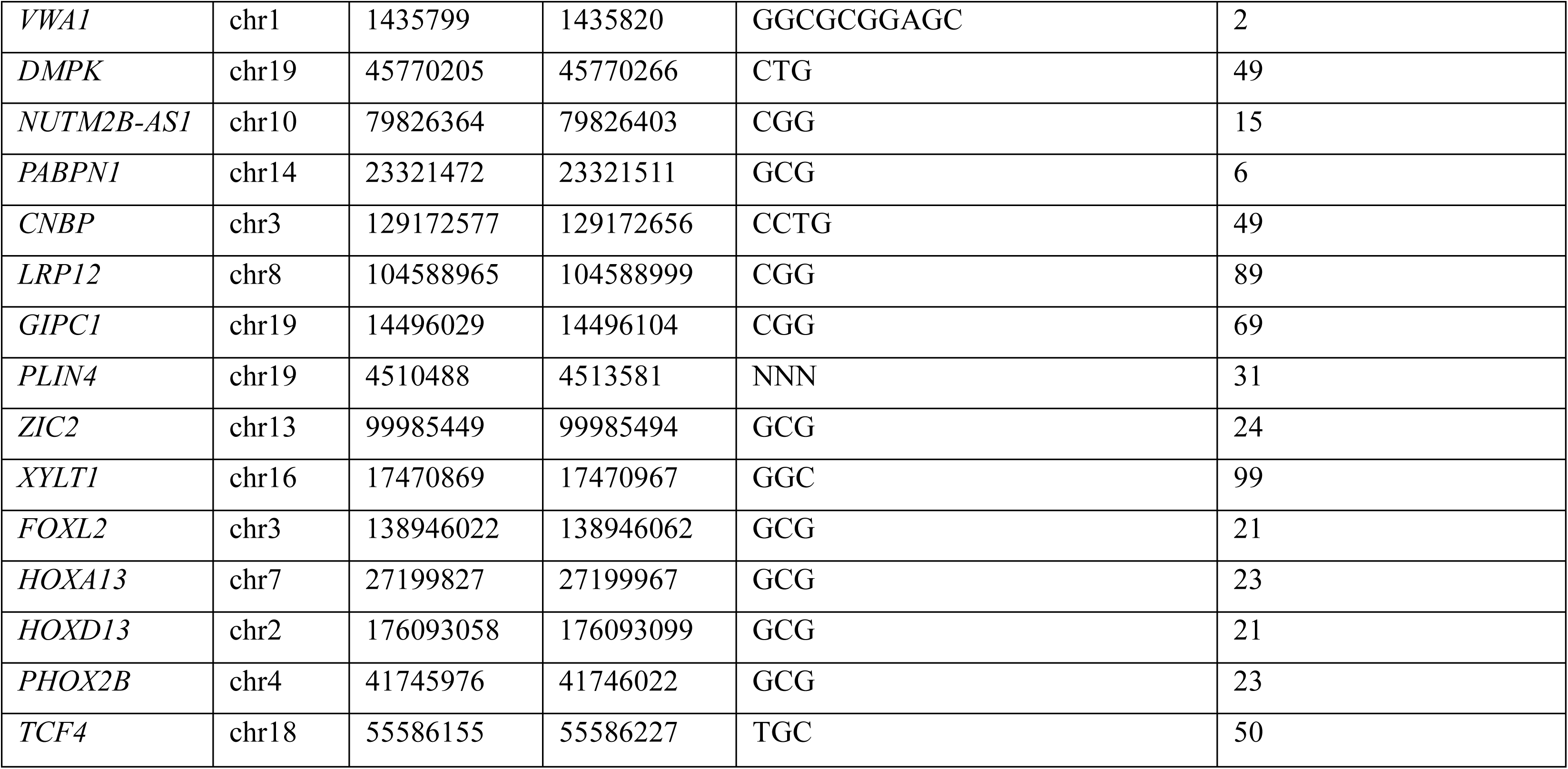
Known STR disease repeat expansion information.

**Table S8.**
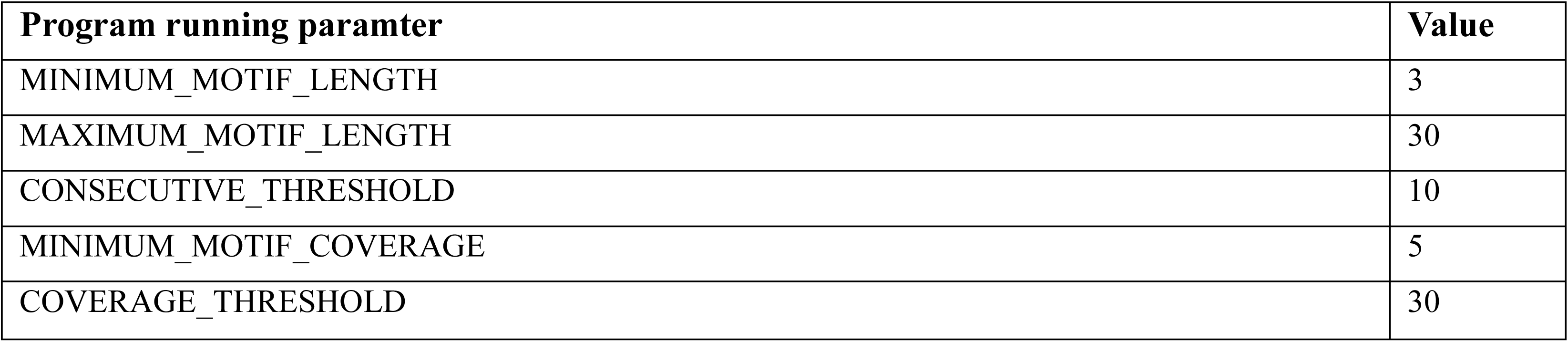
STRiker configuration parameters used in benchmark analysis.

**Table S9.**
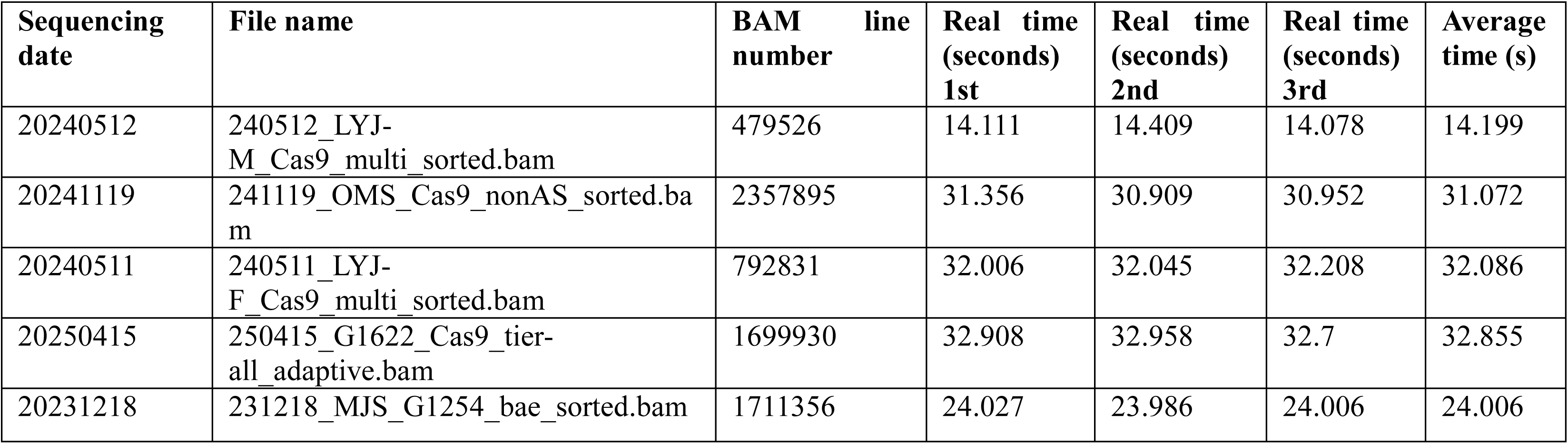
STRiker computational performance benchmarking.

