## Supplementary Materials for "Cas9-enriched nanopore sequencing enables comprehensive and multiplexed detection of repeat expansions"

**List of Supplementary Materials:**

Fig S1 to S6 for multiple supplementary figures

Table S1 to S9 for multiple supplementary tables

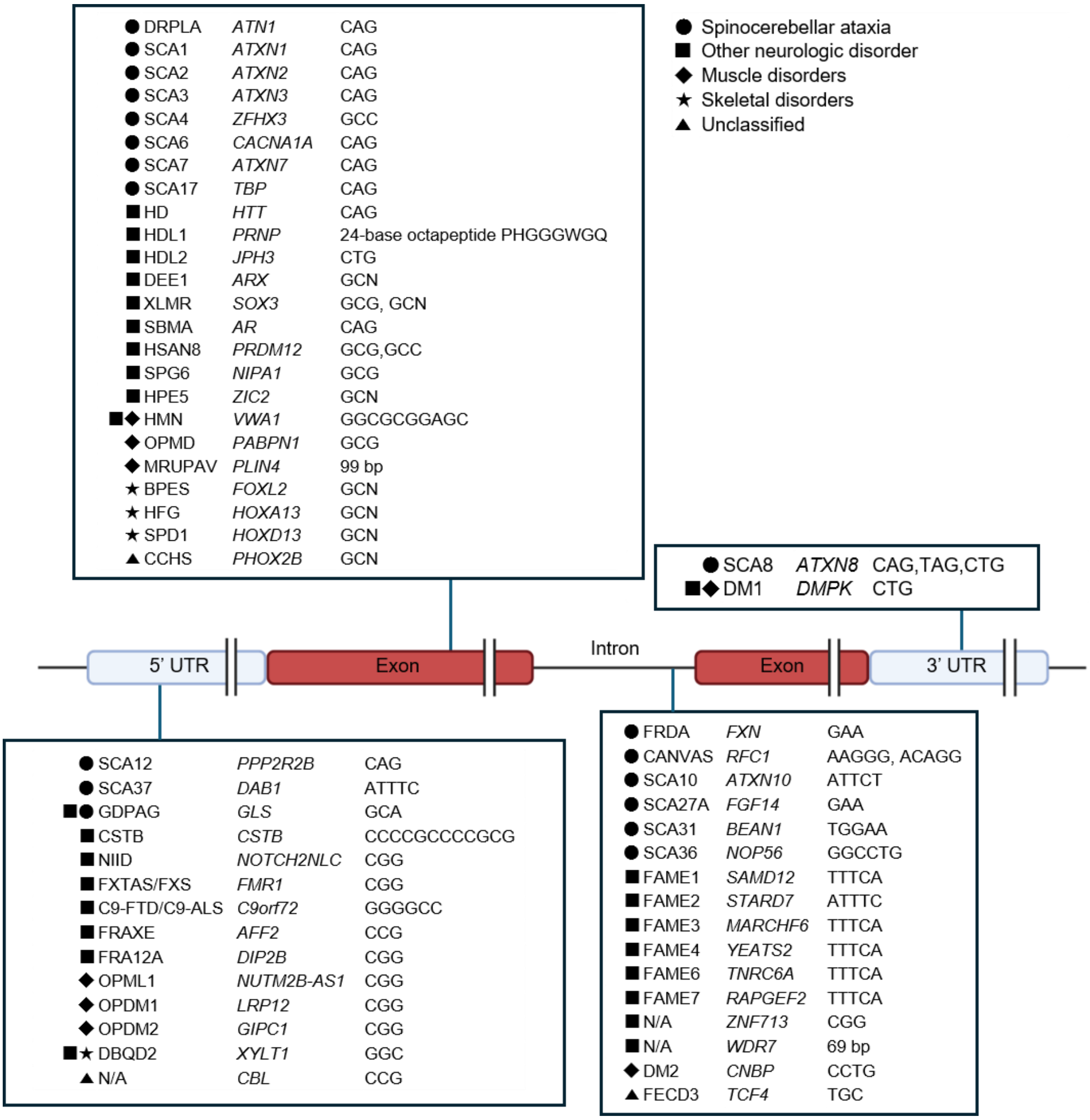


**Fig. S1. Overview of STR loci included in the 56-gene panel.**

Schematic representation of disease-associated short tandem repeat (STR) loci categorized by their genomic location (5′ untranslated region (UTR), exon, intron, or 3′ UTR) and repeat motif type. Each box lists representative disorders, associated genes, and repeat motifs. Symbols indicate the major clinical categories. This panel comprises 56 known pathogenic or candidate STR loci encompassing a wide range of repeat motifs implicated in neurological, muscular, and developmental disorders. Created with BioRender.com

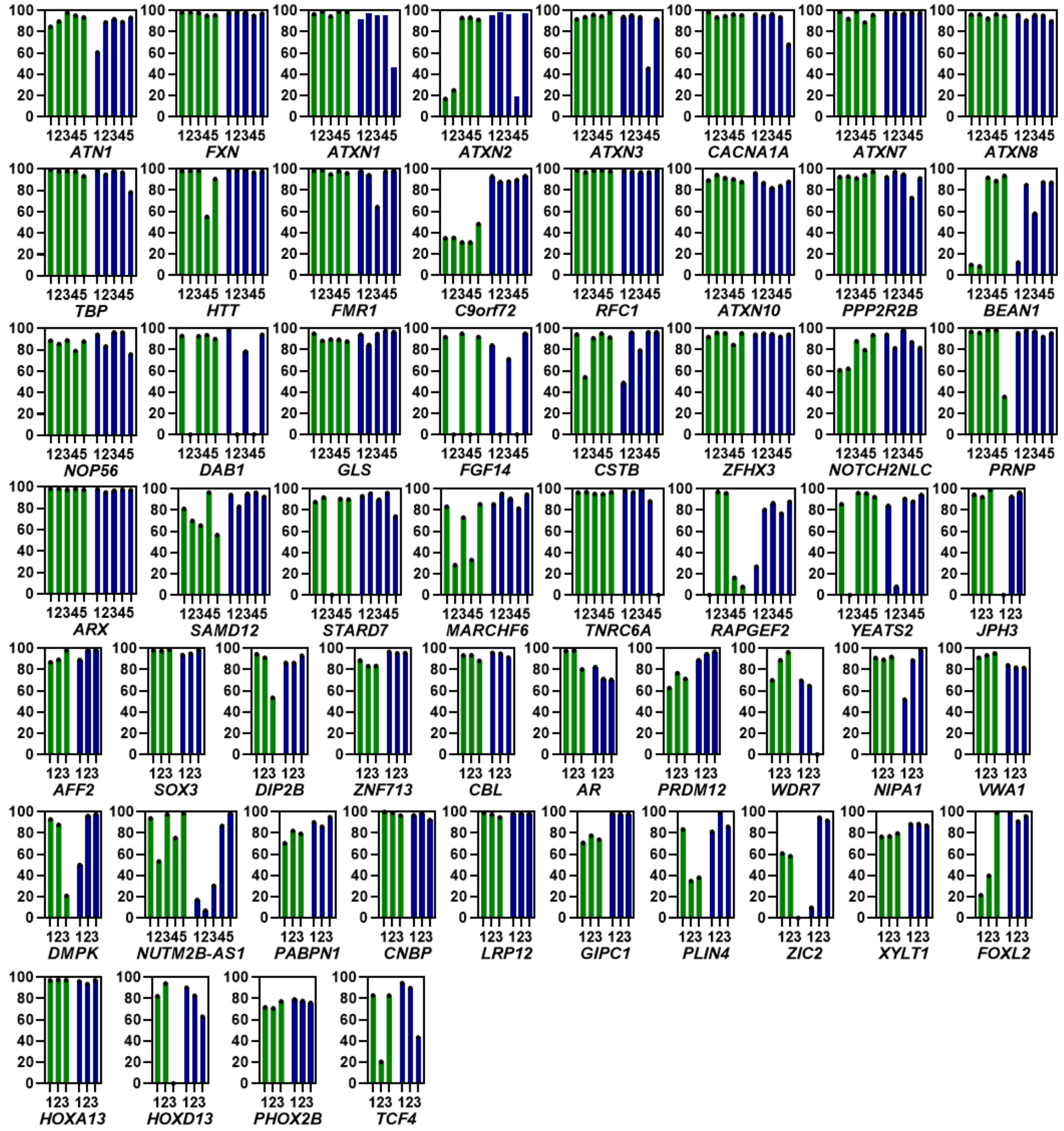


**Fig. S2. sgRNA screening for high cleavage efficiency via *in vitro* cleavage assay.**

Cleavage efficiency of sgRNAs targeting upstream and downstream regions of repeat expansion disease-associated gene regions of interest (ROIs), assessed by *in vitro* cleavage assay. Three or five different sgRNAs were tested for each region.

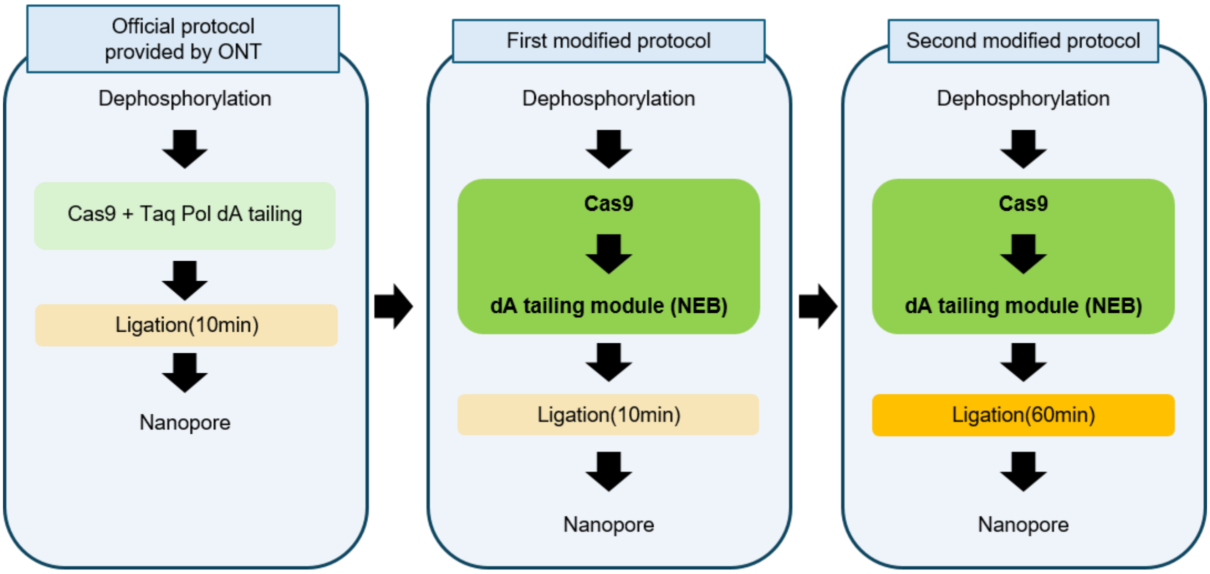


**Fig. S3. Schematic diagram with a comparison of the optimized nCATS protocol with the conventional protocol.**

The optimized protocol employs sequential Cas9 cleavage at both ROI termini, followed by bead purification, rather than simultaneous Taq polymerase and Cas9 treatment. Further optimization includes dA-tailing using a Klenow fragment (exo-) (NEB) and an extended nanopore adapter ligation time to 60 minutes.

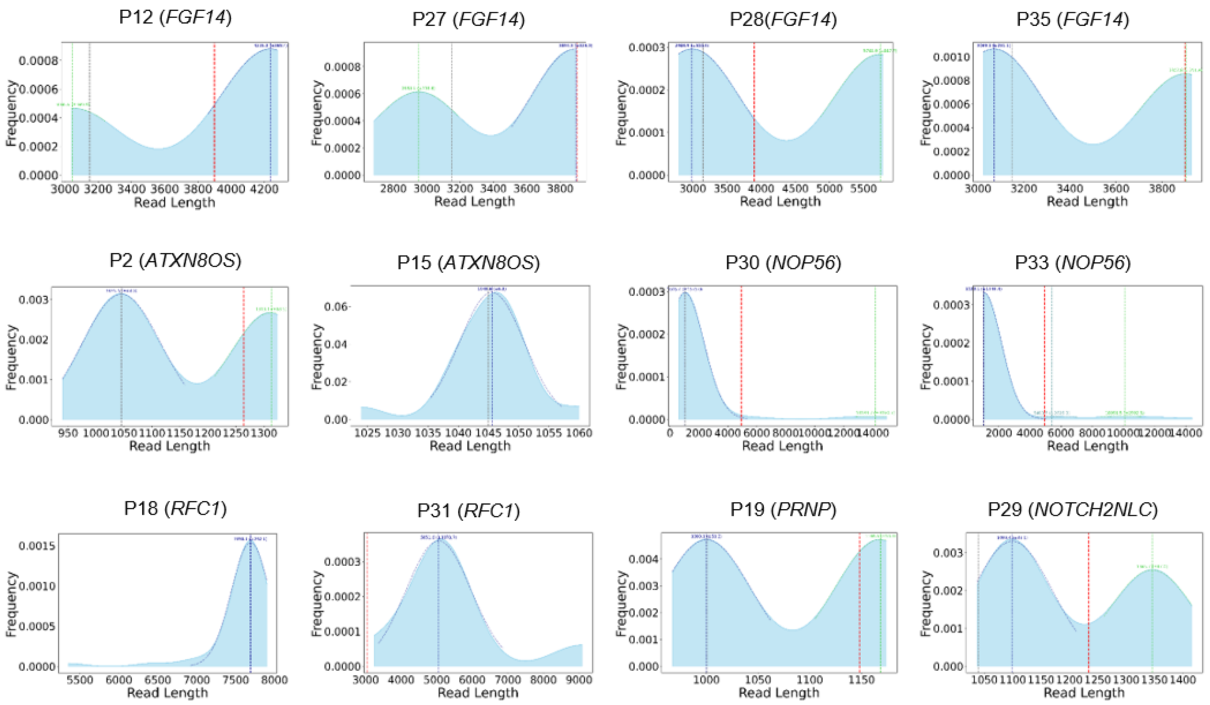


**Fig. S4. KDE read length distributions at expansion sites for newly diagnosed patients using the STRiker pipeline.**

Read lengths span the repeat locus plus 500 bp flanking regions on each side. Gaussian-fitted peaks are shown in blue (primary) and green (secondary). The red line indicates the threshold for pathogenic expansion.

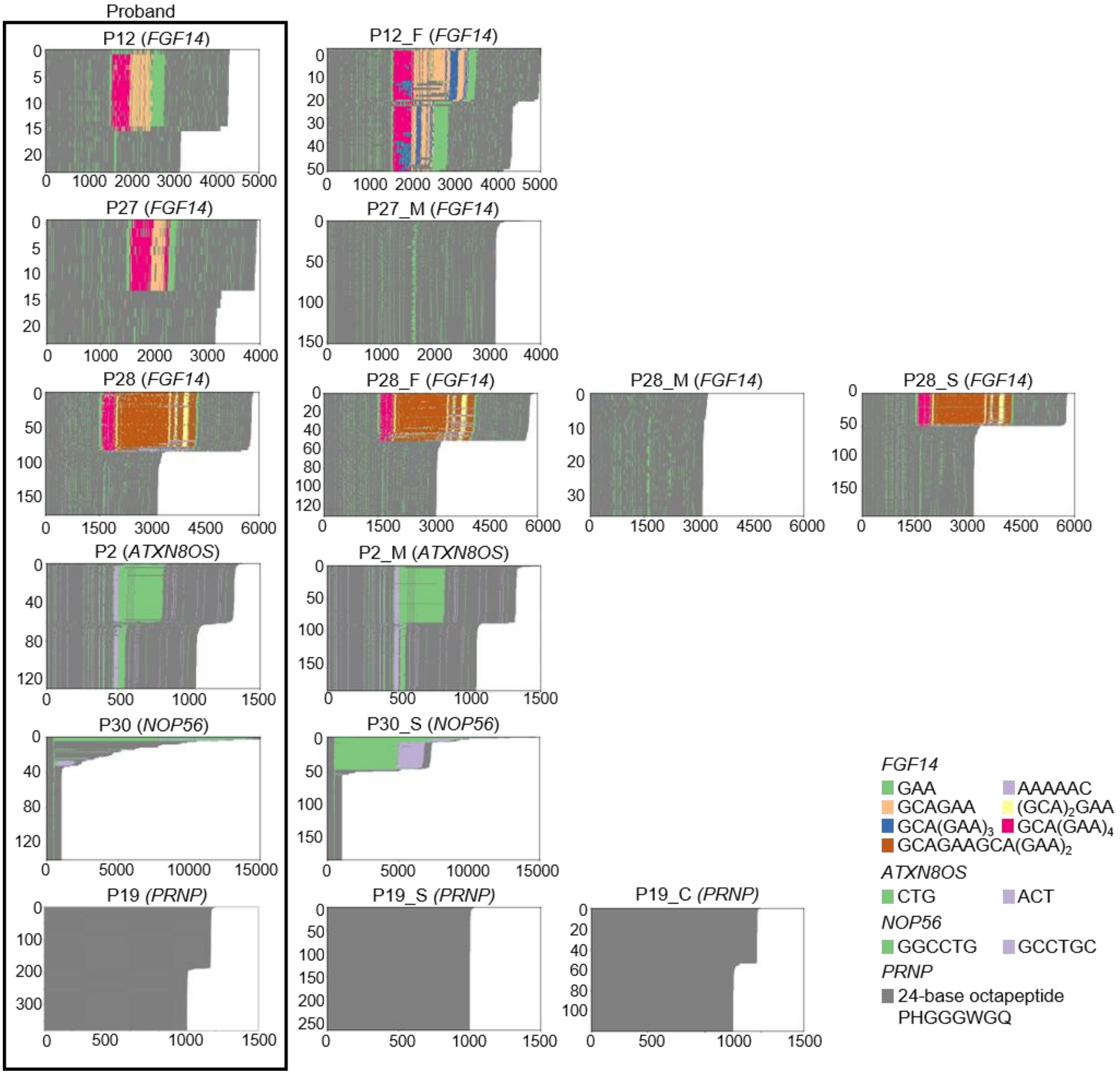


**Fig. S5. Repeat motif composition analysis in family members identified through cascade screening.**

Heatmaps of the repeat motif composition for newly diagnosed patients and their relatives identified by cascade screening. Each panel represents long-read alignments at disease-associated loci, with distinct colors indicating different motif types for each gene. *PRNP* repeat expansions consist of an octapeptide motif of 24 bp (PHGGGWGQ), for which no distinct motif colors are applied.

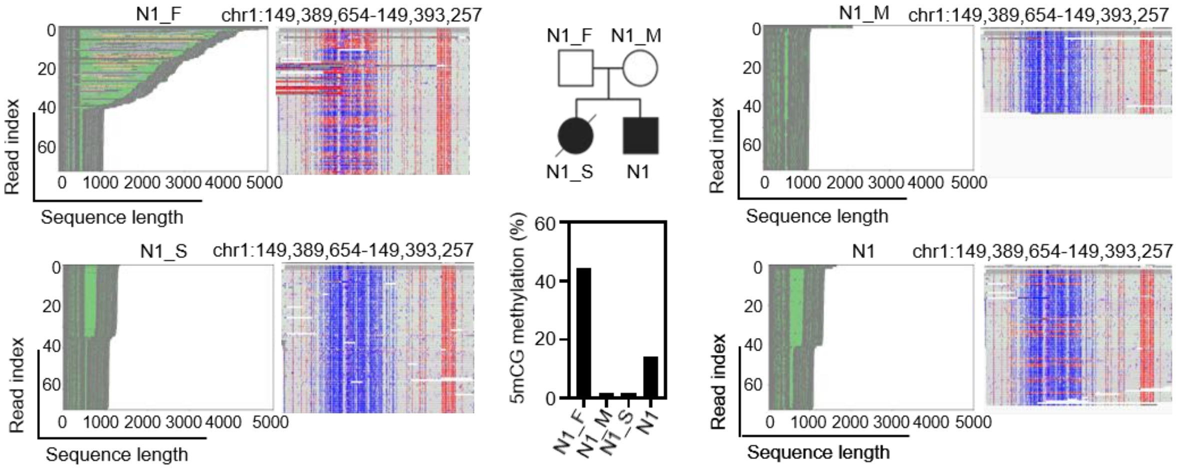


**Fig. S6. Case of contraction during paternal inheritance in a family with *NOTCH2NLC*-related neuronal intranuclear inclusion disease (NIID).** Long-read alignment and methylation profiles of each family member (N1_F, N1_M, N1, and N1_S). The father (N1_M) carried a long GGC repeat with hypermethylation and was asymptomatic, whereas the patient (N1) inherited a shorter allele with hypomethylation, which is likely associated with the development of neuronal intranuclear inclusion disease (NIID). The older sister (N1_S) of the patient also carried a hypomethylated expanded allele, was clinically affected, and died. Created with BioRender.com

**Table S1. Patient demographics and clinical characteristics.**

| **Characteristics** | **N (%)** |
|---|---|
| Total patients | 37 |
| Gender | |
| - Male | 18 (48.6) |
| - Female | 19 (51.4) |
| Age (yr), mean ± SD | 43.7 ± 14.2 |
| Family history | |
| - Yes | 14 (37.8) |
| - No | 23 (62.2) |
| Onset age (yr), mean ± SD | 35.5 ± 17.5 |
| Cerebellar atrophy | |
| - Yes | 30 (88.2) |
| - No | 4 (11.8) |
| Other brain abnormality | |
| - Yes | 18 (52.9) |
| - No | 16 (47.1) |
| Cognitive dysfunction | |
| - Yes | 6 (16.2) |
| - No | 31 (83.8) |
| Epilepsy | |
| - Yes | 1 (2.7) |
| - No | 36 (97.3) |
| Diagnosed | |
| - Yes | 12 (32.4) |
| - No | 25 (67.6) |
| Diagnosed gene | |
| - *FGF14* | 4 (10.8) |
| - *ATXN8OS* | 2 (5.4) |
| - *RFC1* | 2 (5.4) |
| - *NOP56* | 2 (5.4) |
| - *NOTCH2NLC* | 1 (2.7) |
| - *PRNP* | 1 (2.7) |
| - Undiagnosed | 25 (67.6) |

SD, standard deviation

**Table S2. Clinical and genetic information of study participants.**

| ID | Age at last follow-up (yr) | Gender | Family history | Affected family members | Onset age (yr) | Neurologic symptom | Gait at last follow-up | Cerebellar atrophy | Other brain abnormality | Cognitive dysfunction | Epilepsy | Previous genetic test | Diagnosed gene |
|---|---|---|---|---|---|---|---|---|---|---|---|---|---|
| Patient 1 | 42 | Male | No | NA | 40 | Cognitive dysfunction, dysarthria, ataxia | Impossible | Yes | Diffuse brain atrophy | Yes | No | WES, gene panel | NA |
| Patient 2 | 44 | Female | Yes | Mother | 30 | Dysarthria, ataxia | Possible (SARA gait 1) | Yes | No | No | No | Gene panel, SCA1,2,3,6,7,17, DRPLA, FRDA | ATXN8OS |
| Patient 2_M | NA | Female | NA | NA | NA | NA | NA | NA | NA | NA | NA | NA | ATXN8OS |
| Patient 3 | 59 | Female | Yes | Sister | 56 | Ataxia, dysarthria, dysphagia | Possible (ataxic) | Yes | No | No | No | WES, SCA1,2,3,6,7,8,17 | NA |
| Patient 4 | 39 | Male | No | NA | 29 | Dysarthria, ataxia, but with symptom fluctuation | Possible (ataxic) | Yes | No | No | No | WES, gene panel, SCA1,2,3,6,7,17, DRPLA, FRDA | NA |
| Patient 5 | 43 | Male | No | NA | 37 | Ataxia, dizziness, dysarthria, dysphagia, neurogenic bladder, expired | Impossible (since 42yr) | Yes | Severe atrophy of the pons and bilateral cerebellums with: 1) T2 HSI in the bilateral middle cerebellar peduncles, 2) a cruciform T2 HSI in the pons | No | No | WES, gene panel, SCA1,2,3,6,7,8,17, DRPLA, C9orf72 | NA |
| Patient 6 | 51 | Female | Yes | Father, sister | 45 | Ataxia | Possible (SARA gait 2) | NA | NA | No | No | WGS, WES, SCA1,2,3,6,7,17, DRPLA, FRDA | NA |
| Patient 7 | 55 | Male | No | NA | 53 | Ataxia, spasticity | Possible | Yes | 1) Atrophic change of the pons and cerebellum. 2) T2 HSI in the bilateral cerebral white matter, suggestive of small vessel disease | No | No | WES, SCA1,2,3,6,7,17, DRPLA, FRDA | NA |

| ID | Age at last follow-up (yr) | Gender | Family history | Affected family members | Onset age (yr) | Neurologic symptom | Gait at last follow-up | Cerebellar atrophy | Other brain abnormality | Cognitive dysfunction | Epilepsy | Previous genetic test | Diagnosed gene |
|---|---|---|---|---|---|---|---|---|---|---|---|---|---|
| Patient 8 | 41 | Female | No | NA | 29 | Ataxia, dysarthria | Possible (SARA gait 2) | Yes | No | No | No | Gene panel, SCA1,2,3,6,7,8,17, DRPLA, FRDA | NA |
| Patient 9 | 69 | Female | No | NA | 68 | Ataxia, gait disturbance | Possible | Yes | No | No | No | WES, SCA1,2,3,6,7,8,17, DRPLA | NA |
| Patient 10 | 31 | Male | No | NA | 1 | Global developmental delay, intellectual disability, ataxia | Possible | Yes | No | Yes | No | WES, gene panel, SCA1,2,3,6,7,8,17, DRPLA | NA |
| Patient 11 | 48 | Female | Yes | Father, brother, sister | 40 | Ataxia, gait disturbance | Possible | Yes | No | No | No | WES, gene panel, SCA1,2,3,6,7,8,17, DRPLA | NA |
| Patient 12 | 29 | Male | Yes | Father, grandfather | 25 | Ataxia, gait disturbance, dysarthria | Possible | Yes | No | No | No | WES, gene panel, SCA1,2,3,6,7,8,17, C9orf72 | FGF14 |
| Patient 12_F | 61 | Male | NA | NA | 35 | Ataxia, gait disturbance | Possible (ataxic) | Yes | No | No | No | NA | FGF14 |
| Patient 13 | 53 | Female | No | NA | 50 | Ataxia, gait disturbance, dysarthria | Possible | Yes | Diffuse volume decrease at bilateral cerebellum, pons, and medulla | No | No | WES, gene panel, SCA1,2,3,6,7,17, DRPLA | NA |
| Patient 14 | 25 | Female | Yes | Mother | 21 | Ataxia, gait disturbance, dysarthria | Impossible | Yes | No | No | No | Gene panel, SCA1,2,3,6,7,17, DRPLA | NA |
| Patient 15 | 28 | Female | No | NA | 21 | Ataxia, dizziness | Possible (SARA gait 2) | Yes | No | No | No | WES, gene panel, SCA1,2,3,6,7,8,17, DRPLA | ATXN8OS |
| Patient 16 | 61 | Male | No | NA | 35 | Ataxia, gait disturbance, dysarthria | Impossible | Yes | Diffuse brain atrophy | No | No | WGS, WES, SCA1,2,3,6,7,8,17 | NA |
| Patient 17 | 35 | Male | No | NA | 13 | Ataxia, dysarthria | Impossible | Yes | Diffuse brain atrophy | Yes | No | WES, gene panel SCA1,2,3,6,7,8,17, DRPLA | NA |
| Patient 18 | 64 | Male | No | NA | 57 | Ataxia, gait disturbance, scanning speech | Possible (ataxic) | Yes | No | No | No | WES, gene panel SCA1,2,3,6,7,8,17, DRPLA | RFC1 |

| ID | Age at last follow-up (yr) | Gender | Family history | Affected family members | Onset age (yr) | Neurologic symptom | Gait at last follow-up | Cerebellar atrophy | Other brain abnormality | Cognitive dysfunction | Epilepsy | Previous genetic test | Diagnosed gene |
|---|---|---|---|---|---|---|---|---|---|---|---|---|---|
| Patient 19 | 35 | Female | Yes | Father, aunt | 30 | Ataxia, dysarthria, dizziness | Possible (ataxic) | Yes | Diffuse brain atrophy | Yes | No | WES, gene panel SCA1,2,3,6,7,8,17, DRPLA | PRNP |
| Patient 19_S | 34 | Male | NA | NA | NA | NA | NA | NA | NA | NA | NA | NA | NA |
| Patient 19_C | 28 | Male | NA | NA | NA | NA | NA | No | No | No | No | NA | PRNP |
| Patient 20 | 18 | Male | No | NA | 14 | Ataxia, gait disturbance, dysarthria, dysphagia | Possible (ataxic) | Yes | No | No | No | WES, gene panel SCA1,2,3,6,7,8,17, DRPLA, FRDA | NA |
| Patient 21 | 30 | Female | No | NA | 28 | Ataxia, gait disturbance, dysarthria, dysphagia, bradykinesia | Possible (ataxic) | Yes | Chiari malformation 1 with syringohydromyelia, mild midbrain atrophy | Yes | No | Gene panel SCA1,2,3,6,7,8,17, DRPLA | NA |
| Patient 23 | 64 | Male | No | NA | 60 | Ataxia, dysarthria | Possible | Yes | Diffuse brain atrophy | No | No | Gene panel SCA1,2,3,6,7,8,17, DRPLA | NA |
| Patient 24 | 61 | Female | No | NA | 56 | Dysarthria, ataxia, scanning speech, gait disturbance | Possible (ataxic) | Yes | Atrophic change in the bilateral cerebellar and brain stem | No | No | WES, gene panel SCA1,2,3,6,7,8,17, DRPLA | NA |
| Patient 25 | 44 | Male | No | NA | 39 | Dysarthria, ataxia, gait disturbance | Possible (ataxic) | Yes | 1) Underlying atrophic change in the brain parenchyma. 2) Mild degree T2 HSI lesions around bilateral lateral ventricles (posterior horns) | No | No | Gene panel SCA1,2,3,6,7,8,17, DRPLA | NA |
| Patient 26 | 39 | Male | Yes | Mother | 31 | Dysarthria, ataxia | Possible | Yes | No | No | No | Gene panel SCA1,2,3,6,7,8,17, DRPLA | NA |
| Patient 27 | 33 | Male | No | NA | 1 | Developmental delay, dysarthria, ataxia | Possible | Yes | No | No | No | WES, gene panel SCA1,2,3,6,7,8,17, DRPLA, FRDA, HD | FGF14 |
| Patient 27_M | NA | Female | NA | NA | NA | NA | NA | NA | NA | NA | NA | NA | NA |

| ID | Age at last follow-up (yr) | Gender | Family history | Affected family members | Onset age (yr) | Neurologic symptom | Gait at last follow-up | Cerebellar atrophy | Other brain abnormality | Cognitive dysfunction | Epilepsy | Previous genetic test | Diagnosed gene |
|---|---|---|---|---|---|---|---|---|---|---|---|---|---|
| Patient 28 | 26 | Female | Yes | Sister | 7 | Gait disturbance, ataxia, dystonia, spasticity | Possible (ataxic) | No | Bilateral T2 HIS at globus pallidus | No | No | LRS, WGS, WES, gene panel | FGF14 |
| Patient 28_S | 24 | Female | NA | NA | 3 | Periodic paralysis, dystonia, ataxia, spasticity | Possible (ataxic) | No | Sublte T2 signal abnormalities in the both globus pallidus | No | No | LRS, WGS, WES, gene panel | FGF14 |
| Patient 28_M | NA | Female | NA | NA | NA | NA | NA | NA | NA | NA | NA | NA | NA |
| Patient 28_F | NA | Male | NA | NA | NA | NA | NA | NA | NA | NA | NA | NA | FGF14 |
| Patient 29 | 61 | Female | No | NA | 60 | Transient cognitive impairment, ataxia | Possible | No | Bilateral confluent T2 HSI on white matter, positive zigzag edging sign | No | No | NOTCH2NLC | NOTCH2NLC |
| Patient 30 | 49 | Male | Yes | Father, sister | 45 | Dysarthria, ataxia | Possible | NA | NA | No | No | Gene panel, SCA1,2,3,6,7,8,17, DRPLA | NOP56 |
| Patient 30_S | 52 | Female | NA | NA | 44 | Abnormal sensation of lower limb, dysarthria, gait disturbance, ataxia | Possible (ataxic) | NA | NA | No | No | NA | NOP56 |
| Patient 31 | 62 | Female | No | NA | 55 | Gait disturbance, ataxia, peripheral neuropathy | Possible (ataxic) | No | Suspicious two tiny nodular isointense lesions, adjacent to the course of the right anterior inferior cerebellar artery | No | No | None | RFC1 |
| Patient 32 | 17 | Male | No | NA | 12 | Gait disturbance, ataxia | Possible (ataxic) | Yes | Pachy areas of increased T2 signal in the cerebral deep white matter | No | No | WGS, WES, gene panel, SCA1,2,3,6,7,8,17, DRPLA | NA |

| ID | Age at last follow-up (yr) | Gender | Family history | Affected family members | Onset age (yr) | Neurologic symptom | Gait at last follow-up | Cerebellar atrophy | Other brain abnormality | Cognitive dysfunction | Epilepsy | Previous genetic test | Diagnosed gene |
|---|---|---|---|---|---|---|---|---|---|---|---|---|---|
| Patient 33 | 60 | Female | Yes | Aunt, uncle, cousin | 55 | Dizziness, gait disturbance, ataxia, mild scanning speech | Possible | Yes | No | No | No | WGS, WES, gene panel, SCA1,2,3,6,7,8,17, DRPLA, FRDA | NOP56 |
| Patient 34 | 47 | Male | Yes | Brother | 33 | Gait disturbance, dysarthria | Possible (ataxic) | Yes | No | No | No | WGS, WES, gene panel, SCA1,2,3,6,7,8,17, DRPLA | NA |
| Patient 35 | 26 | Female | Yes | Mother, aunt, uncle, grandmother | 25 | Dysarthria, ataxia, mild scanning speech | Possible | NA | NA | No | No | Gene panel, SCA1,2,3,6,7,8,17, DRPLA | FGF14 |
| Patient 36 | 46 | Female | No | NA | 43 | Gait disturbance, dizziness, weakness, movement disorder | Possible | No | A few microbleeds at right frontal and occipital lob | No | No | Gene panel, SCA1,2,3,6,7,8,17, DRPLA | NA |
| Patient 37 | 51 | Male | Yes | Father, grandfather | 47 | Gait disturbance, dysarthria, ataxia | Possible (ataxic) | Yes | 1) Atrophic change of medulla and cervical spinal cord. 2) Confluent periventricular white matter hyperintensity. 3) A large supracerebellar arachnoid cyst. | No | No | None | NA |

Patient IDs ending with M, F, S, and C denote relationships with the proband as follows: mother (M), father (F), sibling (S), and cousin (C).

NA, not available; WES, whole-exome sequencing; SARA, Scale for the Assessment and Rating of Ataxia; SCA, spinocerebellar ataxia; DRPLA, dentatorubral-pallidoluysian atrophy;

FRDA, Friedeich ataxia; HSI, high signal intensity; WGS, whole-genome sequencing; HD, Huntington disease; LRS, long-read sequencing

**Table S3. All crRNAs for nCATS determined through in vitro cleavage.**

| Gene | crRNA sequence |
|---|---|
| *ATN1*_forward | AGGAGGATAGGCTTAAAAAC |
| | ATAGACAGATACGGATGGAT |
| | AATGAATACAGCTGGCTATA |
| | GAAGAGCGCACACACATCAG |
| | GAGCTAGCAGCACTTAATTC |
| *ATN1*_reverse | TGGGTGAAGGGATCCAAGCC |
| | TGTCACCAGAAGGCAGGTTT |
| | CCGGGTAAGGTGAGACCCTG |
| | TCGTTCTCGTGCAGAGGGTG |
| | CTAAAACTCGTGCTTCTCTT |
| *FXN*_forward | ATATCAACTTTGCAGGGAGC |
| | CTTGACTACCTCCAAGGAAG |
| | CAACATCTTTGCCCACGCCA |
| | AGGCTAAAAGCAACCAAGAG |
| | *AATATCCTAATGGCAGAGCA* |
| *FXN*_reverse | *ATCAACCAAATGTCACCTAA* |
| | *TCAGGACCCATACCTCGCAG* |
| | AGGTGTCCCCTTGTCACCCC |
| | GCCTACCCCCTGGTAAGGAC |
| | AAGTCTCAGGCCCAGTGGTC |
| *ATXN1*_forward | GAACAGAAGAGGGCTGTAAG |
| | CTACCTGCTCGTCTGCACCC |
| | AGCACACTTGCCTTTCTTTG |
| | GACAAATCATTGGTTGCGTC |
| | GACTCTGAACCTGTCACTTG |
| *ATXN1*_reverse | ACCAGTCCAGTGAGTCTTCA |
| | TTGTGTCCTTGATTTCTCAT |
| | CTGTCTGCTCCGTGCTGCAC |
| | ATCAGTACTGAAAGCTTACG |
| | AGGAATGTGGTGCCATCCTG |
| *ATXN2*_forward | CCGCACTCATTGCAACCTCC |
| | TAACACGGCTGTAGCTATTT |
| | CCTTAGCTCATGAGGCTGGG |
| | TGAGCCCCACCTGTTAGATC |
| | AAACAAGCCTGTTAGGGGGC |
| *ATXN2*_reverse | AACTGAACGAGACAGGGTCT |
| | CACCCAACAATTTCGTTTGT |
| | ATCATTTGGGACCGCCATAG |
| | AGGATTGCTCGGGACCGCCT |
| | ATGTTCACTTGTTGGTTGGG |

| | |
|---|---|
| *ATXN3*_forward | AGATGCTATGTCATATGGTC |
| | GGACTAAATAATGATCTTCC |
| | ACTACAGTTTGTTTCACTGT |
| | TACCCACTTTCTCTAACTAG |
| | CACTCATAGCATCACCTGTT |
| *ATXN3*_reverse | CACCCTTCCCAAATTGACCC |
| | AGGTAGTTCGTTGTCTCTCT |
| | TGGTGGAGGATTGATTGAGG |
| | TCTGAGGATGTCAGATTATT |
| | TGCAAATACTGGTGAGAATT |
| *CACNA1A*_forward | ATCCCCCCGTCTCCTTACCA |
| | GGGTTCCAGTCCTGGAAGCT |
| | CATTGGTATCTGTAAGAGAC |
| | TTGCCCTGAGTTCCTGAGGC |
| | TAAGAAGGTGACATAAGGCT |
| *CACNA1A*_reverse | CAGTCTCTGAAGCAGTCTGC |
| | ACTCTGCACTGGGTAGGGCC |
| | GAGCACTACCTCCCCATGGA |
| | TCCGTGGAGATGCGAGAGAT |
| | ACACCAGGGATCTGGAGGTC |
| *ATXN7*_forward | CCAATTGAAGACGGTTGTGT |
| | CAAATGACTTCAGTCCTTTG |
| | GTGAGAGAGGATTTGAACAA |
| | GCTCTCTCTGGAGAAGAACG |
| | CACTAGAGGGAGCATAGTTA |
| *ATXN7*_reverse | GAATTTCACAAATCACCTTG |
| | GGGAACTTACTCTCTTAACA |
| | CGAAAAGTGCTGAAACCTTC |
| | ATCAAGTGTGGACATCACAC |
| | CCCCAGGCTCAATCACGTTC |
| *ATXN8*_forward | ATCCTATTTCATTGACCACA |
| | ACACGTCAGATCTTTAGTTC |
| | TCTTGGGATGGAAGTGTTTC |
| | GGAAAAGATATCCCCACACG |
| | TCTGAAGTACCAAAATCTGA |
| *ATXN8*_reverse | AGTTCTCTCTAATTAAGCAC |
| | TCCATACTCCCAAACTTAAA |
| | TTATGCGAAATGTAAACGGA |
| | GACAATTAGAGCCACATTAC |
| | GGACTCTATGCCAGGTTATT |

| | |
|---|---|
| *TBP*_forward | CCCCTTTATAGTCACTCTGC |
| | CCTCAGGTAATATAGCAGGA |
| | AGTTCCAGCGCAAGGGTTTC |
| | GAAGATAACCCAAGGAATTG |
| | TAGCCATAGAACTGGCTTAT |
| *TBP*_reverse | GGGGAAAATATTTCAACTGT |
| | AAAAATCACAACTGGTAGGA |
| | ACCGCAGCAAACCGCTACAA |
| | AAAATCAGTGCCGTGGTTCG |
| | AATTTCTTACGGCTACCTCT |
| *HTT*_forward | CACAAGTGCTCATCTGGAAC |
| | *GGACGTTATCAGGCTCTACA* |
| | *ACTGCTATGACTTGGTGACT* |
| | *CTTCTCGGCAGGACAGGCAC* |
| | *ATCCCACCTGGTCTCGGGTC* |
| *HTT*_reverse | CTAGACTCTTAACTCGCTTG |
| | TTGGACCTGTTCCCCCCATC |
| | CAAGTAGTATTGGTCTGATA |
| | CAGAGTTCAACAAGTGCAGT |
| | CCAGTTTAAGCTATTGCAGC |
| *FMR1*_forward | AGGTAGACCAGATGAATAGG |
| | GTTGTTCCTCAGTATCATGG |
| | AGGCAATGCCCTCAAATATT |
| | TTCTGCTGACACTGTAATGG |
| | TAATAACAGGAGGAACTGTA |
| *FMR1*_reverse | ACAAAGACAAATTTCCTAGC |
| | ACATGTCCAGGACTTCATAT |
| | AACCTGCAGGAGACAGGATA |
| | CAACAGGAACCACTGCTCAC |
| | CAGAAAGTCCATCCTTATTT |
| *C9orf72*_forward | ACAGTGGTGTACAGTGTTCT |
| | AGCTAGAGACTGACACTTGT |
| | TTCTAGTATGACTGGAGATT |
| | GTCCTCTTAAGTCAAAGATG |
| | GTCACATTATCCAAATGCTC |
| *C9orf72*_reverse | ATCTACATCCTAGAGATGTC |
| | GGTACCAGGTTGTTGTCCTC |
| | ATGCTAGTATTAACCACCAT |
| | TGGAGTTGTATTCCCCTCCC |
| | TACCTGTTGGATTAGGTGGG |

| | |
|---|---|
| *RFC1*_forward | TACCACAGCCTAATCTCCCA |
| | GCCCTCACGTTGCCATCTCC |
| | GGGCCAGTCAAGGTACCAGC |
| | ACTCAAGAACATAACACACA |
| | GTAATTGGTCCTTAATCTAC |
| *RFC1*_reverse | GTATTAGTCCCACTTTACTG |
| | GATAGTTCAGAAATGAGGTG |
| | GGTTCTTAAATCCAGTTCTT |
| | TGATTACAACCATCAAGGAT |
| | CTTCAGAGCAGGTGGATTAT |
| *ATXN10*_forward | TGTGATATCTGAGGGAAAGA |
| | GTCTCTTGCCGGAATGTGTG |
| | CTTAATGCCTGTATAGGGTC |
| | CTTGGACCGCTGGTGAGGAG |
| | GTAGGAGTGTGTCCTTCCTT |
| *ATXN10*_reverse | *TCTTCTCATCGAGTTGTAAG* |
| | *GTCCGTTCTATAACACAAAT* |
| | *AATGCCTGGGCAGCAACATA* |
| | CATCATGAGACCACAGGAAA |
| | GTGTACAGTGAGGACTGTAT |
| *PPP2R2B*_forward | AGACTGCTATAATGGAGATA |
| | CAGCTATAATGAATTAAGGG |
| | TTAAAAAGGCTTCTGAGGGA |
| | GACAGAGACCACGTCTGTAT |
| | CTTTCCACCTTATGACCCCA |
| *PPP2R2B*_reverse | GTAAAGCACAAGCCTGAAAT |
| | CTGAACTGGGTAACATAAGA |
| | AGGAAGCATTAGCCTAAGGA |
| | AAGAATCTCCAGACTGGCTT |
| | GAGATCTCATGAAGTACCGC |
| *BEAN1*_forward | GAGCAAGGGGGATTCAGCCA |
| | GGTGCATGCACTAGAGTTCC |
| | GAAGTGGCCCTTGCTCTGGG |
| | GCAAGCAGCCCTCACCCACA |
| | ATGAGCAGGTCCCAATCCCC |
| *BEAN1*_reverse | GTGCCCTGGGGGATATGCCT |
| | TTGGCCTAAGTTGCTCTCCC |
| | ATTTACCCAGATGGGGAATG |
| | AGAGAGATATATGTTGGTCA |
| | AGGAGCTAGGAATTGGATGA |

| | |
|---|---|
| *NOP56*_forward | AAGTCTACCTCAGGAACACT |
| | CTCATCCTGACTTTCATTCA |
| | CTGCAGGACATTTGCTCAGT |
| | TGTGGAACCTGGAGCACATC |
| | GACACAGTTAGCCATAGGCT |
| *NOP56*_reverse | TTCTCCCCGAATACACTCGT |
| | GTGCTCTGAGGACTCCTTTC |
| | CTACTGACCTCACACTCCTC |
| | CTCCTTCCTATAGAATCAAG |
| | TGAGCTACAACCAGCTCTCT |
| *DAB1*_forward | CTGAATCCACACTAGTGAGA |
| | TCTCCATGCCTCGGTTGTCC |
| | GGATGTGGGATACTCGACGG |
| | GTATTGGCTGAACGGCTACC |
| | AGCCCCAGTAAATATTGAGT |
| *DAB1*_reverse | CTATCTTTAAACTACACAGC |
| | TGATTAACCACTGATATGGT |
| | ACCTTAGACCTTTGAGGAAG |
| | AGAATCACATAGCCCAGATG |
| | AATATTCTGTGCTCCATTGT |
| *GLS*_forward | TATGATGTCACAATGAACAG |
| | GGAGGTAAAGTCTGCCCAGT |
| | AGTGAGTTTATGCTCGGCAG |
| | GGAAAGAAGCGTCCCCTCTC |
| | GGTAAAGAAGCATCCCCTCA |
| *GLS*_reverse | TGAAATTCAATGTTCAGACC |
| | GGATTAGCTGACTTTGAGGT |
| | TTGCACCTGAGTAGATTTCA |
| | TGCTTCTACTACGTTGTAGG |
| | ACACTCCCAAATGGAAGTCA |
| *FGF14*_forward | TTTGAACTCCCAATCTAGCA |
| | AATAACCTCTTACTACACCT |
| | CTGGCTTATATCAGCAAATC |
| | ACAGCTTGGGTGGAGCTGCT |
| | CTTACCTTTGTGCAGTGATG |
| *FGF14*_reverse | CTGCACTTGTCTGTTAATGA |
| | GCACGCATCTGCTAGACGAC |
| | AAGCGCTTAAACACTATGCT |
| | TTTCCCTGTAAGGGACAGCA |
| | GCTAGGTTCTTGTGAATCTC |

| | |
|---|---|
| *CSTB*_forward | TGCTGCTGTTGATGCTCCTA |
| | GGTTCGGTTAAACCAGCGGG |
| | GAGACGTGCCAGCAAAAGCG |
| | GGACCCTCCCTTAGGGCACT |
| | TCAAGCCTCAGGTCTGAACT |
| *CSTB*_reverse | CATTAACTGCAGGACTAACT |
| | TTGCTGCTGACCGAGTGGGT |
| | CATTGCAGGCCTGTGGGTTA |
| | ATGGGAGGTAAAAGTCAGCA |
| | ATTCTCTAAGTGGGGAGACA |
| *ZFHX3*_forward | ACTCAAGAAGACAGATTCAG |
| | GAAGGTCATTAGTCTGTAAC |
| | GCCCTTCCTTAATGTAACTT |
| | GCTGCAGAAACTAAGAAAAA |
| | GATACTGCTGTTGTGCAGAC |
| *ZFHX3*_reverse | TCTTATAGTCTCACTTAATA |
| | TGTGGTAGATCCTTAATGTT |
| | TTACTCCATCCAACACAGGT |
| | GTTGATGGCTCAACAAGAGT |
| | CAAAGAGAGTCGTTCAGGTC |
| *NOTCH2NLC*_forward | GGCTGTCTTTGAAATAAGAT |
| | CCTTCAGTTGCAGAAACTAA |
| | TTCTAGAACTTGTCCTGTGT |
| | GCAAGAGAGGGGAGGGTGTC |
| | CTTATGTTTCCTAGTTACAG |
| *NOTCH2NLC*_reverse | TACATGAGATGGAAGCAATG |
| | CTCAGTTGACTCTGAATTTC |
| | ATCCGAGTCACCCTGACTGC |
| | CCTAATAAAGGCCAAACTAT |
| | ACTGCCTAGCCAGAGGACCC |
| *PRNP*_forward | TAAGCTGACACATACTGGCT |
| | GGGTCACCTATATATAGCTG |
| | CCACCTGCCACTGTCTCATG |
| | GTCCTACTAAGACCCTTTAA |
| | ACTCAACCAGCAGGTCTTAT |
| *PRNP*_reverse | GATCCTCTGATCACCGTAGG |
| | CAGGTGGATATTATTCACTC |
| | ATTAAGGCTCCCCTTAGAGT |
| | TTAGATCAGTCGGCAAGGTC |
| | TCAGGATGAGCTGAGTCCTA |

| | |
|---|---|
| *ARX*_forward | CGGAAATGCACTCTGGTATG |
| | TTCACTCAGCTCCTAACTCG |
| | GTCACATCTCTGTTTGACAA |
| | CATCTTCTCCAGTGAATAGG |
| | ATACCTCTGAGGCCCCCCAA |
| *ARX*_reverse | GTGTAAAAACACCGAACTCT |
| | TGGAATGGGCTTTCGGGGCC |
| | CTCCCAACCACCAAGCCGCA |
| | TCTTGTGGATTTCCAACGCG |
| | CAGAGAGCTGAAACTCCGTT |
| *SAMD12*_forward | GTGATAGAGCACTTTGGTGG |
| | CCTACTAGAAAATTCAAGCC |
| | AGTCTGTACATTCTCCTTTG |
| | ATATGACCTAAATGTTCTGG |
| | GAACTAGTGTGCTGGCCTTC |
| *SAMD12*_reverse | ACTCACGCTGTCACCTGTCA |
| | GGGTAGGTCCTCTTAGTTGT |
| | AATGGCCTCTCTCAGGCTGT |
| | GGAGCCATCAGTATCATTAG |
| | TGCCATAAGCACTCACACAT |
| *STARD7*_forward | AGGAACTCGGCAACAGCCAC |
| | TAGCTGGATGGAAGGGGCAC |
| | TTTACGGCTCACTAGGCTGT |
| | GGAGAGCTAAGTTAAGGGGA |
| | GCTCCACAGCACTGGTAAGG |
| *STARD7*_reverse | TACCAGGACATGCTAGGAGC |
| | GGGGCTGCATTCACCTATTT |
| | TGGCATTCTTTCCCCCAGAT |
| | TATTGACAAGGCCAGGGTAC |
| | TGTAACAAAGAAACTGGTCT |
| *MARCHF6*_forward | GAGTTGGATCAGAAGGAGAG |
| | GCCAATACGTGCTCTGCTAC |
| | GAGGGTCACATACCTGCCCT |
| | GGAAGGCACCTAAATACAAA |
| | GTCTGAATGCGAAAAGGTTA |
| *MARCHF6*_reverse | GTCCCCTGAACAAGAGTTAG |
| | ACCCAGCTACCTGGATCAAA |
| | AGTGATCTTCATAGTCTTAC |
| | GTAGACATGTACCCAATATT |
| | TGCAGAAAACATTGTTGCAC |

| | |
|---|---|
| *TNRC6A*_forward | GCAAAGGAGGTTACTACAAT |
| | TACGAGACCTGCAGCATGTT |
| | GCACCTCAAGAACAGAGATG |
| | TTGGGCAGAATAGGGGAGTG |
| | CAGTGCTTAGTATCAATAGT |
| *TNRC6A*_reverse | CTTACAAGGCACATGACTCT |
| | CAGGCGTAGCTGAAGAACAG |
| | ACATCTCCAGAACTGTGCAG |
| | TAATTACTCCTAACAGAAGG |
| | CTGAGAGCCTACCAGGAGTA |
| *RAPGEF2*_forward | ATGACAGCATAGTAGGATTA |
| | GTGTAACTTTGTTTAATCCC |
| | CGTGTCATGTATTACATAGC |
| | CTTAAAGGTTGAGTGCGCGG |
| | GAAAGGCAGTTCATCATGTA |
| *RAPGEF2*_reverse | GGAGAGGTCTCGGATTTGAC |
| | TGAAGGATAAAGGGACACGG |
| | GTAATCACTTACCAAAAGGT |
| | TTGCAAACCATGACAGTGAA |
| | GTTGTACCACTTCCACAAAG |
| *YEATS2*_forward | TAGTAATATTGAGCACTTCC |
| | ACATCTCTTAGGTCATGTTA |
| | TACACTCTTATTAAGATTAG |
| | TGTCTCTTCTAGAACTTAAG |
| | GAAATCATTATCAGCATCTT |
| *YEATS2*_reverse | GCTATAAAATACAACACTGG |
| | AAGACATTCTATAAAATTAT |
| | AAAACTGAAACCACTAATAT |
| | TATAGTCTATGTATATTCCC |
| | GCATAATGCTAACAATTAGG |
| *JPH3*_forward | CTGGAAGGCCAGGCAGACTT |
| | CAAGGCACTTTACGTTGGTG |
| | CTAGTGCCTGGCTCACATGT |
| | CTGGGTGGTACTACGTTTAT |
| | GCCAGCCCTACTTTATCGGG |
| *JPH3*_reverse | GAAGCTACCAGCGGGGGAAG |
| | AGAGATGGTTTGGCTTGAAG |
| | CTGGTGGTTGAGTGTGCTGG |
| | TGCACCGACTGAGTAGGACC |
| | TGCCAAGTGGCAGCCCTTCC |

| | |
|---|---|
| *AFF2*_forward | TACTCAGATTCTGTGAGCCT |
| | ATGATTGACATAAGGTCAGA |
| | ACTTTAGAAGAGCTCTGAAA |
| | ATTGCTACCTATACAGAAGC |
| | GGGAGAACTGAGATTGAAAT |
| *AFF2*_reverse | GCCGGGTTGCTGCAGTGCAT |
| | GATTGTAGCTAACGAGGCCT |
| | TAGTATCTCCAAGTTGGCAA |
| | GAGGGTTTGGGGAGTTGCGG |
| | CTACAGGATGCTTAAATGTA |
| *SOX3*_forward | GAATCAAGCTTTGGTCATTC |
| | ATTGTGGGAAGGTTAGGATT |
| | AACCTTCATTTGACCACATC |
| | CATAGCCCTAACTGTCAAAT |
| | GTCTCCACAAAATTGTTAGT |
| *SOX3*_reverse | CGATATGAAGGGGTAGAAGG |
| | TTGAGCAGAATAATATCCCG |
| | GACCCTTGAAAAGTCCAGTA |
| | GTGACATCTCTACTTAGAAA |
| | TGTGTATCCTGTCTAAATTC |
| *DIP2B*_forward | GACCCTTCGACTTGGATGCG |
| | AATGGCAATTACTATAAGAC |
| | GGGTGTTCCACCTTCCGGCA |
| *DIP2B*_reverse | GGATAGGTTTACAAAAAAGG |
| | GTGGCCCCGTTTCCCATGAT |
| | TGCTAAGTCTTAAACAGGCT |
| *ZNF713*_forward | TTCCTGCCTAAAAGGCTGTT |
| | ACACTCTCCAAAAGGGAAAG |
| | GGCTCTTGGATGTGGTGCTC |
| *ZNF713*_reverse | TGAGCTTCCCTGGTTCTGTA |
| | GAGTGGTCTTACCCCACACT |
| | CAAGCCCACATGACTGGTGT |
| *CBL*_forward | TCCTCATCTAGAAGCCTGTT |
| | CGCTTTGGTCTCCAAGTGTC |
| | AGGAGCTACCTTGCTTGGTC |
| *CBL*_reverse | GTACACGGTTCAGTACTATT |
| | GAGTCGTAACAACAACAACC |
| | AAGGCAAGGTGGATAGGGTA |

| | |
|---|---|
| *AR*_forward | GAACCAACATTGCAGGTAGA |
| | GACAGTGACAGGACTTAAAC |
| | ACTGAAAGCTATACAACTTC |
| | TGCTCTGCATCTTTATTCAT |
| | GTTCCATTGACATAAAACTC |
| *AR*_reverse | CAGCTCCATAAAATATCATC |
| | TAAACCCCAAACTCTAATTC |
| | CTTGATAACTGAGAGGAGAG |
| | TGGGCAGAAATAATCTGTCT |
| | GGTCCTCAGGTGTTGATTGT |
| *PRDM12*_forward | ACTCTAGGCTGAGTCTCAGG |
| | CAATGGTCTGAGGGTCTTCC |
| | AACGCCAGAGTTCTGAGTGT |
| | TGGTTTCAGACACCTCTAGA |
| | GGTGCCAAGGCCAGCACTTC |
| *PRDM12*_reverse | ATCAGAAAGGGCTGAAGGAA |
| | GATTCAAGAATAGTGGGCAT |
| | TGCACACAGATTTCAGCCTA |
| | TTCAATCCTCATCTACCTAC |
| | CAGAAAAGCTTGCCTGACAG |
| *WDR7*_forward | GATCTGATCAATTAGAAAGA |
| | CAACCTGGTATGCTACGTGA |
| | CTGAAATGTACACTACCTGT |
| *WDR7*_reverse | GTCTCTACATATACAAGGAC |
| | ATCTCAGCCTAGCTCCCGAG |
| | GGGTGGAAGTATTTCCCTCC |
| *NIPA1*_forward | GTGTACCACGTATCTCAGTT |
| | GCTAAGGGAGACGTTAAGTT |
| | TACTGGCCCCTGAATCTTTC |
| | ATACACTGCAAAGAGATCTT |
| | GCCTGTTAGGGGCCACAGCA |
| *NIPA1*_reverse | CTGCTGCTTGCAACAGATGT |
| | CTGGGTTAACCTGAAGGGAA |
| | AATCTAGAAATGCTACCACC |
| | GACGACGCTCCCTGGACTAC |
| | CTGCTATTCTAATGTTGCCA |

| | |
|---|---|
| *VWA1*_forward | CGGCGGAAGAGAAGCAAGGA |
| | TCTTCCCCCGGGTTCTTGCC |
| | CCCCTCCACTTCAGGACCCT |
| | GCGTTCTTTCCCGGTGGCTG |
| | CCCACCCCACTGGATGCAGT |
| *VWA1*_reverse | GCTCCTCCCTGAACAGTCCT |
| | AAGCCTTTGCGGACCTGACC |
| | GATTCCCAGTGGCTGCTAGC |
| | TTCCTGTCAGTGCGGTCCCG |
| | CAAGAAGATGGTGTGGTTGC |
| *DMPK*_forward | CCCTGCTGACCAGACAGGCA |
| | TGCCTAGGGCGTGAGCCTCT |
| | CCGTCTCTGGCTTCAGTGGC |
| | TGGGAGACGAGCATGCTGGT |
| | ACGATGGGGCTGCCAGACAC |
| *DMPK*_reverse | AAAACCTCAGAGTCACAGAC |
| | GACTCCTAAGAGGCCAGAGT |
| | TCCTGCTCCTTGGCAGCCAA |
| | TTCGGGAAGGTGCGCCGCTA |
| | CACATCTGTGTGTCTTGCGC |
| *NUTM2B-AS1*_forward | ACGAGGTCAAATGTCAAAGA |
| | AGGAAGCTGCTCTGACATTG |
| | AGGACTGCTGGATCATATAT |
| | GGCTTATCATGATATCCTCC |
| | CTCCATACTGTAATCCACAG |
| *NUTM2B-AS1*_reverse | CTCTAGTGTTATTCATTGTG |
| | AAATTCTTATGGCCATACCA |
| | CCCCCATATACTTTATTTCT |
| | AAGATGCAGGCTTGTTACAT |
| | TGTATACCTGTTAAGTGGTC |
| *PABPN1*_forward | AGTAAAGGAGAAAGACTCCA |
| | ACACTTAGCACATCTCAAGT |
| | CTCCATGGTCCTTTGTTTAT |
| | ATTCTCTATGCATCAACAAC |
| | GAGCACAGAGTGATAGGCAA |
| *PABPN1*_reverse | TCTGCGATCTAAGAGAGAAT |
| | GAGCACACAAAAGCGCCATT |
| | ATCCAAAGCTCTAGGGCTGC |
| | TTGAGGTAGAAAAGGTAGCT |
| | TGAAGTCTGAAGGTTCAACC |

| | |
|---|---|
| *CNBP*_forward | ACATGAGGCCCAGAATTTGG |
| | ACTGACAGAACCCTGCATGA |
| | ATCACACAGCAGCATGGAGG |
| | ATTTCGAAGTGGGGGGAAAC |
| | TATGAAACCCAGGAAAAGGT |
| *CNBP*_reverse | TGTGGATCCTAAATTCATCT |
| | GTAGAAGATACTTCTGTTAG |
| | GTAAATTTCTCAGCCACACG |
| | AGAGGTTCTGGCCTGGTTGC |
| | TAAGTGGTGCCTTATATTAG |
| *LRP12*_forward | AACACAGTTAGCTATGCTTT |
| | AGAAAGAGTACATATCTGAC |
| | ACTTAACTCTCTCAAAAAGC |
| | TAAAGACTTGCAGTAAGAAC |
| | TGTGGTATCACTGGTAAAGT |
| *LRP12*_reverse | TTGACTACTTGGTACAATGG |
| | TTATTGCATGCTCTATGGTC |
| | CCACTGGACTTAAGTATACA |
| | AAGGCTGCTGTTATATAGGT |
| | TAGGGTTCTCTGATCTTCAA |
| *GIPC1*_forward | TGACGCCAGGCACAAGATGC |
| | CAAAGTGGTGGTGCGGTGTT |
| | CTTATGTAAAAGGGACTTCC |
| | GGGCCATGCTGGATTCAGCT |
| | CTAGTGTGGTTGCCTCCTGT |
| *GIPC1*_reverse | TGTGGAAGAGTCAGGTGCGG |
| | GTAACACCATAGAATGGGCC |
| | AACCTCAGGTGGACTACGGA |
| | CTGCTGAACAGCACAGGTTA |
| | GACCGAGGCACAGAAGAGGA |
| *PLIN4*_forward | CGGCATCCCGCTCCTGTGGG |
| | CTCCTCACCATGGCCTTGGT |
| | CATATGCAGACGCGTCCCCT |
| *PLIN4*_reverse | AGGTAGCCATGTGACCGTCC |
| | CAGGACCGTGCCAGGTGCGC |
| | GGGCTCAGCACGAAGGTCCT |

| | |
|---|---|
| *ZIC2*_forward | CGCAGAAAGACTGGAGCGCT |
| | AAAGGGGCGGTTTGACCGGG |
| | ATCGAAATCAACGGAGGCGG |
| | CACTGCAAGTTTACTACGCG |
| | AACAATAACCGCCGCGCGCG |
| *ZIC2*_reverse | AGCTCTGGGACAGCCAGGTA |
| | TGTGATCTGCCCTTGATCTC |
| | CCTTCCTTCCTAGCCAACCC |
| | TGAGCTCACCAGGGACCTAC |
| | TCGGTGGAGTAGTGTGGGGG |
| *XYLT1*_forward | ATCATAAGTGGCAGCTCTTA |
| | TTATGGATCATAAACAGCTC |
| | TTATGTCAGAGATATGCCTA |
| | ATAAGAACTTACGCCATGGC |
| | TAAAGTATGAAAAGACCGGA |
| *XYLT1*_reverse | CTGGGGCTTCTTCAGAGCTC |
| | CGATGAAACAAGATTTGGGG |
| | AGAGAAGAACCCTTTCCCCC |
| | TTTGCACACTGACTCCGCTG |
| | TAGCAGAGCTGAGTAAACCT |
| *FOXL2*_forward | TGTCTGGCTGCATGAAGCCT |
| | GCTCATCTAAGCCTCAGATC |
| | TACCCTCTGGAGTAGGGTAC |
| | CCTTGGTGCATGTACTTCAA |
| | CCAACCCACTTACTCTGACT |
| *FOXL2*_reverse | ACTGGAACCGTATCCGCAGG |
| | AGATTGCTCGCGCCGGCTGT |
| | TTGCAGGATTCCCGGCCCTG |
| | GACGCTTCCCTAGTAGAGCT |
| | AACCTGGTTCAGGCAGCCGG |
| *HOXA13*_forward | CAACATTATCCATGGGTCAG |
| | CATGCCCACTTTAGGTTATC |
| | TTGGGAGAAAGTTGTTACTT |
| | GTACTGTTTGGAACAAGAGG |
| | TCGTTCTGTATCCAATCCTG |
| *HOXA13*_reverse | GATGCAAAGAAATAGTGCCA |
| | AGTGTCTTTAGAATCAACCA |
| | GTTAAGATGTTCTGCAAAAG |
| | CCAAATGATCTGCTCAAGTT |
| | GGGTACAGCTTCTACCTGAT |

| | |
|---|---|
| *HOXD13*_forward | GGACTCTGCGGAATGGAGAT |
| | CCGACAGAGTGAGAATTTAC |
| | AACAAGTTACTCTGGGCCTC |
| | AGATTAGTGCTTCTCTTCAA |
| | TTTCTGATTTGGAGGGGGTT |
| *HOXD13*_reverse | CAATCTCCCCTACCCACGAT |
| | TCCAAGCCCCCTTAGCCTTC |
| | CGCGGCGCTAAAAACCTGCC |
| | CTCTTGAATCACGCGACACG |
| | AGCCGGAGATCTTGGCGGCA |
| *PHOX2B*_forward | GAGCTTGTTCACACTGTGCA |
| | CTAAGGCTCCATCATATTAC |
| | GATGACCCAAATTAAAACGA |
| | AATGTAGTCCTCTGGGAAGC |
| | ACTCAAAGTTGCTCAAGACC |
| *PHOX2B*_reverse | CAGGGTTTGTTTACACCACA |
| | ACTTGTTACAGAGCACATAA |
| | GCAATAAAACACGGTGGCAG |
| | TCACCCTGCAAAGAAGCACA |
| | ACCAGGATCCTCGGTAGGTC |
| *TCF4*_forward | CATTGATATCTGTCATAAAT |
| | GTTTCCTCAGCTATGAATAC |
| | ACCAAGGTTATTTCCAATAA |
| *TCF4*_reverse | ATCATCTAAGCAGAACCTGT |
| | CCCAAACTTCATCCTATCAT |
| | GCTCCTGGGGTCCAATCTGC |

**Table S4. Selected crRNAs for nCATS determined through in vitro cleavage.**

| Gene | crRNA sequence (forward) | crRNA sequence (reverse) | Size (bp) |
|---|---|---|---|
| *ATN1* | AATGAATACAGCTGGCTATA | CTAAAACTCGTGCTTCTCTT | 8641 |
| *FXN* | CTTGACTACCTCCAAGGAAG | AGGTGTCCCCTTGTCACCCC | 8498 |
| *ATXN1* | CTACCTGCTCGTCTGCACCC | TTGTGTCCTTGATTTCTCAT | 8991 |
| *ATXN2* | TGAGCCCCACCTGTTAGATC | CACCCAACAATTTCGTTTGT | 12601 |
| *ATXN3* | CACTCATAGCATCACCTGTT | AGGTAGTTCGTTGTCTCTCT | 11304 |
| *CACNA1A* | CATTGGTATCTGTAAGAGAC | GAGCACTACCTCCCCATGGA | 10504 |
| *ATXN7* | GTGAGAGAGGATTTGAACAA | ATCAAGTGTGGACATCACAC | 9654 |
| *ATXN8* | GGAAAAGATATCCCCACACG | TTATGCGAAATGTAAACGGA | 9322 |
| | ACACGTCAGATCTTTAGTTC | GGACTCTATGCCAGGTTATT | 9198 |
| *TBP* | CCCCTTTATAGTCACTCTGC | GGGGAAAATATTTCAACTGT | 8371 |
| | GAAGATAACCCAAGGAATTG | ACCGCAGCAAACCGCTACAA | 9995 |
| *HTT* | ACTGCTATGACTTGGTGACT | CAAGTAGTATTGGTCTGATA | 9359 |
| *FMR1* | GTTGTTCCTCAGTATCATGG | CAGAAAGTCCATCCTTATTT | 11314 |
| *C9orf72* | GTCACATTATCCAAATGCTC | TACCTGTTGGATTAGGTGGG | 12688 |
| *RFC1* | TACCACAGCCTAATCTCCCA | CTTCAGAGCAGGTGGATTAT | 9801 |
| *ATXN10* | GTCTCTTGCCGGAATGTGTG | TCTTCTCATCGAGTTGTAAG | 9994 |
| *PPP2R2B* | CTTTCCACCTTATGACCCCA | CTGAACTGGGTAACATAAGA | 8770 |
| *BEAN1* | ATGAGCAGGTCCCAATCCCC | TTGGCCTAAGTTGCTCTCCC | 11374 |
| *NOP56* | CTGCAGGACATTTGCTCAGT | CTACTGACCTCACACTCCTC | 9473 |
| *DAB1* | GTATTGGCTGAACGGCTACC | CTATCTTTAAACTACACAGC | 8696 |
| | CTGAATCCACACTAGTGAGA | ACCTTAGACCTTTGAGGAAG | 9117 |
| *GLS* | TATGATGTCACAATGAACAG | TGCTTCTACTACGTTGTAGG | 10676 |
| *FGF14* | CTGGCTTATATCAGCAAATC | GCTAGGTTCTTGTGAATCTC | 13422 |
| *CSTB* | TGCTGCTGTTGATGCTCCTA | ATTCTCTAAGTGGGGAGACA | 9485 |
| *ZFHX3* | GAAGGTCATTAGTCTGTAAC | TGTGGTAGATCCTTAATGTT | 11202 |
| *NOTCH2NLC* | CTTATGTTTCCTAGTTACAG | ATCCGAGTCACCCTGACTGC | 11331 |
| *PRNP* | CCACCTGCCACTGTCTCATG | CAGGTGGATATTATTCACTC | 10286 |
| *ARX* | TTCACTCAGCTCCTAACTCG | GTGTAAAAACACCGAACTCT | 8150 |
| *SAMD12* | ATATGACCTAAATGTTCTGG | GGAGCCATCAGTATCATTAG | 11347 |
| *STARD7* | TAGCTGGATGGAAGGGGCAC | TATTGACAAGGCCAGGGTAC | 9389 |
| *MARCHF6* | GTCTGAATGCGAAAAGGTTA | ACCCAGCTACCTGGATCAAA | 8806 |
| *TNRC6A* | TACGAGACCTGCAGCATGTT | ACATCTCCAGAACTGTGCAG | 8723 |
| *RAPGEF2* | GTGTAACTTTGTTTAATCCC | GTTGTACCACTTCCACAAAG | 11512 |
| *YEATS2* | TACACTCTTATTAAGATTAG | GCATAATGCTAACAATTAGG | 12905 |
| *JPH3* | GCCAGCCCTACTTTATCGGG | TGCCAAGTGGCAGCCCTTCC | 8888 |
| *AFF2* | GGGAGAACTGAGATTGAAAT | CTACAGGATGCTTAAATGTA | 8957 |
| *SOX3* | GTCTCCACAAAATTGTTAGT | TGTGTATCCTGTCTAAATTC | 9292 |
| *DIP2B* | GACCCTTCGACTTGGATGCG | GGATAGGTTTACAAAAAAGG | 12954 |
| | AATGGCAATTACTATAAGAC | GTGGCCCCGTTTCCCATGAT | 9713 |
| *ZNF713* | TTCCTGCCTAAAAGGCTGTT | TGAGCTTCCCTGGTTCTGTA | 11467 |
| *CBL* | CGCTTTGGTCTCCAAGTGTC | GTACACGGTTCAGTACTATT | 9670 |

| | | | |
|---|---|---|---|
| *AR* | ACTGAAAGCTATACAACTTC | CAGCTCCATAAAATATCATC | 9239 |
| *PRDM12* | AACGCCAGAGTTCTGAGTGT | CAGAAAAGCTTGCCTGACAG | 10313 |
| *WDR7* | CTGAAATGTACACTACCTGT | GTCTCTACATATACAAGGAC | 9856 |
| *NIPA1* | TACTGGCCCCTGAATCTTTC | CTGCTATTCTAATGTTGCCA | 13014 |
| *VWA1* | CCCACCCCACTGGATGCAGT | GCTCCTCCCTGAACAGTCCT | 8290 |
| *DMPK* | CCCTGCTGACCAGACAGGCA | CACATCTGTGTGTCTTGCGC | 9306 |
| *NUTM2B-AS1* | CTCCATACTGTAATCCACAG | TGTATACCTGTTAAGTGGTC | 16227 |
| | AGGACTGCTGGATCATATAT | AAGATGCAGGCTTGTTACAT | 15116 |
| *PABPN1* | GAGCACAGAGTGATAGGCAA | TGAAGTCTGAAGGTTCAACC | 10948 |
| *CNBP* | ACATGAGGCCCAGAATTTGG | GTAAATTTCTCAGCCACACG | 9301 |
| *LRP12* | AACACAGTTAGCTATGCTTT | TTGACTACTTGGTACAATGG | 9613 |
| *GIPC1* | CTTATGTAAAAGGGACTTCC | TGTGGAAGAGTCAGGTGCGG | 9241 |
| *PLIN4* | CGGCATCCCGCTCCTGTGGG | CAGGACCGTGCCAGGTGCGC | 14136 |
| *ZIC2* | CGCAGAAAGACTGGAGCGCT | CCTTCCTTCCTAGCCAACCC | 9340 |
| *XYLT1* | TAAAGTATGAAAAGACCGGA | AGAGAAGAACCCTTTCCCCC | 8547 |
| *FOXL2* | CCAACCCACTTACTCTGACT | ACTGGAACCGTATCCGCAGG | 8330 |
| *HOXA13* | TTGGGAGAAAGTTGTTACTT | GGGTACAGCTTCTACCTGAT | 9327 |
| *HOXD13* | AACAAGTTACTCTGGGCCTC | CAATCTCCCCTACCCACGAT | 8255 |
| *PHOX2B* | ACTCAAAGTTGCTCAAGACC | CAGGGTTTGTTTACACCACA | 8171 |
| *TCF4* | CATTGATATCTGTCATAAAT | ATCATCTAAGCAGAACCTGT | 10746 |

**Table S5. PCR primers targeting DNA fragments for in vitro cleavage assays.**

| Gene | | PCR primers (5'→3') | | PCR product size (bp) |
|---|---|---|---|---|
| *ATN1* | front | forward | GGTGGGTGTTGTTCTTCGGA | 2573 |
| | | reverse | TGTGGGCACCTGGAACTTTT | |
| | back | forward | GGTTTGGGTGTCATAGGCGA | 2418 |
| | | reverse | CCCCCATATCCCGCAAAGAA | |
| *FXN* | front | forward | CAGGCAAAGGGATGGAAGACT | 1757 |
| | | reverse | TGGAAGAAGTGGTGTTGGCA | |
| | back | forward | GCCTTGGGCAGCTTTTAGAC | 2257 |
| | | reverse | AAACCTCACAACCGTGAGCA | |
| *ATXN1* | front | forward | AGTCTTTGGAAGGGCCCAAG | 3573 |
| | | reverse | GCGACAGACGGCCCTAATAA | |
| | back | forward | TCTCCGAGGCTCCATCAACT | 2220 |
| | | reverse | CGCTTAAAGACTGGGACGGA | |
| *ATXN2* | front | forward | CTCTTGCTAGCGGTGCTCTA | 5791 |
| | | reverse | GCCAAGGAGGGCAAATTGTTT | |
| | back | forward | TGTCATCCCCAAATCCCGAG | 3935 |
| | | reverse | GCCTGAAGCTGACATAGCTGA | |
| *ATXN3* | front | forward | CCTTGGAACCAGAGGAGCAG | 5059 |
| | | reverse | ATAAGGCGCAGGAAGAAGGG | |
| | back | forward | AGGCAGGCAGATTACGAAGT | 4497 |
| | | reverse | CTGCCCCTCCACTCAAGAAG | |
| *CACNA1A* | front | forward | CCGAGTGCCAAGAGGTAGAC | 5677 |
| | | reverse | GACAGGAGCCCAGATGGAAC | |
| | back | forward | CTACACCCCCATTGCAGGAG | 5521 |
| | | reverse | CCTAGAGATCCCCTGAACCG | |
| *ATXN7* | front | forward | GCTGAGCCCCCAAAAATGGT | 2362 |
| | | reverse | AGCAAGCTGACACACTGCAT | |
| | back | forward | TACACGTCCCCACCCCTAAA | 2652 |
| | | reverse | AACTGGGGAAAAGTCCCTGC | |
| *ATXN8* | front | forward | TCCTTCCAGAAATTAACAAGCTG | 2429 |
| | | reverse | TGGATCTTGGAAGACTGTTCTTG | |
| | back | forward | ATGTGGCAACCCCTACAGATT | 5698 |
| | | reverse | CAGTCATTCCCATTTGCAAAAACAC | |
| *TBP* | front | forward | TCCAGGGGTGTATATGGGCT | 3661 |
| | | reverse | GGTGCAAGAGGCCTAGCTTT | |
| | back | forward | AGAATCCCTGGGAGTACAGCA | 2732 |
| | | reverse | CCACAGCCCCCTTCTATGTG | |
| *HTT* | front | forward | ATGGAAAATTCGGGGGCCAAT | 3457 |
| | | reverse | ACAGTGCTGGGGTGATTTGT | |
| | back | forward | CTAGCATGCTTGGGAGGGTC | 3397 |
| | | reverse | ACCTAGGAGTGGGAAAACTATAACC | |
| *FMR1* | front | forward | GTCTGCAAGAAGCCCAATTT | 4913 |
| | | reverse | GATCATGAGCCTGCTCTTCC | |

| | | | | |
|---|---|---|---|---|
| | back | forward | GAGTTGCTTGTCCAGTGGGA | 4527 |
| | | reverse | CGGGTCATGTGCGACTACTT | |
| *C9orf72* | front | forward | TTTTGCCAGGCTCTGTATGC | 4292 |
| | | reverse | GCAGGGGTAGGAAATTAACCCA | |
| | back | forward | AAGCCATCTTTCATGCTGCT | 1591 |
| | | reverse | TCCTCAGCCTGGACTTCACT | |
| *RFC1* | front | forward | TCCAGTTGAGGTTGTTGGAAG | 3299 |
| | | reverse | ACAGCCCTGATCCACAGTTG | |
| | back | forward | GAGTGTGGGGGAGTTTAGGG | 2097 |
| | | reverse | AGGCCAAAAACAGGCCAAAAT | |
| *ATXN10* | front | forward | CTCACCTGACCACCCCTCTA | 2224 |
| | | reverse | GATACTGTCTCCTGGCACTGA | |
| | back | forward | TGCTATACCAGTCAGAATGCCAG | 3640 |
| | | reverse | TGTCAACTGAGGGTATGCGT | |
| *PPP2R2B* | front | forward | TGACTGCAGGCAAGTTCATC | 2124 |
| | | reverse | CATCCTACACAGCAGCCAGA | |
| | back | forward | TTGCTCTAGGTGCCAAGGATA | 1925 |
| | | reverse | GAGGGGACTTTGCACCGATA | |
| *BEAN1* | front | forward | GCCCAGCAGAGTCTCATAGAA | 1906 |
| | | reverse | CCTCGGAGAGGCAGTAACAG | |
| | back | forward | ACTTGGAACCAAGCCAAATG | 1939 |
| | | reverse | ATTGCAGGAATGTGGTGACA | |
| *NOP56* | front | forward | TAGCCAGTGCATGCCTGTAG | 3446 |
| | | reverse | TGGGACAGGGTCTCCTGTAG | |
| | back | forward | TCAGCGACATTGGATGCCTT | 2226 |
| | | reverse | GCCCTGTCCATACCCATGTA | |
| *DAB1* | front | forward | CCTCAGGTCCCATGTGTCTT | 1383 |
| | | reverse | GCCATCGCCCTAATTTTGGT | |
| | back | forward | TATGTCACGGGTGTCCCTCA | 2293 |
| | | reverse | AATGGCATCCCTCCTAGCCT | |
| *GLS* | front | forward | CGGTCACAAACCAACCCTTG | 2478 |
| | | reverse | GTTTCCTGCTTTAACTGGCCAAA | |
| | back | forward | GCTGTTTCCATCACGCAACT | 5736 |
| | | reverse | TCGGGTGGGAAGACTAGGAT | |
| *FGF14* | front | forward | CGTGTCTGCATGTGAGATGG | 2366 |
| | | reverse | CCTGGGTCTTCACATGGTCT | |
| | back | forward | AGACGCAACCTGAGGTAACAA | 2356 |
| | | reverse | TCAGCAACCAATGAGTGACCTA | |
| *CSTB* | front | forward | TGGGGGAAGCTGGATTTAGG | 2303 |
| | | reverse | CTTCAGAGTGAGGGTCTGCG | |
| | back | forward | TGCTGAGGAAGAAGGCACAT | 3160 |
| | | reverse | TGGCTAGCCGTGAAAGTTGA | |
| *ZFHX3* | front | forward | ACTACTGGTGGGCATTGAAATACT | 3951 |
| | | reverse | GGTTTTCTCCCTATAGTGTGGGTT | |
| | back | forward | GACGAAAAGAACAAGACTGCTTCA | 5412 |

| | | | | |
|---|---|---|---|---|
| | | reverse | GGTCTTCAAGAACAAGGTTTACGG | |
| *NOTCH2NLC* | front | forward | CATCCAACTTTGAGGGGGCA | 5053 |
| | | reverse | CAGTGATCCAGTCCCATCCG | |
| | front | forward | AGGGCAAAACTATGTCAACAAGC | 1523 |
| | | reverse | AACTGGGGAGCAATGCAAGA | |
| | back | forward | TGGGTGCAGTATGTCTGCATT | 2234 |
| | | reverse | TGTGATTCTCCCACCAGTGC | |
| | back | forward | GTGGTGTATTGGGGGAGCAT | 1712 |
| | | reverse | TGTGTCTTGACAGCGTCTGA | |
| *PRNP* | front | forward | ACATCTCAGGTGAAGACGCC | 4006 |
| | | reverse | GCGAGGACATGGCATAGTGA | |
| | back | forward | TTCCAACCCGTGGTGTTCAA | 2225 |
| | | reverse | AGGATCCCCAGGCTCTCATT | |
| *ARX* | front | forward | CCCTAGCCCCAAAGCTCCTA | 1677 |
| | | reverse | GCGCATAGAAATCAGAGGGC | |
| | back | forward | CTTCTCCCCGAGGTGAATGG | 1675 |
| | | reverse | TCTTCCCTCTTTCCCCCACA | |
| *SAMD12* | front | forward | CTCAAGGGACCAATCGACTCC | 2078 |
| | | reverse | CAACACCACAGAAACGAGGAAGA | |
| | back | forward | GAACCCATGGACACAGGGAG | 2565 |
| | | reverse | ATTTGGGCACCAAGACCCAA | |
| *STARD7* | front | forward | AAGGCAGACAGCATGAGTCC | 2032 |
| | | reverse | TGGGATCGCCCAGAGTTCTA | |
| | back | forward | GACTCACCGGCTGCAAAAAG | 2577 |
| | | reverse | AAGCAGTGGGTCAGATGCTC | |
| *MARCHF6* | front | forward | CAGGCAACTAGCTGGACCAA | 1434 |
| | | reverse | AGAATTGCCTGAGTGGGAGC | |
| | back | forward | AGTGATGCTGGAAGTGCTCC | 5229 |
| | | reverse | ACACCTTCATACAGCCCTGC | |
| *TNRC6A* | front | forward | CAGGGTCAGTTTGGCAACTC | 2320 |
| | | reverse | TTTGAACCCACGTAGCCCTG | |
| | back | forward | GAACTCCTGGGCAATCCTCC | 1137 |
| | | reverse | ACTTGGGGCCTCAGTTGTTT | |
| *RAPGEF2* | front | forward | GGTAGGCACAATCGGGTGAT | 3291 |
| | | reverse | ACTCACCAGGAGTAGCACTGA | |
| | back | forward | TAGCCCAGACAACACCGAAC | 2638 |
| | | reverse | GCTGTAACTGTGCAGGGACT | |
| *YEATS2* | front | forward | TTCCAGGTGCATTAATGGAGAGAA | 7660 |
| | | reverse | GCATGTGGGCATATATTTGAGCTT | |
| | back | forward | AAGTCTGGTTCCTTTCCTTTCCTT | 7206 |
| | | reverse | CAGGAGATGATGACCTAGATGGTG | |
| *JPH3* | front | forward | GATGCAGGGGGAGCTTTCTT | 533 |
| | | reverse | TGCAGTCAGTTACTCCAGCG | |
| | back | forward | TCCACATTGGTCCGTGTCTC | 793 |
| | | reverse | CTCCCAGTTGACCAGCAGAG | |

| | | | | |
|---|---|---|---|---|
| *AFF2* | front | forward | CCATGTGCCCTTGGTGAAAT | 1597 |
| | | reverse | GTCATGCAGGTTCGGGTAGT | |
| | back | forward | CTGTCAAGGCTGAGAAGGGG | 1147 |
| | | reverse | GGCAGTTTTCCGCTTTGAGG | |
| *SOX3* | front | forward | CAGACGGGCAGATAAGCACT | 5991 |
| | | reverse | CCTGGCATTTTCCCCGTAGA | |
| | back | forward | CATGGGTTCGGTGGTCAAGT | 6248 |
| | | reverse | GTACACGCAGGCAGACTAGG | |
| *DIP2B* | front | forward | ATTGGACCCTTCACAATGCAAAAT | 5942 |
| | | reverse | GCACCTTTACCAGTGCTGTTTATT | |
| | back | forward | CTGCTTCCAGCTGATTGTTGTTTA | 6322 |
| | | reverse | GTATAGGGTATAGGTGCCAACAGG | |
| *ZNF713* | front | forward | CATGCTTGTGGATGACTCTGTAAC | 6498 |
| | | reverse | ACAAGAGAAAACTCCATCTTCAAAA | |
| | back | forward | GTTCTTGTAACACGTTGGTGTCAT | 6372 |
| | | reverse | CTCATTGTCTTGGTTTCACTGCTC | |
| *CBL* | front | forward | GGAGCAGATCTGTAGAAATAGCCA | 6193 |
| | | reverse | CCCTTATTCCATCTGCCTGAAGTA | |
| | back | forward | CCTCCCTTAAGTCTTACTCCTTCA | 6495 |
| | | reverse | CCAGGACCAATTTGACCATGATTT | |
| *AR* | front | forward | GGCCACATTCCTGACTGACA | 2695 |
| | | reverse | GGTCATCCTCCCTCCACTCT | |
| | back | forward | GGGACAACTGGTGTTCCTGT | 2586 |
| | | reverse | TCTTCCGTACGACAAAGGGC | |
| *PRDM12* | front | forward | CTCCAGACTGAGCAGGAACC | 5714 |
| | | reverse | CTTTTCGGGCGTACGATTGC | |
| | back | forward | GACACCCCACCATTAGGCTC | 6802 |
| | | reverse | GGCTGATCAACGGCATCTCT | |
| *WDR7* | front | forward | AGATTTTCATCTGTCTGACCTGCT | 5739 |
| | | reverse | AGCATCTACTCTTACCTGCCAAAA | |
| | back | forward | CTGTGTTGTGTTCAGCGCTATTAT | 6432 |
| | | reverse | GAATACAAACAGAACCACAGGAGC | |
| *NIPA1* | front | forward | GTACCAACACCAGCAATGTTCC | 7869 |
| | | reverse | TCCTTTTATGGTCAGTGACTCTGG | |
| | back | forward | TTGTGGTTCTTGGCCATACTCTTA | 5650 |
| | | reverse | TCTCACTCCGTAGCCTGG | |
| *VWA1* | front | forward | ATTCCCCTTTTCCAGGCGAG | 3040 |
| | | reverse | TCTGTCCATCCATCCCTCGT | |
| | back | forward | AGGCACGTTCTGAGAATCCC | 5317 |
| | | reverse | CCATAGGGCTCCCTGAAAGC | |
| *DMPK* | front | forward | CCCTGGTGGTGGTGTAATCC | 1411 |
| | | reverse | GCTGCACAGTCTCCACTTCT | |
| | back | forward | AGTCCTGTGGCTCTGTGTACTA | 491 |
| | | reverse | CTGGAGGGGCCACTTTAGATA | |
| *NUTM2B-AS1* | front | forward | TCCTTCAATGTAACAGAAGTAACCA | 6600 |

| | | | | |
|---|---|---|---|---|
| | | reverse | TTTGACTGGGTCCCCCACTA | |
| | back | forward | ACTACTCCCCACCTCAGTGATCC | 6032 |
| | | reverse | AACTGATCAAGCCAGAGACCAG | |
| *PABPN1* | front | forward | CCAGCCAAATGGTCAAAGCC | 6127 |
| | | reverse | ACTCCATAGCCCAACCAAGC | |
| | back | forward | GTGGCCATCCCAAAGGGTAA | 6458 |
| | | reverse | GCCGATCAGGAGTGACTGAC | |
| *CNBP* | front | forward | CAGCACCCCGTCCTTTATCC | 6892 |
| | | reverse | CGTGCCACTGGTGAATAGGG | |
| | back | forward | TTCTAGGGGGACAGGTGTCTTT | 961 |
| | | reverse | AACAATTCTTACCACACACATCAG | |
| *LRP12* | front | forward | CTCGGGCCCAAAGCTTCTTA | 6813 |
| | | reverse | GCTACGCATTGTCCTAGGCT | |
| | back | forward | CGGAGTGCCCCTCTTTCAAT | 6964 |
| | | reverse | ACTGGCCGTCTCCTACTCAT | |
| *GIPC1* | front | forward | TGACTACGGTGAATGGTGCC | 6680 |
| | | reverse | CAGCCTGAAGCTGGAAGACA | |
| | back | forward | CCTCCTTCCCGATTTTATAGCTCA | 4421 |
| | | reverse | GCAAGACAGAGAAATACCTCAGGA | |
| *PLIN4* | front | forward | GAGAAGCGACTAAAAGGCACTCT | 5928 |
| | | reverse | GGACACAAAGACAGCAAAACAATC | |
| | back | forward | GAAGCCCGGAAATTGTATTCTCAG | 6171 |
| | | reverse | GAAAAGATGGGGAAGGAGTCAGAT | |
| *ZIC2* | front | forward | GGAGACAGTTGTGGGTCCTG | 4459 |
| | | reverse | CGTTGAGCACATTCTGCGAG | |
| | back | forward | TCCCGGAGGTCACAATACCT | 2439 |
| | | reverse | TGAGGGTTTTTGCCCGATCA | |
| *XYLT1* | front | forward | GATGAGACACCCTCGTCCAC | 5800 |
| | | reverse | GACCAGACCTTTTCGCAAGC | |
| | back | forward | TGTTGCCAGAACGCTCTCTT | 4708 |
| | | reverse | GTGATTTGGGGACTACGGGG | |
| *FOXL2* | front | forward | GTTTCCTGGCCTTAGACCCC | 979 |
| | | reverse | GCCCTCGGGATTTCTCCAAA | |
| | back | forward | CAGCCCTGGTGAAGAAGAGG | 1262 |
| | | reverse | ACGTGTAGCTGGTGAGTGTG | |
| *HOXA13* | front | forward | TCTGTGTCAACTCTCCGCAC | 2535 |
| | | reverse | CCAGGCTCCCTGTGGAAAAT | |
| | back | forward | CAGGCAGCCAACAAACTGAC | 2233 |
| | | reverse | TGTTTTGTTGCGGGCGTAAG | |
| *HOXD13* | front | forward | GAGTTCCTGTGCCCTCAGAC | 1807 |
| | | reverse | TTCCAGCATAGCCTTTGGGG | |
| | back | forward | CACCATCCCCCACCTTACAC | 1428 |
| | | reverse | AGCCCAGGCATAGAGACTCA | |
| *PHOX2B* | front | forward | GGCCTTGGGGTAATCCAACA | 5798 |
| | | reverse | TAGAGAGGCCACGGTTCAGA | |

| | | | | |
|---|---|---|---|---|
| | back | forward | GTGAGGTCGATCTTCAGGGC | 3981 |
| | | reverse | CTCGACTAGGGCGTAGGGAT | |
| *TCF4* | front | forward | AAGCCAAATGGTTTATGAACAGCA | 5741 |
| | | reverse | TGGTGCACATTGATTAAAAGGACC | |
| | back | forward | ATCCCTGACTCTTAACACCAACTC | 7179 |
| | | reverse | GTACGATGACAGCACCTTGTATTG | |

**Table S6. Adaptive sampling BED file.**

| Chr | Start | End | Target gene |
|---|---|---|---|
| 12 | 6932419 | 6941271 | *ATN1* |
| 9 | 69033039 | 69041748 | *FXN* |
| 6 | 16322944 | 16332146 | *ATXN1* |
| 12 | 111590652 | 111603464 | *ATXN2* |
| 14 | 92064299 | 92075814 | *ATXN3* |
| 19 | 13201533 | 13212248 | *CACNA1A* |
| 3 | 63908291 | 63918156 | *ATXN7* |
| 13 | 70134809 | 70144568 | *ATXN8* |
| 6 | 170556802 | 170567008 | *TBP* |
| 4 | 3070150 | 3079720 | *HTT* |
| X | 147905849 | 147917374 | *FMR1* |
| 9 | 27567035 | 27579934 | *C9orf72* |
| 4 | 39343362 | 39353374 | *RFC1* |
| 22 | 45790160 | 45800365 | *ATXN10* |
| 5 | 146874004 | 146882985 | *PPP2R2B* |
| 16 | 66490940 | 66502525 | *BEAN1* |
| 20 | 2647619 | 2657303 | *NOP56* |
| 1 | 57362564 | 57371471 | *DAB1* |
| 2 | 190876543 | 190887430 | *GLS* |
| 13 | 102153291 | 102166924 | *FGF14* |
| 21 | 43771993 | 43781689 | *CSTB* |
| 16 | 72781103 | 72792516 | *ZFHX3* |
| 1 | 149386630 | 149398172 | *NOTCH2NLC* |
| 20 | 4693389 | 4704003 | *PRNP* |
| X | 25009435 | 25018182 | *ARX* |
| 8 | 118360904 | 118372462 | *SAMD12* |
| 2 | 96192357 | 96202438 | *STARD7* |
| 5 | 10351801 | 10360818 | *MARCHF6* |
| 16 | 24608987 | 24617921 | *TNRC6A* |
| 4 | 159335854 | 159347916 | *RAPGEF2* |
| 3 | 183706443 | 183719559 | *YEATS2* |
| 16 | 87600093 | 87609192 | *JPH3* |
| X | 148496407 | 148505575 | *AFF2* |
| 3 | 181708220 | 181717723 | *SOX3* |
| 12 | 50498184 | 50511349 | *DIP2B* |
| 7 | 55880761 | 55892439 | *ZNF713* |
| 11 | 119201125 | 119211006 | *CBL* |
| X | 67540174 | 67549624 | *AR* |
| 9 | 130677215 | 130687739 | *PRDM12* |
| 18 | 57019595 | 57029662 | *WDR7* |
| 15 | 22779040 | 22792265 | *NIPA1* |
| 1 | 1431673 | 1440174 | *VWA1* |

| | | | |
|---|---|---|---|
| 19 | 45765616 | 45775133 | *DMPK* |
| 10 | 79819073 | 79834400 | *NUTM2B-AS1* |
| 14 | 23316393 | 23327552 | *PABPN1* |
| 3 | 129167564 | 129177076 | *CNBP* |
| 8 | 104583324 | 104593148 | *LRP12* |
| 19 | 14491078 | 14500530 | *GIPC1* |
| 19 | 4504672 | 4519019 | *PLIN4* |
| 13 | 99980276 | 99989827 | *ZIC2* |
| 16 | 17466557 | 17475315 | *XYLT1* |
| 3 | 138941676 | 138950217 | *FOXL2* |
| 7 | 27194894 | 27204432 | *HOXA13* |
| 2 | 176088736 | 176097202 | *HOXD13* |
| 4 | 41741799 | 41750181 | *PHOX2B* |
| 18 | 55217865 | 55643677 | *TCF4* |

**Table S7. Known STR disease repeat expansion information.**

| Gene | Chr | Start | End | Known motif | Pathogenic expansion repeat number |
|---|---|---|---|---|---|
| *ATN1* | chr12 | 6936717 | 6936775 | CAG | 48 |
| *FXN* | chr9 | 69037275 | 69037314 | GAA | 66 |
| *ATXN1* | chr6 | 16327636 | 16327723 | CAG | 38 |
| *ATXN2* | chr12 | 111598950 | 111599019 | CAG | 32 |
| *ATXN3* | chr14 | 92071011 | 92071052 | CAG | 52 |
| *CACNA1A* | chr19 | 13207858 | 13207897 | CAG | 18 |
| *ATXN7* | chr3 | 63912685 | 63912716 | CAG | 33 |
| *ATXN8OS* | chr13 | 70139383 | 70139428 | CAG/TAG | 73 |
| *TBP* | chr6 | 170561907 | 170562017 | CAG | 42 |
| *HTT* | chr4 | 3074876 | 3074941 | CAG | 35 |
| *FMR1* | chrX | 147911979 | 147912111 | CGG | 54 |
| *C9orf72* | chr9 | 27573485 | 27573546 | GGGGCC | 23 |
| *RFC1* | chr4 | 39348425 | 39348483 | AAGGG/ACAGG | 399 |
| *ATXN10* | chr22 | 45795355 | 45795424 | ATTCT | 279 |
| *PPP2R2B* | chr5 | 146878729 | 146878758 | CAG | 50 |
| *BEAN1* | chr16 | 66495475 | 66495509 | TGGAA | 109 |
| *NOP56* | chr20 | 2652733 | 2652775 | GGCCTG | 649 |
| *DAB1* | chr1 | 57367044 | 57367125 | ATTTC | 30 |
| *GLS* | chr2 | 190880873 | 190880920 | GCA | 300 |
| *FGF14* | chr13 | 102160577 | 102162726 | GAA | 250 |
| *CSTB* | chr21 | 43776429 | 43776470 | CCCCGCCCCGCG | 30 |
| *ZFHX3* | chr16 | 72787693 | 72787757 | GCC | 40 |
| *NOTCH2NLC* | chr1 | 149390803 | 149390842 | CGG | 65 |
| *PRNP* | chr20 | 4699379 | 4699380 | CCTCAGGGCGGTGGTGGCTGG | 7 |
| *ARX* | chrX | 25013654 | 25013697 | GCC | 16 |
| *SAMD12* | chr8 | 118366813 | 118366918 | TTTCA | 104 |
| *STARD7* | chr2 | 96197067 | 96197124 | ATTTC | 149 |
| *MARCHF6* | chr5 | 10356339 | 10356411 | TTTCA | 699 |
| *TNRC6A* | chr16 | 24613439 | 24613532 | TTTTA | 18 |
| *RAPGEF2* | chr4 | 159342527 | 159342618 | TTTTA | 17 |
| *YEATS2* | chr3 | 183712113 | 183712552 | TTTCA | 0 |
| *JPH3* | chr16 | 87604283 | 87604329 | CTG | 39 |
| *AFF2* | chrX | 148500605 | 148500753 | CCG | 200 |
| *SOX3* | chr3 | 181712415 | 181712456 | GCG | 14 |
| *DIP2B* | chr12 | 50505001 | 50505022 | CGG | 350 |
| *ZNF713* | chr7 | 55887537 | 55887689 | CGG | 22 |
| *CBL* | chr11 | 119206289 | 119206322 | CCG | 3 |
| *AR* | chrX | 67545317 | 67545419 | CAG | 37 |
| *PRDM12* | chr9 | 130681606 | 130681641 | GCG | 17 |
| *WDR7* | chr18 | 57024413 | 57025039 | NNN | 1 |
| *NIPA1* | chr15 | 22786677 | 22786701 | GCG | 10 |

| | | | | | |
|---|---|---|---|---|---|
| *VWA1* | chr1 | 1435799 | 1435820 | GGCGCGGAGC | 2 |
| *DMPK* | chr19 | 45770205 | 45770266 | CTG | 49 |
| *NUTM2B-AS1* | chr10 | 79826364 | 79826403 | CGG | 15 |
| *PABPN1* | chr14 | 23321472 | 23321511 | GCG | 6 |
| *CNBP* | chr3 | 129172577 | 129172656 | CCTG | 49 |
| *LRP12* | chr8 | 104588965 | 104588999 | CGG | 89 |
| *GIPC1* | chr19 | 14496029 | 14496104 | CGG | 69 |
| *PLIN4* | chr19 | 4510488 | 4513581 | NNN | 31 |
| *ZIC2* | chr13 | 99985449 | 99985494 | GCG | 24 |
| *XYLT1* | chr16 | 17470869 | 17470967 | GGC | 99 |
| *FOXL2* | chr3 | 138946022 | 138946062 | GCG | 21 |
| *HOXA13* | chr7 | 27199827 | 27199967 | GCG | 23 |
| *HOXD13* | chr2 | 176093058 | 176093099 | GCG | 21 |
| *PHOX2B* | chr4 | 41745976 | 41746022 | GCG | 23 |
| *TCF4* | chr18 | 55586155 | 55586227 | TGC | 50 |

**Table S8. STRiker configuration parameters used in benchmark analysis.**

| **Program running paramter** | **Value** |
|---|---|
| MINIMUM_MOTIF_LENGTH | 3 |
| MAXIMUM_MOTIF_LENGTH | 30 |
| CONSECUTIVE_THRESHOLD | 10 |
| MINIMUM_MOTIF_COVERAGE | 5 |
| COVERAGE_THRESHOLD | 30 |

### Table S9. STRiker computational performance benchmarking.

| Sequencing date | File name | BAM line number | Real time (seconds) 1st | Real time (seconds) 2nd | Real time (seconds) 3rd | Average time (s) |
|---|---|---|---|---|---|---|
| 20240512 | 240512_LYJ-M_Cas9_multi_sorted.bam | 479526 | 14.111 | 14.409 | 14.078 | 14.199 |
| 20241119 | 241119_OMS_Cas9_nonAS_sorted.bam | 2357895 | 31.356 | 30.909 | 30.952 | 31.072 |
| 20240511 | 240511_LYJ-F_Cas9_multi_sorted.bam | 792831 | 32.006 | 32.045 | 32.208 | 32.086 |
| 20250415 | 250415_G1622_Cas9_tier-all_adaptive.bam | 1699930 | 32.908 | 32.958 | 32.7 | 32.855 |
| 20231218 | 231218_MJS_G1254_bae_sorted.bam | 1711356 | 24.027 | 23.986 | 24.006 | 24.006 |
